# Resting EEG Periodic and Aperiodic Components Predict Cognitive Decline Over 10 Years

**DOI:** 10.1101/2023.07.17.549371

**Authors:** Anna J. Finley, Douglas J. Angus, Erik Knight, Carien M. van Reekum, Margie E. Lachman, Richard J. Davidson, Stacey M. Schaefer

## Abstract

Measures of intrinsic brain function at rest show promise as predictors of cognitive decline in humans, including EEG metrics such as individual alpha peak frequency (IAPF) and the aperiodic exponent, reflecting the strongest frequency of alpha oscillations and the relative balance of excitatory:inhibitory neural activity, respectively. Both IAPF and the aperiodic exponent decrease with age and have been associated with worse executive function and working memory. However, few studies have jointly examined their associations with cognitive function, and none have examined their association with longitudinal cognitive decline rather than cross-sectional impairment. In a preregistered secondary analysis of data from the longitudinal Midlife in the United States (MIDUS) study, we tested whether IAPF and aperiodic exponent measured at rest predict cognitive function (*N* = 235; age at EEG recording *M* = 55.10, SD = 10.71) over 10 years. The IAPF and the aperiodic exponent interacted to predict decline in overall cognitive ability, even after controlling for age, sex, education, and lag between data collection timepoints. Post-hoc tests showed that “mismatched” IAPF and aperiodic exponents (e.g., higher exponent with lower IAPF) predicted greater cognitive decline compared to “matching” IAPF and aperiodic exponents (e.g., higher exponent with higher IAPF; lower IAPF with lower aperiodic exponent). These effects were largely driven by measures of executive function. Our findings provide the first evidence that IAPF and the aperiodic exponent are joint predictors of cognitive decline from midlife into old age and thus may offer a useful clinical tool for predicting cognitive risk in aging.

**Significance Statement:** Measures of intrinsic brain function at rest assessed noninvasively from the scalp using electroencephalography (EEG) show promise as predictors of cognitive decline in humans. Using data from 235 participants from the Midlife in the United States (MIDUS) longitudinal study, we found two resting EEG markers (individual peak alpha frequency and aperiodic exponent) interacted to predict cognitive decline over a span of 10 years. Follow-up analyses revealed that “mismatched” markers (i.e., high in one and low in the other) predicted greater cognitive decline compared to “matching” markers. Because of the low cost and ease of collecting EEG data at rest, the current research provides evidence for possible scalable clinical applications for identifying individuals at risk for accelerated cognitive decline.

## Introduction

Measures of spontaneous (i.e., resting-state) neural activity yield important insights into the intrinsic functioning of the brain. For example, individual alpha peak frequency (IAPF), the frequency at which power in the alpha band (i.e., 7-13 Hz) is the strongest, is negatively correlated with age (Klimesch, 1997; Clark et al., 2004; Finley et al., 2022; Merkin et al., 2023), and may reflect neuroanatomical differences and age-related changes in white matter (Babiloni et al., 2008; Valdés-Hernández et al., 2010; Kramberger et al., 2017). Across adulthood, higher IAPF is associated with better performance on multiple metrics of cognitive function, including working memory, reading comprehension, and a general intelligence factor (e.g., (Klimesch, 1997; Angelakis et al., 2004; Clark et al., 2004; Grandy et al., 2013a).

In addition to the periodic (i.e., oscillatory) activity found in canonical EEG bands, aperiodic activity is present across all frequencies. Aperiodic activity follows a 1/*f* function and can be described by the slope of the function (referred to as the exponent), and where the function crosses the y-axis (referred to as the offset; Donoghue et al., 2020). The aperiodic exponent is thought to correspond to the synchronized firing of neurons, such that flatter spectra are indicative of reduced synchronization, or greater neural noise (Voytek and Knight, 2015). Recent data suggest that the aperiodic exponent is related to the ratio of excitatory to inhibitory neural activity, such that flatter slopes (i.e., smaller exponents) relate to greater excitatory to inhibitory activity (Gao et al., 2017), while the offset is related to neural spiking rates, such that greater spiking activity is reflected in greater overall spectral power (Manning et al., 2009; Miller et al., 2012). Although research on aperiodic activity is in its infancy, work has associated aperiodic activity, particularly the aperiodic exponent, with age and cognitive functioning, such that flatter spectra are associated with older age (Voytek et al., 2015; Finley et al., 2022; Merkin et al., 2023), physiological markers of cognitive decline (Tran et al., 2020), reduced processing speed (Ouyang et al., 2020), and mediates cross-sectional associations between age and cognitive function (Voytek et al., 2015).

This study aims to answer two main research questions: 1. To what extent are individual differences in the slope of the aperiodic exponent and IAPF measured at fronto-central sites associated with cognitive function in adults, both cross-sectionally and longitudinally? 2. Is the slope of the aperiodic exponent or IAPF more strongly associated with cognitive function in adults both cross-sectionally and longitudinally?

To answer these questions, we examined the relationship between aperiodic exponent and IAPF in preregistered analyses with longitudinally assessed cognitive function in the Midlife in the United States (MIDUS) dataset. Prior work with MIDUS EEG data has found IAPF and aperiodic exponent are both negatively correlated with age, such that older individuals have lower IAPF and flatter aperiodic component slopes (Finley et al., 2022). Prior analyses of the full MIDUS2 and MIDUS3 longitudinal sample with Cognitive Project data found cross-sectional negative relationships between cognitive performance and age (n = 4,268; Lachman et al., 2014) as well as longitudinal negative relationships (n = 2,518; Hughes et al., 2018), such that older adults showed a steeper longitudinal decline. Sex was related to initial performance, such that women performed better on the episodic memory factor and men performed better on the executive functioning factor, with no influence of sex on the rate of longitudinal change (Hughes et al., 2018). The relationship between cognitive function and age was replicated in two subsamples (Hamm et al., 2020, n = 732; Knight et al., 2020, n = 843). No work to date has examined the MIDUS2 resting EEG data with any cognitive data.

## Materials and Methods

Preregistration of the following methods, hypotheses, and analyses are publicly available on OSF at https://osf.io/wyuca.

### Code Accessibility

All code used for all analyses and plots are publicly available on OSF at https://osf.io/sr4mb/. Additionally, all data are available at https://midus.wisc.edu/data/index.php. The demographic and cognitive task data are available through the MIDUS Portal or via the University of Michigan’s Inter-university Consortium of Political and Social Research (ICPSR). The EEG data are available upon request through the MIDUS Neuroimaging and Psychophysiology Repository.

### Participants

This study uses data from the MIDUS longitudinal dataset, with variables collected during the MIDUS 2 Survey, Cognitive, and Neuroscience Projects and as well as MIDUS 3 Cognitive Project. See Figure 1 for a diagram of the study design and participant flow. Education, sex, and race demographics were collected during the MIDUS 2 Survey Project, which was a prerequisite for participation in additional MIDUS 2 Projects. As depicted in Figure 1, MIDUS 2 and MIDUS 3 Cognitive Project are longitudinal and collected approximately 10 years apart (i.e., total lag; *M* = 9.71 years, *SD* = 0.92), while the MIDUS 2 Cognitive Project was completed before the MIDUS 2 Neuroscience Project (*M* = 2.06 years, *SD* = 1.26). Additional details about the study are available at http://midus.wisc.edu.

**Figure 1.**
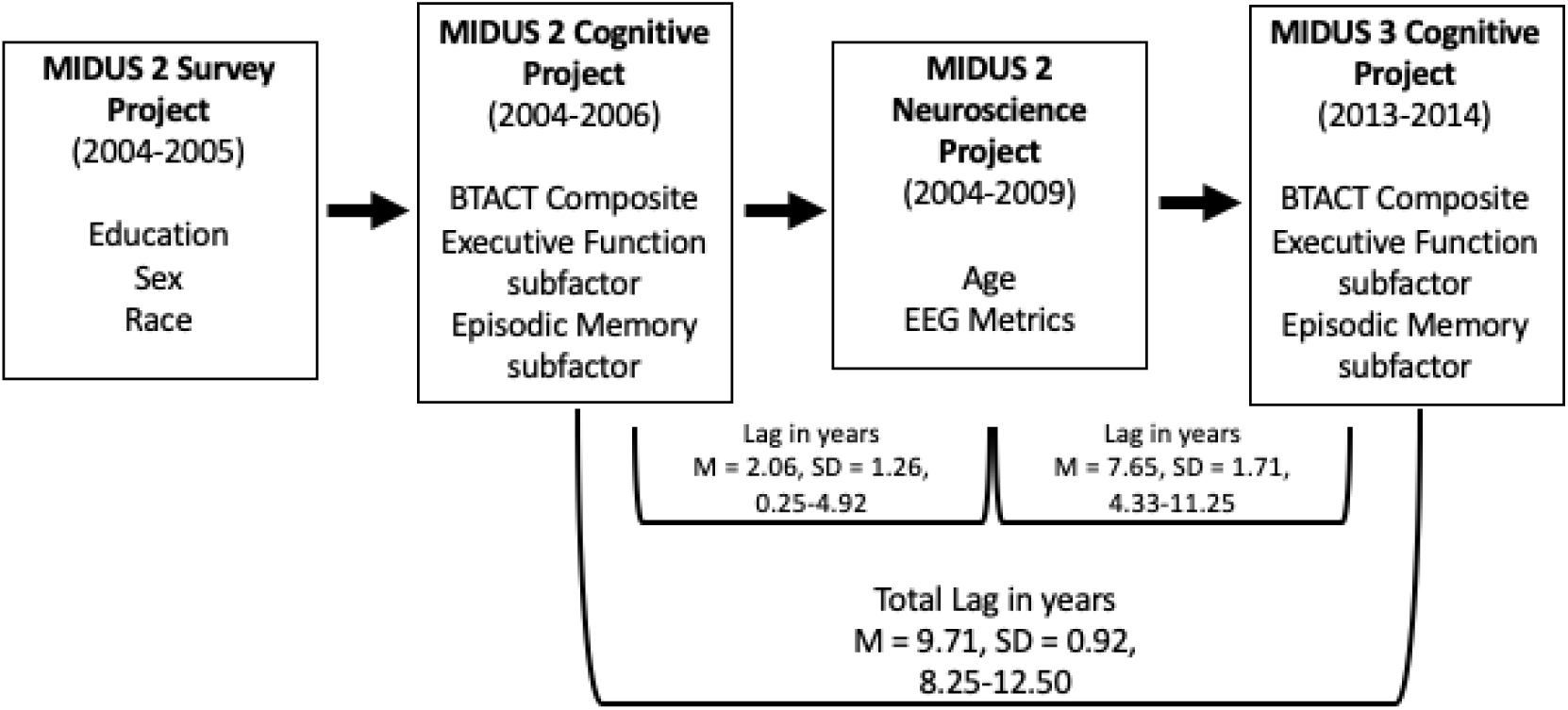
Participant flow and at which timepoint data was collected.

Based on our preregistered exclusion criteria, participants were excluded if they had poor FOOOF algorithm fit (defined as more than 3 standard deviations below the mean in R^2^ model fit for the frontal composite (*n* = 4); see section “Spectral parametrization: Fitting Oscillations and 1/*f* (FOOOF)” for more information), or had missing data from more than 50% of the frontal composite for any one EEG measure (note that no participants were excluded for this reason). Additionally, participants needed to participate in at least one wave of the Cognitive Project and have sufficient task data to compute at least one cognitive function metric. Our final sample consisted of *n* = 235 participants. See Table 1 for demographic information.

**Table 1:**
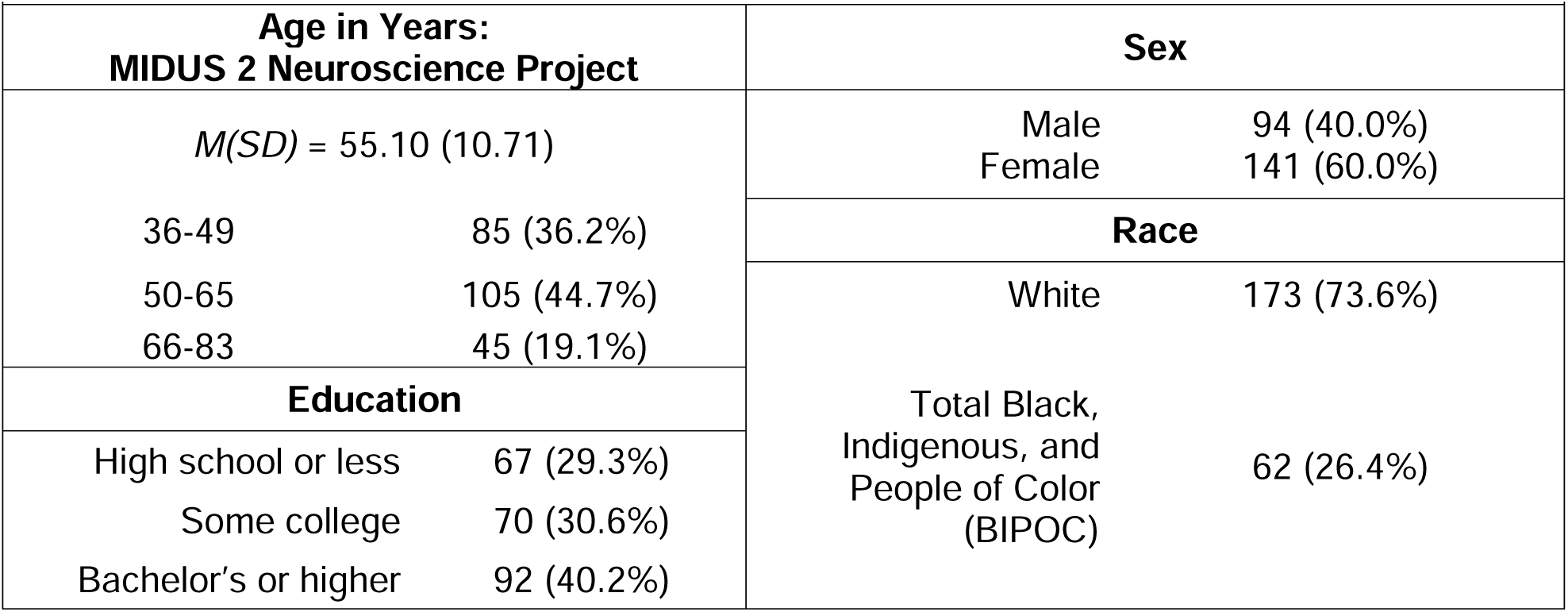
Participant Demographics, *n* = 235.

| **Table 1: Participant Demographics, $n = 235$** |  |  |  |
| --- | --- | --- | --- |
| **Age in Years:**<br>**MIDUS 2 Neuroscience Project** |  | **Sex** |  |
| $M(SD) = 55.10 (10.71)$ | | Male | 94 (40.0%) |
|  |  | Female | 141 (60.0%) |
| 36-49 | 85 (36.2%) | **Race** |  |
| 50-65 | 105 (44.7%) | White | 173 (73.6%) |
| 66-83 | 45 (19.1%) | Total Black,<br>Indigenous, and<br>People of Color<br>(BIPOC) | 62 (26.4%) |
| **Education** |  |  |  |
| High school or less | 67 (29.3%) |  |  |
| Some college | 70 (30.6%) |  |  |
| Bachelor's or higher | 92 (40.2%) |  |  |

### Brief Test of Adult Cognition by Telephone (BTACT)

During the MIDUS 2 and MIDUS 3 Cognitive Project, participants completed the Brief Test of Adult Cognition by Telephone (BTACT; Tun and Lachman, 2006; Lachman et al., 2014; Hughes et al., 2018), which includes 7 neuropsychological tasks that load onto an episodic memory factor (immediate word list recall, delayed word list recall) and executive functioning factor (backward digit span, category verbal fluency, number series, 30 seconds and counting task, stop and go switch task mixed trials), as well as an overall BTACT composite score. The BTACT has good construct validity and performs comparably to lab-based assessments (Lachman et al., 2014). Participants’ performance on the BTACT (composite and separate episodic memory and the executive functioning factors) were standardized to the MIDUS 2 sample scores (i.e., individual task measures were *z*-scored within the retained sample) and averaged across relevant tasks. BTACT scores were not computed for factors or the overall composite if participants were missing data for one or more tasks. See (Lachman et al., 2014) for additional details on the BTACT procedure.

### EEG Recording and Preprocessing

Resting-state EEG data were recorded in one-minute periods (3 minutes eyes open, 3 minutes eyes closed) using a 128-channel geodesic net of Ag/AgCl electrodes encased in saline-dampened sponges with an online vertex (Cz) reference (Electrical Geodesics, Inc [EGI], Eugene, OR). Signals were amplified and sampled at 500 Hz with an online bandpass filter (0.1 to 100 Hz, 16-bit precision). Offline EEG data were filtered with a 60 Hz notch filter, 0.5 Hz high-pass filter, bad channels identified and removed, and bad sections of data identified and removed. A 20-component PCA/ICA was used to visually identify and remove obvious blink, eye movement, and other artifactual components. Bad channels were replaced using a spherical spline interpolation. Data from the eyes open and eyes closed conditions were collapsed for all analyses^1^ using a pre-registered fronto-central composite of F3/Fz/F4 analog channels^2^. See Finley et al., 2022 for additional details.

### Spectral parametrization: Fitting Oscillations and 1/f (FOOOF)

EEG data were re-referenced to the average and Cz was imputed before the continuous resting data was epoched into 2 second segments with 50% overlap. Bad segments were rejected if there was a voltage deviation of +/- 100 μV in one or more channels. EEG spectral power was extracted using a 2 second Hamming window padded by a factor of 2 from 0 to 250 Hz in 0.25 Hz increments for all sensors, then analyzed using FOOOF 1.0.0 (Donoghue et al., 2020) to fit aperiodic and periodic components from 2 to 40 Hz (estimated without a knee, peaks limited in width from 1-6 Hz, minimum peak height of 0.05, relative peak threshold of 1.5 standard deviations, and maximum of 6 peaks). Aperiodic offset, exponent, and IAPF measures were extracted individually for the channels in the frontal F3/Fz/F4 ROI composite and then averaged. See Finley et al., 2022 for additional details.

### Experimental Design and Statistical Analysis

A summary of preregistered hypotheses and analyses are reported in Table 2^3^. Additional details are available in our preregistration (https://osf.io/wyuca). We used multilevel modeling implemented in R version 4.2.1 using the lmer() function within the lme4 package, which implements empirical Bayes slope estimation to handle missing data (Bates et al., 2015; R Core Team, 2022).

**Table 2:**
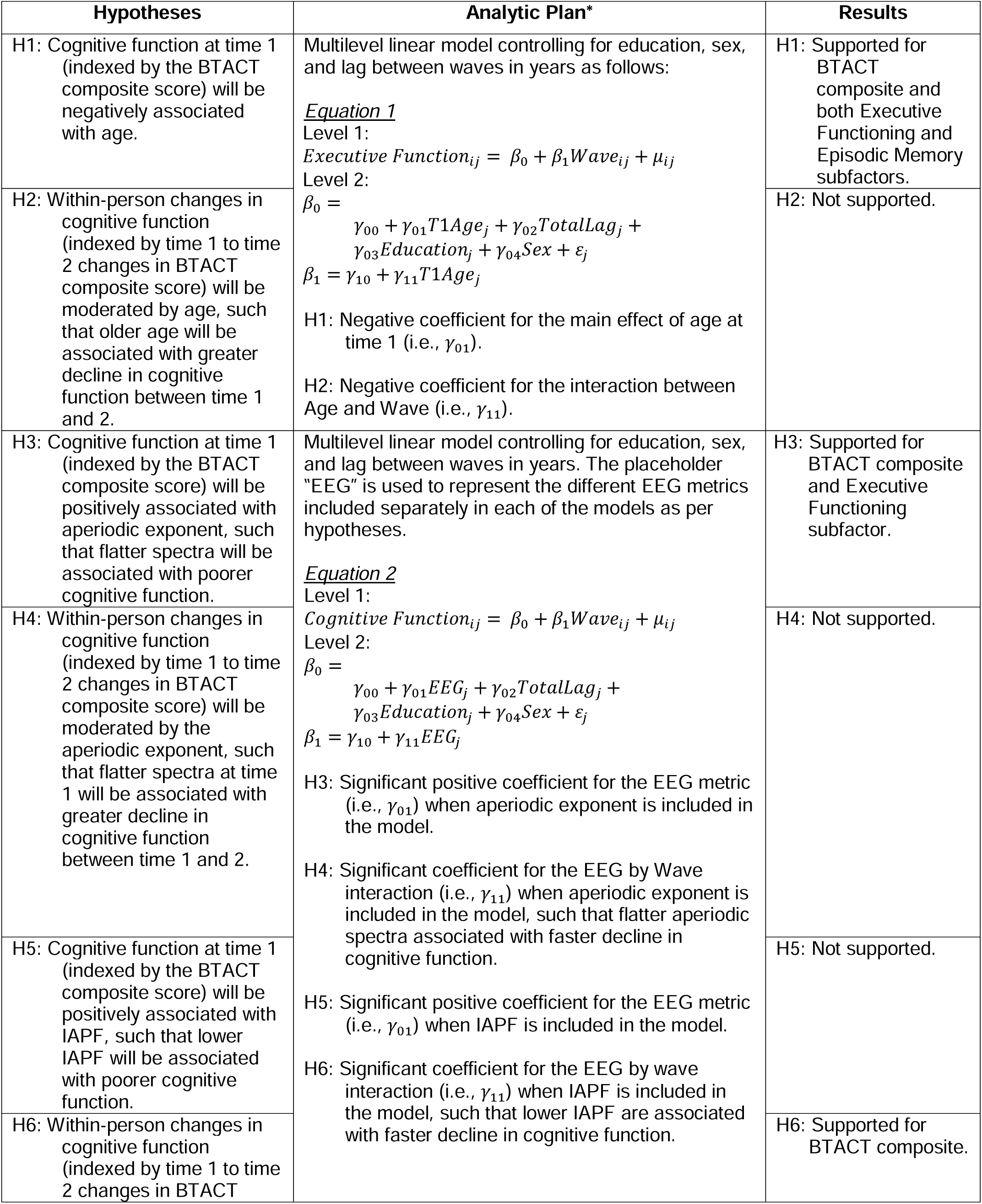
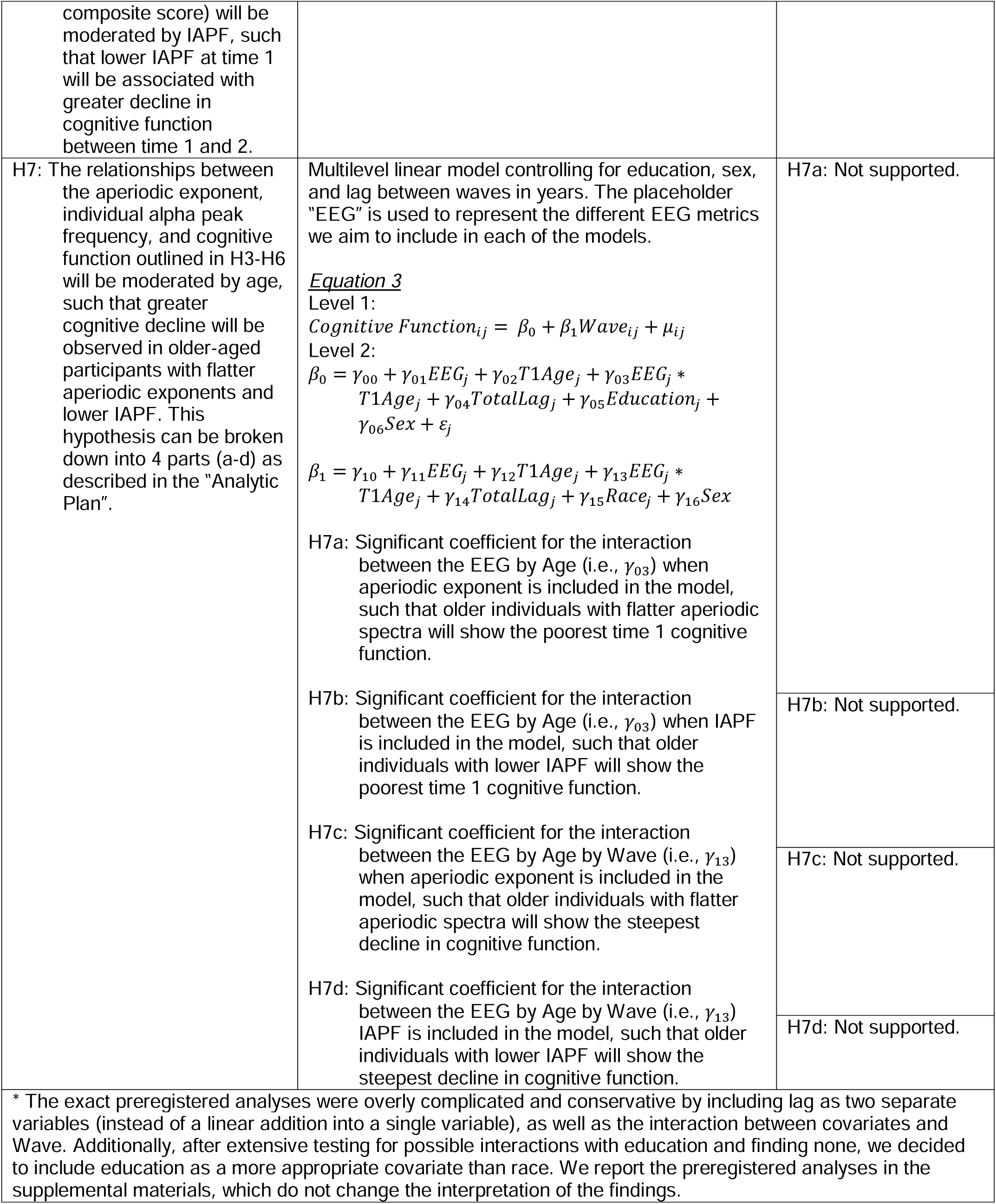
Summary of preregistered hypotheses and analyses.

**Table 3:**
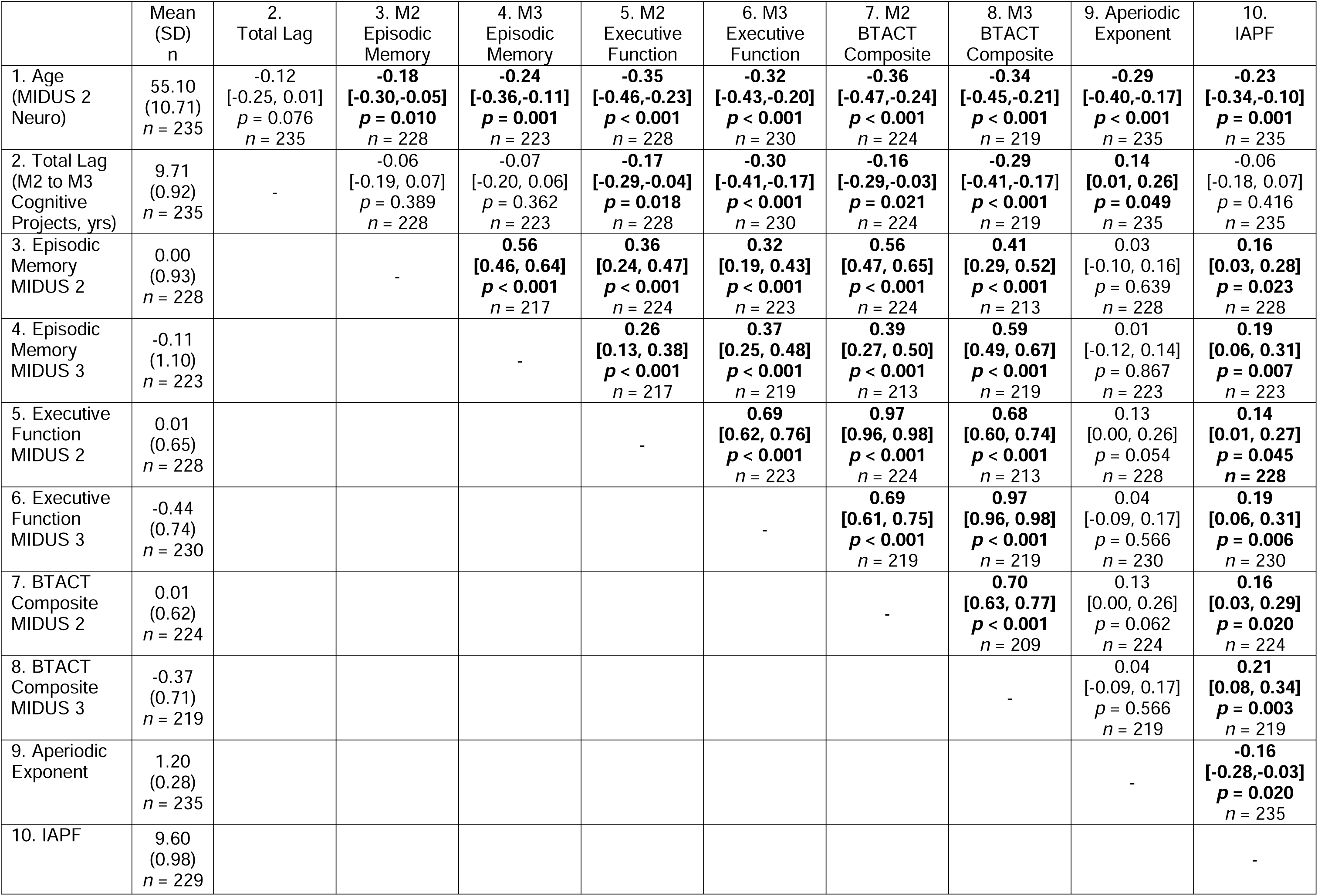
Correlations and descriptive statistics.

As reported in our preregistration (https://osf.io/wyuca), we conducted a sensitivity analysis in G*Power 3.1. Based on the most conservative estimate of complete data from our preregistration of *n* = 207, we have 95% power to detect a Pearson’s correlation of *r* = |.24|. After our final sample size was known (*n* = 235), we conducted a simulation sensitivity analysis in R based on our most complex preregistered analysis in hypothesis 7 (i.e., hypothesis 7b and 7d, the interaction between MIDUS wave, age, and EEG metric), and determined we have 80% power to detect a small effect of B = 0.15. Simulation code is available at https://osf.io/sr4mb/.

We conducted all analyses on the full BTACT composite as well as separately for the episodic memory and executive functioning factors to explore if one or both of the BTACT factors are driving effects. Parallel exploratory analyses on the associations with the aperiodic offset are described in the supplemental materials. We also explored whether the aperiodic exponent or IAPF are independently and uniquely associated with cognitive functioning, as well as if there is an interaction between the aperiodic exponent and IAPF associated with cognitive functioning, as follows:

#### Equation 4

Level 1:

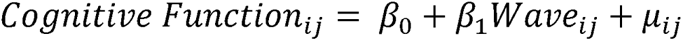

Level 2:

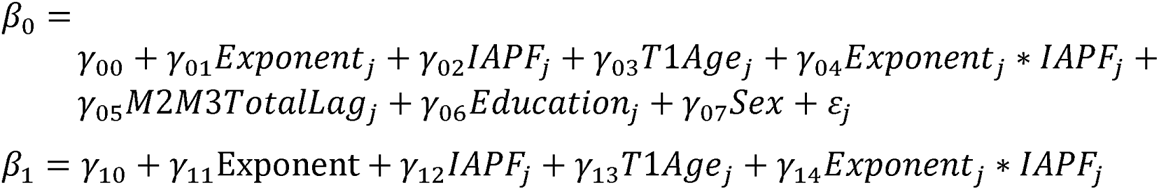

## Results

Descriptive statistics for all variables as well as zero-order correlations are presented in Table 3. Additional analyses are reported in the supplemental materials, including analyses without controlling for sex and education and analyses accounting for the presence of twins and siblings to control for genetic dependencies. None of these variations on the analyses change the interpretations of the following analyses.

### Time Effects: Hypotheses 1 and 2

To test hypothesis 1 (cognitive function at time 1 will be negatively associated with age) and hypothesis 2 (within-person changes in cognitive function will be moderated by age), we conducted a multilevel model as described in Table 2. Results are reported in Table 4. Although age was significantly related to episodic memory, executive functioning, and overall BTACT composite scores (*p*’s < .010) in support of hypothesis 1, the age-by-wave interaction was not significant for any analysis, (*p*’s > 0.096), not supporting hypothesis 2. Given our sample size with Neuroscience data (*n* = 235) is much smaller than the smallest MIDUS subsample that previously reported an age by wave interaction (e.g. n = 2,518; δ_Episodic Memory_ = −0.010, δ_Executive Function_ = −0.012, (Hughes et al., 2018), we may have been underpowered to detect this small interaction effect.

**Table 4:** Multilevel models to test hypothesis 1 and 2.

| *Predictors* | **Episodic Memory** |  |  | **Executive Functioning** |  |  | **BTACT Composite** |  |  |
| --- | --- | --- | --- | --- | --- | --- | --- | --- | --- |
|  | *Estimates* | *CI* | *p* | *Estimates* | *CI* | *p* | *Estimates* | *CI* | *p* |
| Intercept (M2) | **-0.43** | **-0.60 – -0.26** | **<0.001** | -0.07 | -0.19 – 0.05 | 0.256 | **-0.12** | **-0.23 – -0.01** | **0.028** |
| MIDUS Wave | **-0.13** | **-0.26 – -0.00** | **0.043** | **-0.43** | **-0.50 – -0.36** | **<0.001** | **-0.38** | **-0.45 – -0.31** | **<0.001** |
| Age | **-0.02** | **-0.03 – -0.01** | **0.001** | **-0.02** | **-0.03 – -0.02** | **<0.001** | **-0.02** | **-0.03 – -0.02** | **<0.001** |
| Sex | **0.74** | **0.53 – 0.94** | **<0.001** | 0.11 | -0.03 – 0.26 | 0.128 | **0.22** | **0.08 – 0.35** | **0.001** |
| Education | **0.17** | **0.04 – 0.29** | **0.008** | **0.13** | **0.05 – 0.21** | **0.001** | **0.14** | **0.06 – 0.21** | **0.001** |
| Lag between Waves | -0.10 | -0.21 – 0.01 | 0.073 | **-0.21** | **-0.29 – -0.14** | **<0.001** | **-0.20** | **-0.27 – -0.13** | **<0.001** |
| MIDUS Wave X Age | -0.01 | -0.02 – 0.00 | 0.096 | 0.00 | -0.01 – 0.01 | 0.994 | -0.00 | -0.01 – 0.01 | 0.707 |
| **Random Effects** |  |  |  |  |  |  |  |  |  |
| $\sigma^2$ | 0.46 | | | 0.16 | | | 0.14 | | |
| $\tau_{00}$ | 0.38 | M2ID | | 0.22 | M2ID | | 0.19 | M2ID | |
| ICC | 0.45 |  |  | 0.59 |  |  | 0.58 |  |  |
| N | 234 | M2ID |  | 235 | M2ID |  | 234 | M2ID |  |
| Observations | 451 |  |  | 458 |  |  | 443 |  |  |
| Marginal R<sup>2</sup> / Conditional R<sup>2</sup> | 0.195 / 0.559 |  |  | 0.292 / 0.707 |  |  | 0.311 / 0.708 |  |  |

Note: Repeated measured cognitive data nested within participant, indicated by M2ID. Sex coded as 0 = male, 1 = female. Education coded as -1 = high school or less, 0 = some college, 1 = Bachelor's degree or higher. MIDUS Wave coded as 0 = MIDUS 2, 1 = MIDUS 3. Years between Waves (i.e., between MIDUS 2 and MIDUS 3 Cognitive Projects) and Age at M2 Neuroscience Project are mean centered. Empirical Bayes slope

### Aperiodic Exponent Effects: Hypotheses 3 and 4

To test hypothesis 3 (cognitive function at time 1 will be positively associated with aperiodic exponent) and hypothesis 4 (within-person changes in cognitive function will be moderated by aperiodic exponent), we conducted a multilevel model as described in Table 2. Results are reported in Table 5. We observed a positive association between the aperiodic exponent and the overall BTACT composite score (*p* = 0.018), such that larger aperiodic exponents were associated with better cognitive function, consistent with hypothesis 3. This association appeared to be primarily driven by the Executive Function factor (*p* = 0.012), while the effect for the Episodic Memory factor was in the same direction but not-significant (*p* = 0.254). However, flatter spectra at time 1 was not associated with greater declines in cognitive function, *p*’s > 0.120, not supporting hypothesis 4.

**Table 5:** Multilevel models to test hypothesis 3 and 4.

| *Predictors* | **Episodic Memory** |  |  | **Executive Functioning** |  |  | **BTACT Composite** |  |  |
| --- | --- | --- | --- | --- | --- | --- | --- | --- | --- |
|  | *Estimates* | *CI* | *p* | *Estimates* | *CI* | *p* | *Estimates* | *CI* | *p* |
| Intercept (M2) | **-0.41** | **-0.59 – -0.23** | **<0.001** | -0.05 | -0.18 – 0.08 | 0.450 | -0.11 | -0.23 – 0.01 | 0.085 |
| MIDUS Wave | -0.13 | -0.26 – 0.00 | 0.051 | **-0.43** | **-0.51 – -0.36** | **<0.001** | **-0.38** | **-0.45 – -0.31** | **<0.001** |
| Exponent | 0.26 | -0.19 – 0.71 | 0.254 | **0.40** | **0.09 – 0.71** | **0.012** | **0.39** | **0.10 – 0.68** | **0.009** |
| Sex | **0.71** | **0.49 – 0.93** | **<0.001** | 0.08 | -0.08 – 0.24 | 0.306 | **0.19** | **0.04 – 0.34** | **0.012** |
| Education | **0.17** | **0.04 – 0.30** | **0.008** | **0.13** | **0.04 – 0.21** | **0.003** | **0.13** | **0.05 – 0.21** | **0.002** |
| Lag between Waves | -0.08 | -0.19 – 0.04 | 0.187 | **-0.19** | **-0.28 – -0.11** | **<0.001** | **-0.18** | **-0.26 – -0.10** | **<0.001** |
| Wave x Exponent | -0.02 | -0.48 – 0.44 | 0.926 | -0.21 | -0.47 – 0.05 | 0.120 | -0.16 | -0.42 – 0.09 | 0.204 |
| **Random Effects** |  |  |  |  |  |  |  |  |  |
| $\sigma^2$ | 0.47 | | | 0.15 | | | 0.14 | | |
| $\tau_{00}$ | 0.44 | M2ID | | 0.28 | M2ID | | 0.25 | M2ID | |
| ICC | 0.48 |  |  | 0.65 |  |  | 0.65 |  |  |
| N | 234 | M2ID |  | 235 | M2ID |  | 234 | M2ID |  |
| Observations | 451 |  |  | 458 |  |  | 443 |  |  |
| Marginal $R^2$ /<br>Conditional $R^2$ | 0.136 / 0.551 | | | 0.179 / 0.710 | | | 0.182 / 0.710 | | |

Note: Repeated measured cognitive data nested within participant, indicated by M2ID. Sex coded as 0 = male, 1 = female. Education coded as -1 = high school or less, 0 = some college, 1 = Bachelor's degree or higher. MIDUS Wave coded as 0 = MIDUS 2, 1 = MIDUS 3. Years between Waves (i.e., between MIDUS 2 and MIDUS 3 Cognitive Projects) and Age at M2 Neuroscience Project are mean centered. Empirical Bayes slope estimation used (Bates et al., 2015).

### Individual Alpha Peak Frequency Effects: Hypotheses 5 and 6

To test hypothesis 5 (cognitive function at time 1 will be positively associated with IAPF) and hypothesis 6 (within-person changes in cognitive function will be moderated by IAPF, such that lower IAPF at time 1 will be associated with greater decline in cognitive function), we conducted a multilevel model as described in Table 2. Results are reported in Table 6. Hypothesis 5 was not supported. The effect of IAPF on episodic memory scores, executive function, or the overall composite were not significant, *p*’s > 0.055. However, the direction of the coefficients were in the predicted direction.

**Table 6:** Multilevel models to test hypothesis 5 and 6.

| *Predictors* | **Episodic Memory** |  |  | **Executive Functioning** |  |  | **BTACT Composite** |  |  |
| --- | --- | --- | --- | --- | --- | --- | --- | --- | --- |
|  | *Estimates* | *CI* | *p* | *Estimates* | *CI* | *p* | *Estimates* | *CI* | *p* |
| Intercept (M2) | **-0.40** | **-0.57 – -0.22** | **<0.001** | -0.03 | -0.16 – 0.09 | 0.596 | -0.09 | -0.21 – 0.03 | 0.146 |
| MIDUS Wave | -0.13 | -0.25 – 0.00 | 0.052 | **-0.43** | **-0.51 – -0.36** | **<0.001** | **-0.38** | **-0.45 – -0.31** | **<0.001** |
| IAPF | 0.12 | -0.00 – 0.25 | 0.055 | 0.07 | -0.02 – 0.16 | 0.115 | 0.08 | -0.00 – 0.16 | 0.058 |
| Sex | **0.68** | **0.47 – 0.89** | **<0.001** | 0.06 | -0.10 – 0.21 | 0.469 | **0.16** | **0.02 – 0.31** | **0.030** |
| Education | **0.17** | **0.05 – 0.30** | **0.007** | **0.14** | **0.05 – 0.22** | **0.002** | **0.13** | **0.05 – 0.21** | **0.001** |
| Lag between Waves | -0.06 | -0.17 – 0.06 | 0.326 | **-0.17** | **-0.26 – -0.09** | **<0.001** | **-0.16** | **-0.24 – -0.08** | **<0.001** |
| Wave x IAPF | 0.07 | -0.06 – 0.20 | 0.281 | 0.07 | -0.00 – 0.15 | 0.061 | **0.07** | **0.00 – 0.14** | **0.047** |
| **Random Effects** |  |  |  |  |  |  |  |  |  |
| $\sigma^2$ | 0.47 | | | 0.15 | | | 0.13 | | |
| $\tau_{00}$ | 0.42<sub>M2ID</sub> | | | 0.28<sub>M2ID</sub> | | | 0.25<sub>M2ID</sub> | | |
| ICC | 0.47 |  |  | 0.64 |  |  | 0.65 |  |  |
| N | 234<sub>M2ID</sub> |  |  | 235<sub>M2ID</sub> |  |  | 234<sub>M2ID</sub> |  |  |
| Observations | 451 |  |  | 458 |  |  | 443 |  |  |
| Marginal R<sup>2</sup> / Conditional R<sup>2</sup> | 0.156 / 0.553 |  |  | 0.188 / 0.711 |  |  | 0.196 / 0.715 |  |  |

Note: Repeated measured cognitive data nested within participant, indicated by M2ID. Sex coded as 0 = male, 1 = female. Education coded as -1 = high school or less, 0 = some college, 1 = Bachelor's degree or higher. MIDUS Wave coded as 0 = MIDUS 2, 1 = MIDUS 3. Years between Waves (i.e., between MIDUS 2 and MIDUS 3 Cognitive Projects) and Age at M2 Neuroscience Project are mean centered. Empirical Bayes slope estimation used (Bates et al., 2015).

Hypothesis 6 was supported and in the predicted direction (*p* = 0.047), such that lower IAPF at time 1 were associated with greater declines in cognitive function as depicted in Figure 2.

**Figure 2:**
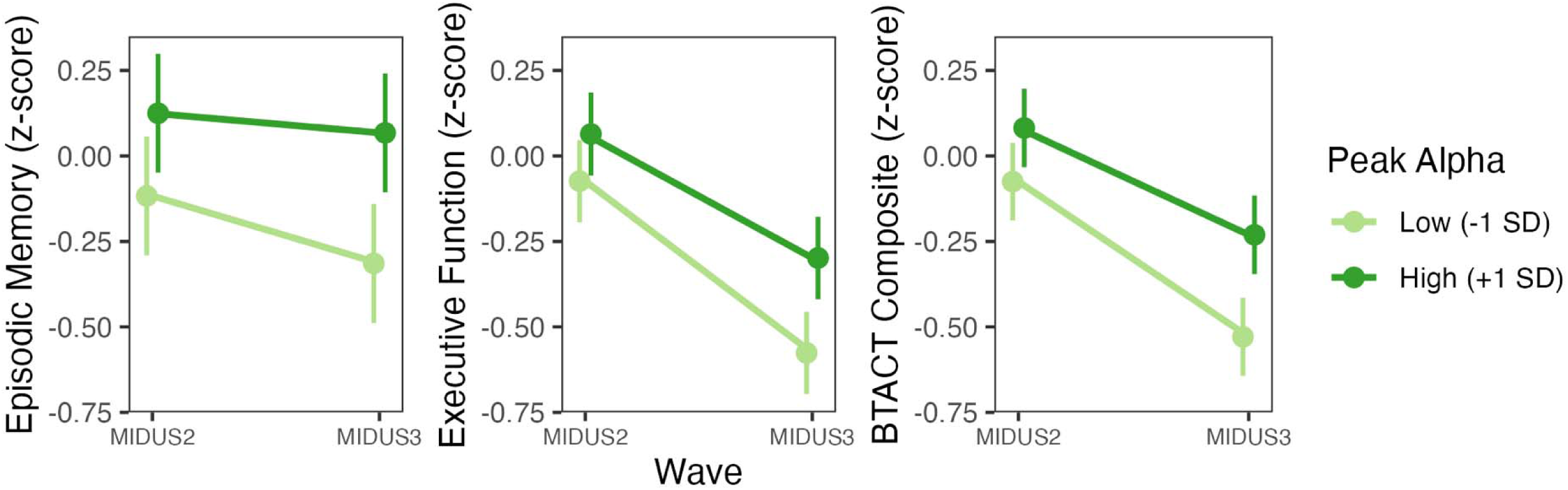
Wave by Individual Peak Alpha Frequency Interaction Plot. Plot depicting the two-way interaction wave X individual peak alpha frequency reported in Table 6 with 95% confidence interval error bars. Time 1 cognition assessed at MIDUS2 Cognitive Project, and time 2 cognition was assessed at the MIDUS 3 Cognitive Project.

### Moderation of EEG Metrics by Age: Hypothesis 7

To test hypothesis 7a (older individuals with lower aperiodic exponents will show the poorest time 1 cognitive function) and hypothesis 7c (older individuals with lower aperiodic exponents will show the steepest decline in cognitive function), we conducted a multilevel model as described in Table 2. Results are reported in Table 7. Hypothesis 7a was not confirmed as the interaction between aperiodic exponent and age was non-significant for all BTACT scores, *p*’s > 0.632. Hypothesis 7c was not supported as the wave by aperiodic exponent by age interaction was non-significant, *p*’s > 0.201.

**Table 7:** Multilevel models to test hypothesis 7a and 7c.

| *Predictors* | **Episodic Memory** |  |  | **Executive Functioning** |  |  | **BTACT Composite** |  |  |
| --- | --- | --- | --- | --- | --- | --- | --- | --- | --- |
|  | *Estimates* | *CI* | *p* | *Estimates* | *CI* | *p* | *Estimates* | *CI* | *p* |
| Intercept (M2) | -0.44 | -0.62 – -0.26 | <0.001 | -0.06 | -0.18 – 0.06 | 0.340 | -0.12 | -0.23 – -0.00 | 0.046 |
| MIDUS Wave | -0.11 | -0.24 – 0.03 | 0.122 | -0.43 | -0.50 – -0.35 | <0.001 | -0.38 | -0.46 – -0.31 | <0.001 |
| Exponent | 0.07 | -0.38 – 0.52 | 0.748 | 0.16 | -0.14 – 0.46 | 0.304 | 0.15 | -0.13 – 0.43 | 0.290 |
| Age | -0.02 | -0.03 – -0.01 | 0.002 | -0.02 | -0.03 – -0.01 | <0.001 | -0.02 | -0.03 – -0.01 | <0.001 |
| Sex | 0.73 | 0.52 – 0.94 | <0.001 | 0.11 | -0.04 – 0.26 | 0.140 | 0.22 | 0.08 – 0.36 | 0.002 |
| Education | 0.16 | 0.04 – 0.29 | 0.010 | 0.13 | 0.05 – 0.21 | 0.002 | 0.13 | 0.06 – 0.21 | 0.001 |
| Lag between Waves | -0.10 | -0.21 – 0.01 | 0.080 | -0.22 | -0.29 – -0.14 | <0.001 | -0.20 | -0.27 – -0.13 | <0.001 |
| Wave x Exponent | -0.14 | -0.62 – 0.33 | 0.557 | -0.23 | -0.50 – 0.05 | 0.104 | -0.19 | -0.45 – 0.08 | 0.166 |
| Wave X Age | -0.01 | -0.02 – 0.00 | 0.122 | -0.00 | -0.01 – 0.01 | 0.687 | -0.00 | -0.01 – 0.00 | 0.476 |
| Exponent X Age | -0.01 | -0.06 – 0.04 | 0.677 | 0.01 | -0.02 – 0.04 | 0.632 | 0.01 | -0.02 – 0.03 | 0.695 |
| Wave X Exponent X Age | 0.03 | -0.02 – 0.08 | 0.201 | 0.00 | -0.02 – 0.03 | 0.755 | 0.00 | -0.02 – 0.03 | 0.858 |
| **Random Effects** |  |  |  |  |  |  |  |  |  |
| $\sigma^2$ | 0.46 | | | 0.16 | | | 0.14 | | |
| $\tau_{00}$ | 0.39 M2ID | | | 0.22 M2ID | | | 0.19 M2ID | | |
| ICC | 0.46 |  |  | 0.59 |  |  | 0.58 |  |  |
| N | 234 M2ID |  |  | 235 M2ID |  |  | 234 M2ID |  |  |
| Observations | 451 |  |  | 458 |  |  | 443 |  |  |
| Marginal R<sup>2</sup> /<br>Conditional R<sup>2</sup> | 0.195 / 0.563 |  |  | 0.294 / 0.710 |  |  | 0.311 / 0.710 |  |  |

Note: Repeated measured cognitive data nested within participant, indicated by M2ID. Sex coded as 0 = male, 1 = female. Education coded as -1 = high school or less, 0 = some college, 1 = Bachelor's degree or higher. MIDUS Wave coded as 0 = MIDUS 2, 1 = MIDUS 3. Years between Waves (i.e., between MIDUS 2 and MIDUS 3 Cognitive Projects) and Age at M2 Neuroscience Project are mean centered. Empirical Bayes slope estimation used (Bates et al., 2015).

To test hypothesis 7b (older individuals with lower IAPF will show the poorest time 1 cognitive function) and hypothesis 7d (older individuals with lower IAPF will show the steepest decline in cognitive function), we conducted a multilevel model as described in Table 2. Results are reported in Table 8. Hypothesis 7b was not confirmed as the interaction between IAPF and age was non-significant for all BTACT scores, *p*’s > 0.374. Hypothesis 7c was not significant as the wave by IAPF by age interaction was non-significant, *p*’s > 0.301.

**Table 8:** Multilevel models to test hypothesis 7b and 7d.

| *Predictors* | **Episodic Memory** |  |  | **Executive Functioning** |  |  | **BTACT** |  |  |
| --- | --- | --- | --- | --- | --- | --- | --- | --- | --- |
|  | *Estimates* | *CI* | *p* | *Estimates* | *CI* | *p* | *Estimates* | *CI* | *p* |
| Intercept (M2) | **-0.42** | **-0.59 – -0.24** | **<0.001** | -0.06 | -0.18 – 0.06 | 0.360 | -0.11 | -0.22 – 0.00 | 0.052 |
| MIDUS Wave | -0.13 | -0.26 – 0.00 | 0.057 | **-0.44** | **-0.52 – -0.36** | **<0.001** | **-0.39** | **-0.46 – -0.32** | **<0.001** |
| IAPF | 0.08 | -0.05 – 0.21 | 0.225 | 0.01 | -0.07 – 0.10 | 0.797 | 0.02 | -0.06 – 0.10 | 0.610 |
| Age | **-0.02** | **-0.03 – -0.01** | **0.005** | **-0.02** | **-0.03 – -0.02** | **<0.001** | **-0.02** | **-0.03 – -0.02** | **<0.001** |
| Sex | **0.72** | **0.51 – 0.93** | **<0.001** | 0.11 | -0.04 – 0.25 | 0.145 | **0.21** | **0.08 – 0.35** | **0.002** |
| Education | **0.17** | **0.04 – 0.29** | **0.009** | **0.14** | **0.06 – 0.22** | **0.001** | **0.14** | **0.06 – 0.21** | **<0.001** |
| Lag between Waves | -0.09 | -0.20 – 0.02 | 0.116 | **-0.21** | **-0.29 – -0.13** | **<0.001** | **-0.19** | **-0.26 – -0.12** | **<0.001** |
| Wave x IAPF | 0.05 | -0.09 – 0.18 | 0.483 | 0.07 | -0.01 – 0.15 | 0.071 | 0.07 | -0.01 – 0.14 | 0.076 |
| Wave X Age | -0.01 | -0.02 – 0.00 | 0.165 | 0.00 | -0.01 – 0.01 | 0.741 | -0.00 | -0.01 – 0.01 | 0.994 |
| IAPF X Age | 0.00 | -0.01 – 0.01 | 0.836 | 0.00 | -0.00 – 0.01 | 0.350 | 0.00 | -0.00 – 0.01 | 0.374 |
| Wave X IAPF X Age | 0.00 | -0.01 – 0.01 | 0.896 | -0.00 | -0.01 – 0.00 | 0.301 | -0.00 | -0.01 – 0.00 | 0.379 |
| **Random Effects** |  |  |  |  |  |  |  |  |  |
| $\sigma^2$ | 0.47 | | | 0.15 | | | 0.14 | | |
| $\tau_{00}$ | 0.38 | M2ID | | 0.22 | M2ID | | 0.19 | M2ID | |
| ICC | 0.45 |  |  | 0.59 |  |  | 0.58 |  |  |
| N | 234 | M2ID |  | 235 | M2ID |  | 234 | M2ID |  |
| Observations | 451 |  |  | 458 |  |  | 443 |  |  |
| Marginal $R^2$ / Conditional $R^2$ | 0.203 / 0.559 | | | 0.298 / 0.713 | | | 0.318 / 0.715 | | |

Note: Repeated measured cognitive data nested within participant, indicated by M2ID. Sex coded as 0 = male, 1 = female. Education coded as -1 = high school or less, 0 = some college, 1 = Bachelor's degree or higher. MIDUS Wave coded as 0 = MIDUS 2, 1 = MIDUS 3. Years between Waves (i.e., between MIDUS 2 and MIDUS 3 Cognitive Projects) and Age at M2 Neuroscience Project are mean centered. Empirical Bayes slope estimation used (Bates et al., 2015).

### Combined Effects of Aperiodic Exponent and Individual Peak Alpha Frequency: Exploratory Analysis

We also explored whether the aperiodic exponent or IAPF are independently and uniquely associated with cognitive functioning, as well as if there an interaction between the aperiodic exponent and IAPF associated with cognitive functioning. Results of these analyses are in Table 9. There was a significant Wave by Aperiodic Exponent by IAPF interaction on the overall BTACT composite (*p* = 0.010), which was driven primarily by the Executive Functioning Factor (*p* = 0.013). These interactions are plotted in Figure 3 with 95% confidence bands.

**Figure 3:**
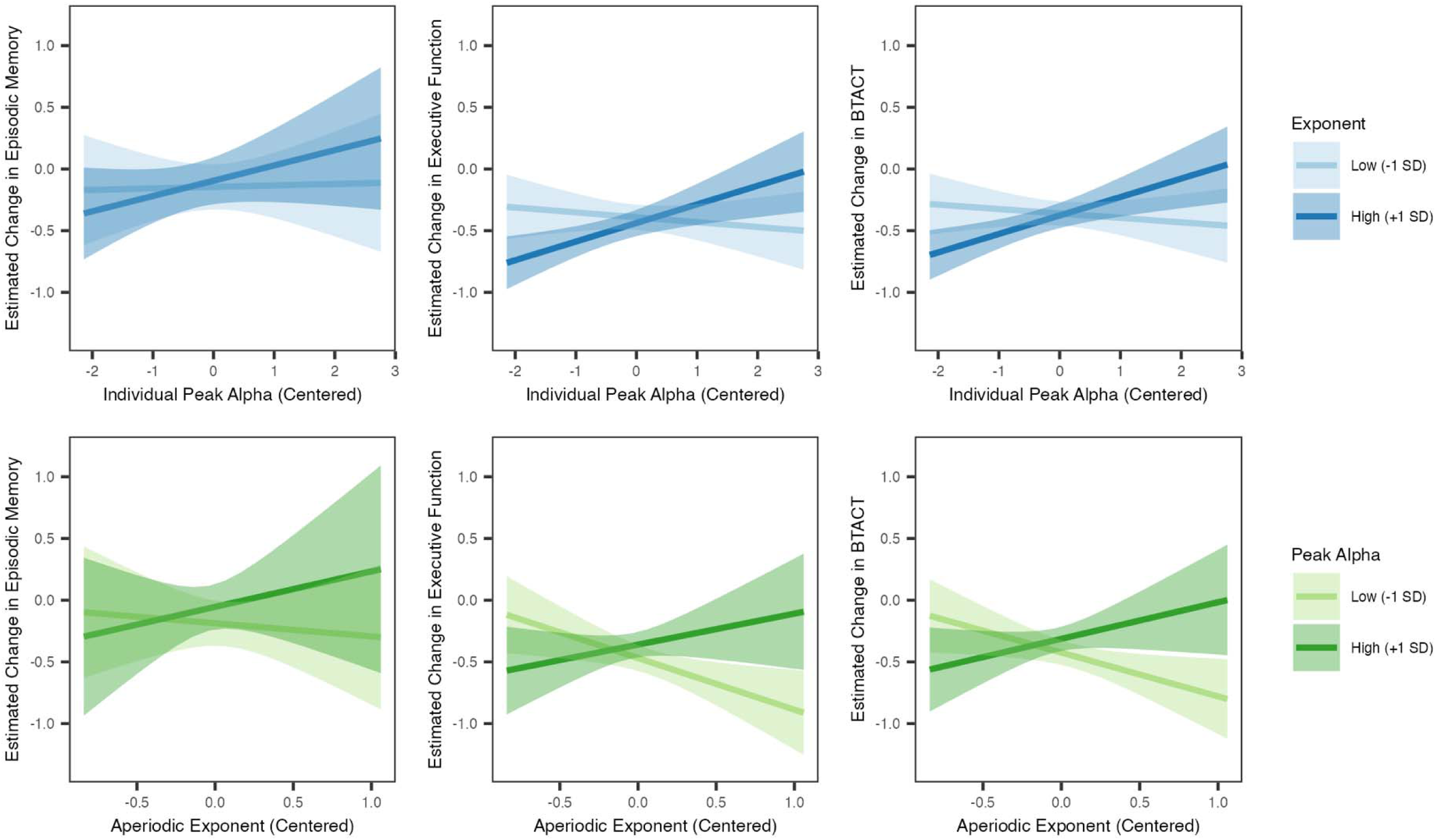
Wave by Aperiodic Exponent by Individual Peak Alpha Frequency Interaction Plot. Plot depicting the three-way interaction wave X aperiodic exponent X individual peak alpha frequency reported in Table 9, with wave depicted as the estimated change in cognitive function between the M2 and M3 Cognitive Pprojects.

**Table 9:** Multilevel models examine the interaction between aperiodic exponent and individual peak alpha frequency.

| *Predictors* | **Episodic Memory** |  |  | **Executive Functioning** |  |  | **BTACT** |  |  |
| --- | --- | --- | --- | --- | --- | --- | --- | --- | --- |
|  | *Estimates* | *CI* | *p* | *Estimates* | *CI* | *p* | *Estimates* | *CI* | *p* |
| Intercept (M2) | **-0.43** | **-0.60 – -0.25** | **<0.001** | -0.07 | -0.19 – 0.05 | 0.228 | **-0.13** | **-0.24 – -0.02** | **0.024** |
| MIDUS Wave | -0.12 | -0.25 – 0.01 | 0.069 | **-0.41** | **-0.49 – -0.34** | **<0.001** | **-0.37** | **-0.44 – -0.30** | **<0.001** |
| Exponent | 0.06 | -0.41 – 0.53 | 0.802 | 0.13 | -0.19 – 0.45 | 0.417 | 0.13 | -0.17 – 0.42 | 0.393 |
| IAPF | 0.07 | -0.06 – 0.20 | 0.269 | 0.02 | -0.07 – 0.11 | 0.634 | 0.03 | -0.05 – 0.11 | 0.482 |
| Age | **-0.02** | **-0.03 – -0.01** | **<0.001** | **-0.02** | **-0.03 – -0.02** | **<0.001** | **-0.02** | **-0.03 – -0.02** | **<0.001** |
| Sex | **0.73** | **0.52 – 0.94** | **<0.001** | 0.11 | -0.03 – 0.26 | 0.135 | **0.22** | **0.08 – 0.35** | **0.002** |
| Education | **0.17** | **0.05 – 0.30** | **0.007** | **0.14** | **0.06 – 0.22** | **0.001** | **0.14** | **0.07 – 0.22** | **<0.001** |
| Lag between Waves | -0.09 | -0.21 – 0.02 | 0.116 | **-0.21** | **-0.29 – -0.13** | **<0.001** | **-0.20** | **-0.27 – -0.12** | **<0.001** |
| Wave X Exponent | 0.09 | -0.39 – 0.58 | 0.714 | -0.08 | -0.36 – 0.19 | 0.546 | -0.03 | -0.29 – 0.23 | 0.826 |
| Wave X IAPF | 0.07 | -0.06 – 0.20 | 0.314 | 0.06 | -0.02 – 0.13 | 0.146 | 0.06 | -0.01 – 0.13 | 0.118 |
| Exponent X IAPF | -0.07 | -0.51 – 0.38 | 0.773 | -0.15 | -0.44 – 0.15 | 0.327 | -0.14 | -0.42 – 0.13 | 0.316 |
| Wave X Exponent X IAPF | **0.20** | **-0.26 – 0.67** | **0.397** | **0.34** | **0.07 – 0.61** | **0.013** | **0.33** | **0.08 – 0.59** | **0.010** |
| **Random Effects** |  |  |  |  |  |  |  |  |  |
| $\sigma^2$ | 0.47 | | | 0.15 | | | 0.13 | | |
| $\tau_{00}$ | 0.38 | M2ID | | 0.23 | M2ID | | 0.19 | M2ID | |
| ICC | 0.44 |  |  | 0.60 |  |  | 0.59 |  |  |
| N | 234 | M2ID |  | 235 | M2ID |  | 234 | M2ID |  |
| Observations | 451 |  |  | 458 |  |  | 443 |  |  |
| Marginal R<sup>2</sup> / Conditional R<sup>2</sup> | 0.202 / 0.557 |  |  | 0.302 / 0.722 |  |  | 0.323 / 0.724 |  |  |

Note: Repeated measured cognitive data nested within participant, indicated by M2ID. Sex coded as 0 = male, 1 = female. Education coded as -1 = high school or less, 0 = some college, 1 = Bachelor's degree or higher. MIDUS Wave coded as 0 = MIDUS 2, 1 = MIDUS 3. Years between Waves (i.e., between MIDUS 2 and MIDUS 3 Cognitive Projects) and Age at M2 Neuroscience Project are mean centered. Empirical Bayes slope estimation used (Bates et al., 2015).

As shown in Table 10, we examined the 3-way interaction by calculating the slope of the change in cognitive function over waves by each EEG metric while holding the other EEG metric constant at a low or high level by centering each EEG metric separately at low (−1 SD below the mean) and high (+1 above the mean). This is computationally equivalent to simple slopes analyses in regression (Aiken and West, 1991) at the second level of the multilevel model, and represents the slopes of the lines in Figure 2. More specifically, after centering one EEG metric at the low or high level, we examined the *γ*_14_ term from Equation 4. These analyses suggest that for individuals who have higher aperiodic exponents, having higher IAPF is associated with less decline in the BTACT overall composite (*b* = 0.15, *p* = 0.002) driven primarily by the executive function factor (*b* = 0.15, *p* = 0.004), whereas there was no significant relationship between IAPF and cognitive decline for individuals with low aperiodic exponents. For individuals with low IAPF, having a steeper aperiodic exponent is associated with faster cognitive decline for the overall BTACT composite (*b* = −0.36, *p* = 0.025) driven primarily by the executive function factor (*b* = −0.42, *p* = 0.013), whereas there was no significant relationship between aperiodic exponent and cognitive decline for individuals with high IAPF. Put another way, this suggests that individuals with “mismatched” IAPF and aperiodic exponents (e.g., higher exponent with lower IAPF) tend to experience faster rates of cognitive decline over a 10-year period compared to individuals with “matching” IAPF and aperiodic exponents (e.g., higher exponent with higher IAPF; lower IAPF with lower aperiodic exponent). As shown in Figure 3, the pattern of association is similar in direction for episodic memory, although the interaction fails to reach significance. This may be because there was substantially less decline in episodic memory performance (*M* = −0.11) than in executive function in performance (*M* = −0.44) in standardized units, limiting our power to detect an effect.

**Table 10:** Examining the Wave X Aperiodic Exponent X Individual Peak Alpha Frequency interaction through the slope of the change in cognitive function over waves for each BTACT measure at high and low levels of each EEG metric.

| **Change in BTACT Episodic Memory Factor from M2 to M3** |  |  |  |
| --- | --- | --- | --- |
|  | *Slope of IAPF* | *95% CI* | *p-value* |
| Low Exponent (-1 SD) | 0.01 | -0.18 – 0.20 | 0.903 |
| High Exponent (+1 SD) | 0.12 | -0.05 – 0.30 | 0.173 |
|  | *Slope of Exponent* | *95% CI* | *p-value* |
| Low IAPF (-1 SD) | -0.11 | -0.67 – 0.45 | 0.709 |
| High IAPF (+1 SD) | 0.29 | -0.47 – 1.05 | 0.457 |
| **Change in BTACT Executive Functioning Factor from M2 to M3** |  |  |  |
|  | *Slope of IAPF* | *95% CI* | *p-value* |
| Low Exponent (-1 SD) | -0.04 | -0.15 – 0.07 | 0.482 |
| High Exponent (+1 SD) | **0.15** | **0.05 – 0.25** | **0.004** |
|  | *Slope of Exponent* | *95% CI* | *p-value* |
| Low IAPF (-1 SD) | **-0.42** | **-0.75 – -0.09** | **0.013** |
| High IAPF (+1 SD) | 0.25 | -0.17 – 0.67 | 0.244 |
| **Change in BTACT Overall Composite from M2 to M3** |  |  |  |
|  | *Slope of IAPF* | *95% CI* | *p-value* |
| Low Exponent (-1 SD) | -0.04 | -0.14 – 0.07 | 0.502 |
| High Exponent (+1 SD) | **0.15** | **0.05 – 0.25** | **0.002** |
|  | *Slope of Exponent* | *95% CI* | *p-value* |
| Low IAPF (-1 SD) | **-0.36** | **-0.67 – -0.04** | **0.025** |
| High IAPF (+1 SD) | 0.30 | -0.11 – 0.70 | 0.149 |

## Discussion

In the current study, we investigated the role of periodic and aperiodic neural activity at rest measured from fronto-central sites in predicting cognitive decline in midlife and old age community dwelling adults. Due to their strong associations with age and cognitive impairment, we focused on the individual peak alpha frequency, or the frequency at which alpha oscillations peak (i.e., IAPF), and the slope of 1/*f*-like non-oscillatory (i.e. the aperiodic exponent) activity computed from a composite of frontal sites. Using a sample across the adult lifespan (age range 36-83 at time of EEG assessment), we showed that the fronto-central aperiodic exponent was related to cognitive function, such that flatter aperiodic exponents were related to worse cognitive function overall (e.g., hypothesis 3, Table 5). Additionally, IAPF was predictive of cognitive decline over approximately 10 years, such that lower IAPF was associated with more cognitive decline (e.g., hypothesis 6, Table 6). However, our exploratory analyses demonstrated that the relationships between aperiodic exponent, IAPF, and cognitive decline was moderated by the interaction between the fronto-central IAPF and fronto-central aperiodic exponent: decline was more severe in participants with “mismatched” measures (e.g., higher exponent with lower IAPF) compared to participants with “matching” measures (e.g., higher exponent with higher IAPF; lower IAPF with lower aperiodic exponent). Importantly, our results provide support for recent work and theoretical models that have linked both IAPF and the aperiodic exponent to individual differences in cognitive function and provide the first evidence that these measures of intrinsic brain function interact to predict cognitive decline and not just impairment.

The declines in cognitive function associated with the IAPF and aperiodic exponent were largely driven by the executive function component of the BTACT. This may be due to the relatively modest decline in the episodic memory component resulting in a floor effect due to relatively restricted range of episodic memory decline. Alternatively, it may be that our choice of fronto-central sites is uniquely sensitive to changes in executive functioning as they are closer to prefrontal cortex regions. Additional research in samples with larger declines in episodic memory are needed to begin to tease apart these possibilities.

In previous studies, age-related slowing of IAPF (e.g., (Grandy et al., 2013b; Scally et al., 2018), and slower IAPF in general, have been consistently associated with reduced processing speed, poorer working memory, and reduced cognitive capacity (Grandy et al., 2013a). The age-related slowing of alpha has been linked to alterations in inhibitory neural processes (e.g., the timing of neural inhibition), with the slowing observed in older adults attributed to an array of CNS pathology (e.g., vascular changes, white-matter lesions), as well as linked to mild and severe cognitive impairment (Babiloni et al., 2008; Kramberger et al., 2017). The frequency of alpha oscillations is also instrumental in the ‘gating’ of stimuli, with relatively slower IAPF being observed in individuals who struggle to rapidly adjust their attention to novel or task-relevant stimuli (Ramsay et al., 2021). However, previous work has almost exclusively focused on variations in the speed of oscillatory activity. While there was a significant IAPF by wave interaction, such that individuals with higher IAPF showed less cognitive decline, it was moderated by the higher-order aperiodic exponent by IAPF by wave interaction. This three-way interaction suggests that considering IAPF alone provides an incomplete understanding of neural activity and cognitive decline, and that consideration of non-oscillatory, aperiodic activity is also necessary.

Current models of the aperiodic exponent propose that individual differences – and state differences – in the aperiodic exponent <u>size</u> reflect excitatory:inhibitory balance (Gao et al., 2017; Waschke et al., 2021). Within this framework, relatively flatter slopes (i.e., smaller exponents) are associated with poorer cognitive performance due to the propagation of relatively dysregulated excitatory activity, which manifests in ‘noisier’, less efficient processing (Voytek et al., 2015; Dave et al., 2018; Pertermann et al., 2019). Our findings are broadly consistent with this perspective, with flatter exponents predicting overall reduced executive function and BTACT scores, but highlight the need to consider periodic oscillatory activity in conjunction with aperiodic metrics.

Simultaneous EEG/fMRI eyes-open resting recordings have found that the aperiodic exponent is related to increased BOLD signal in the auditory-salience-cerebellar network (including components of the salience network), and decreased BOLD signal in prefrontal networks, suggesting that steeper aperiodic exponents may be associated with increased arousal and/or increased attention to external stimuli (Jacob et al., 2021). It may be the case that individuals with “mismatched” aperiodic exponent and IAPF reflect a suboptimal balance between arousal and attention to external stimuli (indexed by the aperiodic exponent) with the ability to flexibly gate external stimuli (indexed by the IAPF) to perform complex cognitive tasks. Future research should attend to this possibility and examine if there are differences in the neurobiological mechanisms underlying increased rates of decline between individuals with low aperiodic exponents plus high IAPF vs. individuals with high aperiodic exponents plus low IAPF, and if these differences may signal different underlying pathologies or vulnerabilities.

While our work focused on periodic and aperiodic measures at rest, recent work suggests that the aperiodic exponent may change in response to a stimulus itself, consistent with an increase in inhibitory activity with an increase in attentional demand, independent from ERPs elicited by the stimulus. This suggests flexible shifts in the aperiodic exponent in response to task demands may be important for attention and cognitive function (Gyurkovics et al., 2022). Future work would benefit from exploration of whether flexible adjustments in aperiodic activity during tasks are integral to long term cognitive function and decline, and what if any role changes in IAPF during a task may play in moderating these effects.

Given research into the aperiodic exponent is in its infancy, it is unclear exactly why a high exponent paired with a low IAPF would be associated with increased rates of cognitive decline. It may be that the optimal excitatory:inhibitory balance reflected in the aperiodic exponent is not uniformly consistent across participants, but may vary with IAPF, such that higher aperiodic exponents may not always be better. Alternatively, excitatory:inhibitory balance can be shifted in complex ways between and across neural circuits, and the same endpoint may be achieved from reduction in excitatory activity or an increase in inhibitory activity, or some combination of both (Sohal and Rubenstein, 2019). It is possible that age-related slowing of IAPF may be associated with specific patterns of changes in inhibitory and or excitatory activity, such that lower IAPF associated with higher aperiodic exponents may reflect a suboptimal shift in activity. Future research would benefit from examining IAPF and aperiodic exponent in normally and pathologically aging participants to begin to tease apart these potential explanations and to determine when – or if – these shifts reflect pathological aging. Future work should also focus on better understanding what is causing age-related shifts in IAPF and how this may impact excitatory:inhibitory balance.

Overall, our findings challenge a simplistic view of the neurobehavioral and neuropsychological consequences of varied aperiodic and periodic activity. On one hand, gradual flattening is typically associated with poorer performance – potentially reflecting an excess of excitatory to inhibitory activity, resulting in elevated noise. However, many diseases, such as Parkinson’s Disease, are characterized by an excess of inhibitory activity, and previous studies have emphasized that excessive inhibitory activity reduces behavioral flexibility (Song et al., 2021; Vinding et al., 2022; McKeown et al., 2023). These results hint at the importance of considering excitatory:inhibitory balance within an individual differences context, as what is optimal may differ based on a variety of neuroanatomical and physiological parameters.

Our findings are particularly striking given the nearly 10-year span between data collection waves. This suggests that EEG resting measures of periodic and aperiodic neural activity may be a promising biomarker for predicting who is at risk for cognitive decline. Given the relative ease and low cost of collecting EEG data, these metrics could be easily scalable to provide important information to clinicians for early interventions in a rapidly aging population. However, our sample is relatively modest in size and is composed of community-dwelling aging individuals who are able and willing to travel to participate in a multi-component study. Future work is needed to replicate these results in additional samples as well as investigate these measures in a variety of clinical samples and samples varying in demographic characteristics (including but not limited to race, ethnicity, education, and socioeconomic status) to further investigate the utility of IAPF and aperiodic exponent as a risk factor for accelerated cognitive decline. Particularly important would be a longitudinal study with repeated EEG and cognitive assessments completed at smaller time lags to assess *when* in aging measures of IAPF and aperiodic exponent signal increase the risk of cognitive decline.

Our results are, however, limited by the lack of resting EEG measures at both time points. Although the MIDUS Neuroscience M3 project was recently completed, EEG data was not recorded. The lack of a second measurement point prevents us from partially out the variance associated with longitudinal change in aperiodic activity and IAPF and examining whether this predicts a change in cognitive function. Moreover, we are unable to examine how individual differences in EEG predict cognitive change independently from the intra-individual changes. Given the substantial age-related differences (Hill et al., 2022; Merkin et al., 2022) and changes (Chini et al., 2022) in aperiodic activity and IAPF, we anticipate that the inclusion of a second measurement point would increase the sensitivity of our model.

In summary, our study highlights the importance of considering periodic and aperiodic measures in combination when examining resting-state EEG and measures of cognitive decline. In particular, a “mismatch” between low IAPF and high aperiodic exponent is associated with faster rates of cognitive decline over 10 years. Once considered meaningless, invariant noise, the features of the 1/*f* aperiodic neural activity are being recognized as an important feature of EEG signals, potentially reflecting global excitatory:inhibitory balance. Our work further emphasizes that aperiodic activity is a critical feature of EEG signals and needs to be systematically investigated in conjunction with more typical periodic features, to fully understand the links between neural activity and cognition across the lifespan.

## Acknowledgements

The Midlife in the United States (MIDUS) study was funded by the John D. and Catherine T. MacArthur Foundation Research Network and the National Institute on Aging (P01-AG020166, U19-AG051426, U01-AG077928). The MIDUS Neuroscience Project was also funded by Waisman Intellectual and Developmental Disabilities Research Center (U54-HD090256) awarded by the National Institute of Child Health and Human Development. Anna J. Finley was funded by the National Institute of Mental Health (F32-MH126537). Carien M. van Reekum was supported by the Biotechnology and Biological Sciences Research Council (BB/J009539/1 and BB/L02697X/1).

## Data availability

Data are available at https://midus.wisc.edu/data/index.php. All code used for all analyses and plots are publicly available on OSF at https://osf.io/sr4mb/.

Preregistration of analyses are available at https://osf.io/wyuca.

## Footnotes

1 Additional parallel analyses were conducted on eyes open and eyes closed data separately and are reported in the supplemental materials. Overall these analyses were consistent with the findings on the combined data.

2 The fronto-central composite of F3/Fz/F4 was comprised of the EGI GSN200 electrode montage (Electrical Geodesics, Inc, 2007) sensors 12, 20, 21, 25, 29 (comprising the analog for F3), sensors 4, 5, 118, 119, 124 (comprising the analog for F4), and sensor 11 (comprising the analog for Fz). Note this is an older montage than the EGI HydroCel nets.

3 As described in the Table 1 note, analyses reported in the manuscript deviate from the preregistered analyses, such that A) after careful examination for possible interactions with education and finding none, we decided to include education as a covariate instead of race, and B) the specific equations preregistered for analyses were overly conservative by including the interaction between the covariates with wave, as well as splitting up lag into separate terms instead of adding into a single term. These more conservative, complex analyses do not change the interpretations of our findings. Because the results do not change regardless of covariates or complexity of the analyses, we report the simplified analyses with education here, and report the preregistered analyses as well as simplified analyses with race instead of education as robustness checks in the supplemental materials.

