## Supplemental Materials for "Resting EEG Periodic and Aperiodic Components Predict Cognitive Decline Over 10 Years" (from another version of this preprint: version 3, 2024-01-25)

These supplemental analyses contain robustness analyses, including original preregistered models described at <https://osf.io/wyuca>, parallel models with and without controlling for different covariates, models controlling for genetic dependencies among twins, and exploratory models examining the relationship between aperiodic offset and cognitive decline as well as measures calculated from a parietal cluster from eyes-open and eyes-closed combined resting EEG data as well as a frontal cluster calculated separately from eyes-open-only and eyes-closed-only resting EEG data. For ease of the reader, Tables S1-S3 compares the key model terms for each analysis across the supplemental models on aperiodic exponent and individual peak alpha frequency (IAPF) calculated from the eyes-open and eyes-closed combined data. All analyses are reported in-full in subsequent sections. See Table 2 in “*Resting EEG Periodic and Aperiodic Components Predict Cognitive Decline Over 10 Years*” for a summary of each preregistered hypothesis.

We use the shorthand “simplified” refer to analyses that parallel what is described in the main manuscript (i.e., lag as a single term, no covariate by wave interaction), while “complex” refers to analyses that parallel what was explicitly preregistered (i.e., lag as two terns, covariate by wave interactions). We also delineate which covariates, if any, are accounted for in each variant model.

Age and lags are mean-centered. Education is coded as -1 = high school or less, 0 = some college, 1 = Bachelor’s degree or more. Sex coded as 0 = male, 1 = female. Race coded as 0 = White, 1 = BIPOC (Black, Indigenous, and People of Color). MIDUS Wave coded as 0 = MIDUS 2, 1 = MIDUS 3. Repeated measured cognitive data nested within participant, indicated by M2ID. For analyses controlling for genetic dependencies, participants are additionally nested within families, as indicated by M2ID:M2FAMNUM. Empirical Bayes slope estimation used handle missing data (Bates et al., 2015).

Figure S8 shows how associations between IPAF and cognitive impairment (and decline) are most pronounced in participants with greater aperiodic exponents. Table S49, S50, and Figure S9 show that the analyses reported in the main manuscript is similar – but weaker - when using a pooling of parietal electrodes. Finally, Figure S12 depicts the grand average power spectra for eyes open only, eyes closed only, and combined eyes open and eyes closed data.

**Comparison of Key Model Terms**

As shown in Table S1, none of the key terms significantly change by models.

| **Table S1:** Summary of key terms across various model specifications, BTACT composite score | | | | | | | |
| --- | --- | --- | --- | --- | --- | --- | --- |
|  | Simplified: Sex, Education, Total Lag (from manuscript) | Complex: Sex, Race, two Lags (preregistered) | Simplified: Sex, Race, Total Lag | Complex: Sex, Education, two Lags | Simplified: Family ID (for genetic dependencies) | Complex: Family ID (for genetic dependencies) | No Covariates |
| H1: Age main effect | **-0.02**  **[-0.03 – -0.02]**  ***p* < 0.001** | **-0.02**  **[-0.03– -0.01]**  ***p* < .001** | **-0.02**  **[-0.03– -0.02]**  ***p* < .001** | **-0.02**  **[-0.03– -0.01]**  ***p* < .001** | **-0.02**  **[-0.03– -0.02]**  ***p* < .001** | **-0.02**  **[-0.03– -0.01]**  ***p* < .001** | **-0.02**  **[-0.03– -0.01]**  ***p* < .001** |
| H2: Age X wave interaction | -0.00  [-0.01 – 0.01]  *p* = 0.707 | -0.00  [-0.01 – 0.00]  *p* = 0.334 | -0.00  [-0.01 – 0.01]  *p* = 0.704 | -0.00  [-0.01 – 0.00]  *p* = 0.319 | -0.00  [-0.01 – 0.01]  *p* = 0.693 | -0.00  [-0.01 – 0.00]  *p* = 0.293 | -0.00  [-0.01 – 0.01]  *p* = 0.712 |
| H3: Aperiodic exponent main effect | **0.39**  **[0.10 – 0.68]**  ***p* = 0.009** | **0.38**  **[0.09 – 0.67]**  ***p* = 0.010** | **0.43**  **[0.14 – 0.73]**  ***p* = 0.004** | **0.33**  **[0.04 – 0.62]**  ***p* = 0.024** | **0.37**  **[0.07 – 0.67]**  ***p* = 0.016** | **0.31**  **[0.01 – 0.60]**  ***p* = 0.042** | 0.29  [-0.02 – 0.59]  *p* = 0.067 |
| H4: Aperiodic exponent X wave interaction | -0.16  [-0.42 – 0. 09]  *p* = 0.204 | -0.13  [-0.38 – 0.13]  *p* = 0.327 | -0.17  [-0.42 – 0.08]  *p* = 0.175 | -0.11  [-0.36 – 0.14]  *p* = 0.396 | -0.17  [-0.42 – 0.08]  *p* = 0.176 | -0.11  [-0.36 – 0.14]  *p* = 0.375 | -0.17  [-0.42 – 0.08]  *p* = 0.177 |
| H5: IAPF main effect | 0.08  [-0.00 – 0.16]  *p* = 0.056 | 0.08  [-0.00 – 0.16]  *p* = 0.057 | 0.08  [-0.00 – 0.16]  *p* = 0.060 | 0.08  [-0.00 – 0.16]  *p* = 0.060 | **0.09**  **[0.00 – 0.17]**  ***p* = 0.039** | **0.09**  **[0.01 – 0.17]**  ***p* = 0.036** | **0.10**  **[0.01 – 0.19]**  ***p* = 0.027** |
| H6: IAPF X wave interaction | **0.07**  **[0.00 – 0.14]**  ***p* = 0.047** | 0.06  [-0.01 – 0.13]  *p* = 0.074 | 0.07  [-0.00 – 0.14]  *p* = 0.063 | 0.07  [-0.00 – 0.14]  *p* = 0.050 | 0.07  [-0.00 – 0.14]  *p* = 0.066 | 0.06  [-0.01 – 0.13]  *p* = 0.082 | 0.07  [-0.00 – 0.14]  *p* = 0.062 |
| H7a: Aperiodic exponent X age interaction | 0.01  [-0.02 – 0.03]  *p* = 0.695 | 0.01  [-0.02 – 0.04]  *p* = 0.528 | 0.01  [-0.02 – 0.04]  *p* = 0.556 | 0.01  [-0.02 – 0.03]  *p* = 0.668 | 0.01  [-0.02 – 0.04]  *p* = 0.493 | 0.01  [-0.02 – 0.04]  *p* = 0.444 | 0.01  [-0.02 – 0.04]  *p* = 0.440 |
| H7b: IAPF X age interaction | 0.00  [-0.00 – 0.01]  *p* = 0.374 | 0.00  [-0.00 – 0.01]  *p* = 0.420 | 0.00  [-0.00 – 0.01]  *p* = 0.288 | 0.00  [-0.01 – 0.01]  *p* = 0.555 | 0.00  [-0.00 – 0.01]  *p* = 0.346 | 0.00  [-0.01 – 0.01]  *p* = 0.531 | -0.00  [-0.01 – 0.01]  *p* = 0.989 |
| H7c: Aperiodic exponent X age X wave interaction | 0.00  [-0.02 – 0.03]  *p* = 0.858 | 0.00  [-0.02 – 0.03]  *p* = 0.898 | 0.00  [-0.02 – 0.03]  *p* = 0.774 | 0.00  [-0.03 – 0.03]  *p* = 0.950 | 0.00  [-0.02 – 0.03]  *p* = 0.816 | 0.00  [-0.03 – 0.03]  *p* = 0.990 | 0.00  [-0.02 – 0.03]  *p* = 0.737 |
| H7d: IAPF and age X wave interaction | -0.00  [-0.01 – 0.00]  *p* = 0.379 | -0.00  [-0.01 – 0.00]  *p* = 0.323 | -0.00  [-0.01 – 0.00]  *p* = 0.409 | -0.00  [-0.01 – 0.00]  *p* = 0.318 | -0.00  [-0.01 – 0.00]  *p* = 0.416 | -0.00  [-0.01 – 0.00]  *p* = 0.340 | -0.00  [-0.01 – 0.00]  *p* = 0.443 |
| Exploratory: Aperiodic Exponent X IAPF X Wave interaction | **0.33**  **[0.08 – 0.59]**  ***p* = 0.010** | 0.24  [-0.02 – 0.49]  *p* = 0.066 | **0.29**  **[0.04 – 0.54]**  ***p* = 0.021** | **0.27**  **[0.01 – 0.52]**  ***p* = 0.042** | **0.29**  **[0.04 – 0.54]**  ***p* = 0.021** | 0.23  [0.02 – 0.48]  *p* = 0.068 | **0.29**  **[0.04 – 0.54]**  ***p* = 0.022** |

| **Table S2:** Summary of key terms across various model specifications, Episodic Memory score | | | | | | | |
| --- | --- | --- | --- | --- | --- | --- | --- |
|  | Simplified: Sex, Education, Total Lag (from manuscript) | Complex: Sex, Race, two Lags (preregistered) | Simplified: Sex, Race, Total Lag | Complex: Sex, Education, two Lags | Simplified: Family ID (for genetic dependencies) | Complex: Family ID (for genetic dependencies) | No Covariates |
| H1: Age main effect | **-0.02**  **[-0.03 – -0.01]**  ***p* = 0.001** | **-0.02**  **[-0.03 – -0.01]**  ***p* = 0.004** | **-0.02**  **[-0.03 – -0.01]**  ***p* = 0.001** | **-0.02**  **[-0.03 – -0.01]**  ***p* = 0.005** | **-0.02**  **[-0.03 – -0.00]**  ***p* = 0.008** | **-0.01**  **[-0.03 – -0.00]**  ***p* = 0.022** | **-0.02**  **[-0.03 – -0.00]**  ***p* = 0.012** |
| H2: Age X wave interaction | -0.01  [-0.02 – 0.00]  *p* = 0.096 | **-0.01**  **[-0.03 – -0.00]**  ***p* = 0.035** | -0.01  [-0.02 – 0.00]  *p* = 0.088 | **-0.01**  **[-0.03 – -0.00]**  ***p* = 0.038** | -0.01  [-0.02 – 0.00]  *p* = 0.097 | **-0.01**  **[-0.02 – -0.00]**  ***p* = 0.048** | -0.01  [-0.02 – 0.00]  *p* = 0.099 |
| H3: Aperiodic exponent main effect | 0.26  [-0.19 – 0.71]  *p* = 0.254 | 0.24  [-0.21 – 0.69]  *p* = 0.300 | 0.29  [-0.16 – 0.74]  *p* = 0.201 | 0.21  [-0.24 – 0.65]  *p* = 0.366 | 0.15  [-0.32 – 0.63]  *p* = 0.531 | 0.11  [-0.36 – 0.59]  *p* = 0.642 | 0.12  [-0.35 – 0.59]  *p* = 0.624 |
| H4: Aperiodic exponent X wave interaction | -0.02  [-0.48 – 0.44]  *p* = 0.926 | 0.03  [-0.44 – 0.50]  *p* = 0.900 | -0.02  [-0.48 – 0.43]  *p* = 0.917 | -0.03  [-0.43 – 0.51]  *p* = 0.872 | -0.03  [-0.49 – 0.43]  *p* = 0.893 | -0.01  [-0.47 – 0.46]  *p* = 0.976 | -0.13  [-0.49 – 0.43]  *p* = 0.889 |
| H5: IAPF main effect | 0.12  [-0.00 – 0.25]  *p* = 0.055 | **0.13**  **[0.00 – 0.26]**  ***p* = 0.046** | **0.13**  **[0.00 – 0.26]**  ***p* = 0.046** | 0.12  [-0.00 – 0.25]  *p* = 0.054 | **0.15**  **[0.01 – 0.28]**  ***p* = 0.031** | **0.14**  **[0.01 – 0.28]**  ***p* = 0.045** | **0.15**  **[0.02 – 0.28]**  ***p* = 0.028** |
| H6: IAPF X wave interaction | 0.07  [-0.06 – 0.20]  *p* = 0.281 | 0.06  [-0.07 – 0.19]  *p* = 0.349 | 0.06  [-0.07 – 0.19]  *p* = 0.338 | 0.07  [-0.06 – 0.20]  *p* = 0.301 | 0.06  [-0.07 – 0.19]  *p* = 0.355 | 0.06  [-0.07 – 0.19]  *p* = 0.343 | 0.06  [-0.07 – 0.19]  *p* = 0.353 |
| H7a: Aperiodic exponent X age interaction | -0.01  [-0.06 – 0.04]  *p* = 0.677 | -0.01  [-0.05 – 0.04]  *p* = 0.818 | -0.01  [-0.05 – 0.04]  *p* = 0.745 | -0.01  [-0.06 – 0.04]  *p* = 0.747 | 0.00  [-0.05 – 0.05]  *p* = 0.863 | 0.01  [-0.04 – 0.05]  *p* = 0.839 | 0.01  [-0.02 – 0.08]  *p* = 0.841 |
| H7b: IAPF X age interaction | 0.00  [-0.01 – 0.01]  *p* = 0.836 | -0.00  [-0.01 – 0.01]  *p* = 0.989 | 0.00  [-0.01 – 0.01]  *p* = 0.744 | -0.00  [-0.01 – 0.01]  *p* = 0.902 | 0.00  [-0.01 – 0.01]  *p* = 0.812 | -0.00  [-0.01 – 0.01]  *p* = 0.922 | -0.01  [-0.02 – 0.00]  *p* = 0.161 |
| H7c: Aperiodic exponent X age X wave interaction | 0.03  [-0.02 – 0.08]  *p* = 0.201 | 0.03  [-0.02 – 0.08]  *p* = 0.253 | 0.03  [-0.02 – 0.08]  *p* = 0.187 | 0.03  [-0.02 – 0.08]  *p* = 0.263 | 0.03  [-0.08 – 0.08]  *p* = 0.182 | 0.03  [-0.02 – 0.08]  *p* = 0.197 | 0.03  [-0.02 – 0.08]  *p* = 0.175 |
| H7d: IAPF and age X wave interaction | 0.00  [-0.01 – 0.01]  *p* = 0.896 | 0.00  [-0.01 – 0.02]  *p* = 0.632 | 0.00  [-0.01 – 0.01]  *p* = 0.870 | 0.00  [-0.01 – 0.02]  *p* = 0.655 | 0.00  [-0.01 – 0.01]  *p* = 0.879 | 0.00  [-0.01 – 0.02]  *p* = 0.656 | 0.00  [-0.01 – 0.01]  *p* = 0.872 |
| Exploratory: Aperiodic Exponent X IAPF X Wave interaction | 0.20  [-0.26 – 0.67]  *p* = 0.397 | 0.11  [-0.37 – 0.58]  *p* = 0.657 | 0.15  [-0.31 – 0.62]  *p* = 0.514 | 0.15  [-0.33 – 0.63]  *p* = 0.535 | 0.12  [-0.34 – 0.59]  *p* = 0.600 | 0.11  [-0.37 – 0.58]  *p* = 0.659 | 0.15  [-0.31 – 0.62]  *p* = 0.524 |

| **Table S3:** Summary of key terms across various model specifications, Executive Function score | | | | | | | |
| --- | --- | --- | --- | --- | --- | --- | --- |
|  | Simplified: Sex, Education, Total Lag (from manuscript) | Complex: Sex, Race, two Lags (preregistered) | Simplified: Sex, Race, Total Lag | Complex: Sex, Education, two Lags | Simplified: Family ID (for genetic dependencies) | Complex: Family ID (for genetic dependencies) | No Covariates |
| H1: Age main effect | **-0.02**  **[-0.03 – -0.02]**  ***p* < 0.001** | **-0.02**  **[-0.03 – -0.01]**  ***p* < 0.001** | **-0.02**  **[-0.03 – -0.02]**  ***p* < 0.001** | **-0.02**  **[-0.03 – -0.01]**  ***p* < 0.001** | **-0.02**  **[-0.03 – -0.02]**  ***p* < 0.001** | **-0.02**  **[-0.03 – -0.01]**  ***p* < 0.001** | **-0.02**  **[-0.03 – -0.01]**  ***p* < 0.001** |
| H2: Age X wave interaction | 0.00  [-0.01 – 0.01]  *p* = 0.994 | -0.00  [-0.01 – 0.01]  *p* = 0.680 | 0.000  [-0.01 – 0.01]  *p* = 0.990 | -0.00  [-0.01 – 0.01]  *p* = 0.634 | -0.00  [-0.01 – 0.01]  *p* = 0.982 | -0.00  [-0.01 – 0.00]  *p* = 0.558 | -0.00  [-0.01 – 0.01]  *p* = 0.979 |
| H3: Aperiodic exponent main effect | **0.40**  **[0.09 – 0.71]**  ***p* = 0.012** | **0.40**  **[0.09 – 0.71]**  ***p* = 0.011** | **0.45**  **[0.14 – 0.76]**  ***p* = 0.005** | **0.35**  **[0.04 – 0.65]**  ***p* = 0.027** | **0.40**  **[0.09 – 0.72]**  ***p* = 0.013** | **0.34**  **[0.03 – 0.65]**  ***p* = 0.031** | 0.31  [-0.01 – 0.63]  *p* = 0.059 |
| H4: Aperiodic exponent X wave interaction | -0.21  [-0.47 – 0.05]  *p* = 0.120 | -0.18  [-0.44 – 0.08]  *p* = 0.170 | -0.22  [-0.48 – 0.04]  *p* = 0.101 | -0.16  [-0.42 – 0.10]  *p* = 0.224 | -0.21  [-0.47 – 0.05]  *p* = 0.106 | -0.16  [-0.42 – 0.10]  *p* = 0.228 | -0.21  [-0.47 – 0.05]  *p* = 0.111 |
| H5: IAPF main effect | 0.07  [-0.02 – 0.16]  *p* = 0.115 | 0.07  [-0.02 – 0.16]  *p* = 0.105 | 0.07  [-0.02 – 0.16]  *p* = 0.109 | 0.07  [-0.06 – 0.28]  *p* = 0.124 | 0.08  [-0.01 – 0.17]  *p* = 0.086 | 0.08  [-0.01 – 0.17]  *p* = 0.078 | 0.09  [-0.00 – 0.18]  *p* = 0.061 |
| H6: IAPF X wave interaction | 0.07  [-0.00 – 0.15]  *p* = 0.061 | 0.06  [-0.01 – 0.13]  *p* = 0.100 | 0.06  [-0.01 – 0.14]  *p* = 0.091 | 0.07  [-0.00 – 0.14]  *p* = 0.060 | 0.06  [-0.01 – 0.14]  *p* = 0.093 | 0.06  [-0.02 – 0.13]  *p* = 0.121 | 0.06  [-0.01 – 0.14]  *p* = 0.092 |
| H7a: Aperiodic exponent X age interaction | 0.01  [-0.02 – 0.04]  *p* = 0.632 | 0.01  [-0.02 – 0.04]  *p* = 0.484 | 0.01  [-0.02 – 0.04]  *p* = 0.495 | 0.01  [-0.02 – 0.04]  *p* = 0.626 | 0.01  [-0.02 – 0.04]  *p* = 0.524 | 0.01  [-0.02 – 0.04]  *p* = 0.472 | 0.01  [-0.02 – 0.05]  *p* = 0.460 |
| H7b: IAPF X age interaction | 0.00  [-0.00 – 0.01]  *p* = 0.350 | 0.00  [-0.00 – 0.01]  *p* = 0.335 | 0.00  [-0.00 – 0.01]  *p* = 0.257 | 0.00  [-0.01 – 0.01]  *p* = 0.481 | 0.00  [-0.00 – 0.01]  *p* = 0.311 | 0.00  [-0.00 – 0.01]  *p* = 0.430 | 0.00  [-0.01 – 0.01]  *p* = 0.807 |
| H7c: Aperiodic exponent X age X wave interaction | 0.00  [-0.02 – 0.03]  *p* = 0.755 | 0.01  [-0.02 – 0.03]  *p* = 0.709 | 0.01  [-0.02 – 0.03]  *p* = 0.690 | 0.00  [-0.02 – 0.04]  *p* = 0.730 | 0.00  [-0.02 – 0.03]  *p* = 0.725 | 0.00  [-0.02 – 0.03]  *p* = 0.836 | 0.01  [-0.02 – 0.03]  *p* = 0.708 |
| H7d: IAPF and age X wave interaction | -0.00  [-0.01 – 0.00]  *p* = 0.301 | -0.00  [-0.01 – 0.00]  *p* = 0.173 | -0.00  [-0.01 – 0.00]  *p* = 0.303 | -0.00  [-0.01 – 0.00]  *p* = 0.193 | -0.00  [-0.01 – 0.00]  *p* = 0.302 | -0.00  [-0.01 – 0.00]  *p* = 0.182 | -0.00  [-0.01 – 0.00]  *p* = 0.309 |
| Exploratory: Aperiodic Exponent X IAPF X Wave interaction | **0.34**  **[0.07 – 0.61]**  ***p* = 0.013** | 0.25  [-0.02 – 0.52]  *p* = 0.067 | **0.31**  **[0.04 – 0.57]**  ***p* = 0.024** | **0.27**  **[0.00 – 0.54]**  ***p* = 0.048** | **0.31**  **[0.04 – 0.57]**  ***p* = 0.023** | 0.24  [-0.02 – 0.51]  *p* = 0.072 | **0.30**  **[0.004 – 0.57]**  ***p* = 0.024** |

**Preregistered Analyses: Complex analyses with sex, race, and lag split into 2 parts**

| **Table S4:** Complex multilevel models to test hypothesis 1 and 2. Sex, race, and lag in 2 parts as covariates. | | | | | | | | | |
| --- | --- | --- | --- | --- | --- | --- | --- | --- | --- |
|  | **Episodic Memory** | | | **Executive Functioning** | | | **BTACT** | | |
| *Predictors* | *Estimates* | *CI* | *p* | *Estimates* | *CI* | *p* | *Estimates* | *CI* | *p* |
| Intercept (M2) | **-0.33** | **-0.54 – -0.12** | **0.002** | 0.01 | -0.13 – 0.14 | 0.922 | -0.05 | -0.17 – 0.08 | 0.487 |
| MIDUS Wave | **-0.26** | **-0.48 – -0.04** | **0.019** | **-0.38** | **-0.51 – -0.26** | **<0.001** | **-0.36** | **-0.48 – -0.24** | **<0.001** |
| Age | **-0.02** | **-0.03 – -0.01** | **0.004** | **-0.02** | **-0.03 – -0.01** | **<0.001** | **-0.02** | **-0.03 – -0.01** | **<0.001** |
| Sex | **0.63** | **0.39 – 0.88** | **<0.001** | 0.16 | -0.00 – 0.32 | 0.054 | **0.24** | **0.09 – 0.39** | **0.002** |
| Race | -0.18 | -0.54 – 0.18 | 0.335 | **-0.41** | **-0.64 – -0.17** | **0.001** | **-0.36** | **-0.59 – -0.14** | **0.001** |
| M2 Cog to Neuro Lag | 0.08 | -0.13 – 0.28 | 0.473 | 0.05 | -0.08 – 0.19 | 0.432 | 0.06 | -0.07 – 0.19 | 0.360 |
| Neuro to M3 Cog Lag | -0.02 | -0.19 – 0.16 | 0.845 | -0.01 | -0.13 – 0.10 | 0.827 | -0.01 | -0.12 – 0.09 | 0.795 |
| Wave X Age | **-0.01** | **-0.03 – -0.00** | **0.035** | -0.00 | -0.01 – 0.01 | 0.680 | -0.00 | -0.01 – 0.00 | 0.334 |
| Wave X Sex | 0.20 | -0.07 – 0.46 | 0.141 | -0.09 | -0.24 – 0.05 | 0.221 | -0.04 | -0.18 – 0.10 | 0.576 |
| Wave X Race | 0.05 | -0.34 – 0.43 | 0.815 | 0.02 | -0.19 – 0.24 | 0.845 | 0.00 | -0.20 – 0.21 | 0.977 |
| Wave X M2 Cog to Neuro Lag | -0.15 | -0.37 – 0.07 | 0.175 | **-0.15** | **-0.28 – -0.03** | **0.018** | **-0.16** | **-0.28 – -0.04** | **0.008** |
| Wave X Neuro to M3 Cog Lag | -0.06 | -0.24 – 0.12 | 0.519 | **-0.14** | **-0.24 – -0.03** | **0.011** | **-0.13** | **-0.22 – -0.03** | **0.012** |
| **Random Effects** | | | | | | | | | |
| σ^2^ | 0.46 | | | 0.15 | | | 0.13 | | |
| τ_00_ | 0.40 _M2ID_ | | | 0.23 _M2ID_ | | | 0.19 _M2ID_ | | |
| ICC | 0.46 | | | 0.61 | | | 0.60 | | |
| N | 234 _M2ID_ | | | 235 _M2ID_ | | | 234 _M2ID_ | | |
| Observations | 451 | | | 458 | | | 443 | | |
| Marginal R^2^ / Conditional R^2^ | 0.190 / 0.566 | | | 0.320 / 0.732 | | | 0.332 / 0.734 | | |

| **Table S5:** Complex multilevel models to test hypothesis 3 and 4. Sex, race, and lag in 2 parts as covariates. | | | | | | | | |  |
| --- | --- | --- | --- | --- | --- | --- | --- | --- | --- |
| **Episodic Memory** | | | **Executive Functioning** | | | **BTACT** | | |  |
| *Predictors* | *Estimates* | *CI* | *p* | *Estimates* | *CI* | *p* | *Estimates* | *CI* | *p* |
| Intercept (M2) | **-0.31** | **-0.53 – -0.10** | **0.005** | 0.03 | -0.12 – 0.17 | 0.735 | -0.03 | -0.17 – 0.11 | 0.682 |
| MIDUS Wave | **-0.25** | **-0.47 – -0.03** | **0.028** | **-0.38** | **-0.50 – -0.26** | **<0.001** | **-0.36** | **-0.47 – -0.24** | **<0.001** |
| Exponent | 0.24 | -0.21 – 0.69 | 0.300 | **0.40** | **0.09 – 0.71** | **0.011** | **0.38** | **0.09 – 0.67** | **0.010** |
| Sex | **0.62** | **0.36 – 0.87** | **<0.001** | 0.14 | -0.03 – 0.31 | 0.115 | **0.22** | **0.06 – 0.39** | **0.008** |
| Race | -0.19 | -0.56 – 0.19 | 0.325 | **-0.43** | **-0.68 – -0.17** | **0.001** | **-0.38** | **-0.62 – -0.14** | **0.002** |
| M2 Cog to Neuro Lag | 0.12 | -0.09 – 0.33 | 0.266 | 0.11 | -0.03 – 0.26 | 0.124 | 0.11 | -0.02 – 0.25 | 0.097 |
| Neuro to M3 Cog Lag | 0.01 | -0.17 – 0.18 | 0.953 | 0.02 | -0.10 – 0.14 | 0.792 | 0.01 | -0.10 – 0.13 | 0.818 |
| Wave X Exponent | 0.03 | -0.44 – 0.50 | 0.900 | -0.18 | -0.44 – 0.08 | 0.170 | -0.13 | -0.38 – 0.13 | 0.327 |
| Wave X Sex | 0.18 | -0.09 – 0.44 | 0.192 | -0.10 | -0.25 – 0.04 | 0.165 | -0.05 | -0.19 – 0.09 | 0.478 |
| Wave X Race | 0.06 | -0.33 – 0.44 | 0.770 | 0.03 | -0.19 – 0.24 | 0.794 | 0.01 | -0.20 – 0.22 | 0.916 |
| Wave X M2 Cog to Neuro Lag | -0.11 | -0.33 – 0.11 | 0.335 | **-0.14** | **-0.26 – -0.02** | **0.026** | **-0.14** | **-0.26 – -0.02** | **0.019** |
| Wave X Neuro to M3 Cog Lag | -0.04 | -0.22 – 0.14 | 0.669 | **-0.13** | **-0.23 – -0.02** | **0.015** | **-0.11** | **-0.21 – -0.02** | **0.022** |
| **Random Effects** | | | | | | | | | |
| σ^2^ | 0.47 | | | 0.15 | | | 0.13 | | |
| τ_00_ | 0.45 _M2ID_ | | | 0.28 _M2ID_ | | | 0.25 _M2ID_ | | |
| ICC | 0.49 | | | 0.66 | | | 0.66 | | |
| N | 234 _M2ID_ | | | 235 _M2ID_ | | | 234 _M2ID_ | | |
| Observations | 451 | | | 458 | | | 443 | | |
| Marginal R^2^ / Conditional R^2^ | 0.135 / 0.555 | | | 0.227 / 0.735 | | | 0.225 / 0.734 | | |

| **Table S6:** Complex multilevel models to test hypothesis 5 and 6. Sex, race, and lag in 2 parts as covariates. | | | | | | | | | |
| --- | --- | --- | --- | --- | --- | --- | --- | --- | --- |
|  | **Episodic Memory** | | | **Executive Functioning** | | | **BTACT** | | |
| *Predictors* | *Estimates* | *CI* | *p* | *Estimates* | *CI* | *p* | *Estimates* | *CI* | *p* |
| Intercept (M2) | **-0.31** | **-0.52 – -0.09** | **0.005** | 0.03 | -0.11 – 0.18 | 0.644 | -0.02 | -0.16 – 0.12 | 0.790 |
| MIDUS Wave | **-0.25** | **-0.47 – -0.03** | **0.029** | **-0.38** | **-0.51 – -0.26** | **<0.001** | **-0.36** | **-0.47 – -0.24** | **<0.001** |
| Alpha Peak | **0.13** | **0.00 – 0.26** | **0.046** | 0.07 | -0.02 – 0.16 | 0.105 | 0.08 | -0.00 – 0.16 | 0.057 |
| Sex | **0.59** | **0.34 – 0.84** | **<0.001** | 0.11 | -0.06 – 0.28 | 0.205 | **0.19** | **0.03 – 0.35** | **0.021** |
| Race | -0.16 | -0.53 – 0.21 | 0.404 | **-0.40** | **-0.65 – -0.14** | **0.002** | **-0.35** | **-0.59 – -0.11** | **0.004** |
| M2 Cog to Neuro Lag | 0.13 | -0.08 – 0.34 | 0.212 | 0.13 | -0.01 – 0.27 | 0.070 | 0.13 | -0.00 – 0.27 | 0.051 |
| Neuro to M3 Cog Lag | 0.02 | -0.16 – 0.19 | 0.865 | 0.03 | -0.09 – 0.15 | 0.624 | 0.03 | -0.09 – 0.14 | 0.642 |
| Wave X Alpha Peak | 0.06 | -0.07 – 0.19 | 0.349 | 0.06 | -0.01 – 0.13 | 0.100 | 0.06 | -0.01 – 0.13 | 0.074 |
| Wave X Sex | 0.17 | -0.09 – 0.43 | 0.206 | -0.10 | -0.24 – 0.05 | 0.181 | -0.05 | -0.19 – 0.09 | 0.490 |
| Wave X Race | 0.07 | -0.32 – 0.45 | 0.734 | 0.02 | -0.19 – 0.24 | 0.838 | 0.01 | -0.20 – 0.21 | 0.950 |
| Wave X M2 Cog to Neuro Lag | -0.10 | -0.32 – 0.11 | 0.345 | **-0.14** | **-0.27 – -0.02** | **0.019** | **-0.14** | **-0.26 – -0.03** | **0.015** |
| Wave X Neuro to M3 Cog Lag | -0.04 | -0.22 – 0.15 | 0.693 | **-0.13** | **-0.23 – -0.03** | **0.012** | **-0.12** | **-0.21 – -0.02** | **0.019** |
| **Random Effects** | | | | | | | | | |
| σ^2^ | 0.47 | | | 0.14 | | | 0.13 | | |
| τ_00_ | 0.43 _M2ID_ | | | 0.28 _M2ID_ | | | 0.24 _M2ID_ | | |
| ICC | 0.47 | | | 0.66 | | | 0.66 | | |
| N | 234 _M2ID_ | | | 235 _M2ID_ | | | 234 _M2ID_ | | |
| Observations | 451 | | | 458 | | | 443 | | |
| Marginal R^2^ / Conditional R^2^ | 0.155 / 0.556 | | | 0.231 / 0.736 | | | 0.235 / 0.738 | | |

| **Table S7:** Complex multilevel models to test hypothesis 7a and 7c. Sex, race, and lag in 2 parts as covariates. | | | | | | | | | |
| --- | --- | --- | --- | --- | --- | --- | --- | --- | --- |
|  | **Episodic Memory** | | | **Executive Functioning** | | | **BTACT** | | |
| *Predictors* | *Estimates* | *CI* | *p* | *Estimates* | *CI* | *p* | *Estimates* | *CI* | *p* |
| (Intercept) | **-0.34** | **-0.56 – -0.12** | **0.003** | 0.02 | -0.13 – 0.16 | 0.803 | -0.04 | -0.17 – 0.10 | 0.591 |
| MIDUS Wave | -0.23 | -0.46 – 0.00 | 0.051 | **-0.38** | **-0.50 – -0.25** | **<0.001** | **-0.36** | **-0.48 – -0.23** | **<0.001** |
| Exponent | 0.08 | -0.38 – 0.53 | 0.739 | 0.18 | -0.11 – 0.48 | 0.226 | 0.17 | -0.11 – 0.45 | 0.230 |
| Age | **-0.02** | **-0.03 – -0.00** | **0.008** | **-0.02** | **-0.03 – -0.01** | **<0.001** | **-0.02** | **-0.03 – -0.01** | **<0.001** |
| Sex | **0.64** | **0.39 – 0.89** | **<0.001** | 0.16 | -0.00 – 0.33 | 0.051 | **0.24** | **0.09 – 0.40** | **0.002** |
| Race | -0.18 | -0.55 – 0.18 | 0.331 | **-0.42** | **-0.66 – -0.18** | **0.001** | **-0.38** | **-0.60 – -0.15** | **0.001** |
| M2 Cog to Neuro Lag | 0.07 | -0.14 – 0.28 | 0.494 | 0.05 | -0.08 – 0.19 | 0.449 | 0.06 | -0.07 – 0.19 | 0.382 |
| Neuro to M3 Cog Lag | -0.02 | -0.19 – 0.15 | 0.826 | -0.01 | -0.13 – 0.10 | 0.811 | -0.02 | -0.12 – 0.09 | 0.771 |
| Wave X Exponent | -0.10 | -0.58 – 0.38 | 0.671 | -0.21 | -0.48 – 0.06 | 0.122 | -0.16 | -0.42 – 0.10 | 0.223 |
| Wave X Age | -0.01 | -0.03 – 0.00 | 0.053 | -0.00 | -0.01 – 0.00 | 0.470 | -0.00 | -0.01 – 0.00 | 0.228 |
| Exponent X Age | -0.01 | -0.05 – 0.04 | 0.818 | 0.01 | -0.02 – 0.04 | 0.484 | 0.01 | -0.02 – 0.04 | 0.528 |
| Wave X Sex | 0.18 | -0.09 – 0.44 | 0.188 | -0.10 | -0.25 – 0.04 | 0.171 | -0.05 | -0.19 – 0.09 | 0.493 |
| Wave X Race | 0.04 | -0.34 – 0.42 | 0.834 | 0.03 | -0.19 – 0.24 | 0.800 | 0.01 | -0.20 – 0.22 | 0.931 |
| Wave X M2 Cog to Neuro Lag | -0.14 | -0.36 – 0.08 | 0.209 | **-0.14** | **-0.27 – -0.02** | **0.024** | **-0.15** | **-0.27 – -0.03** | **0.012** |
| Wave X Neuro to M3 Cog Lag | -0.05 | -0.23 – 0.13 | 0.585 | **-0.13** | **-0.23 – -0.03** | **0.014** | **-0.12** | **-0.22 – -0.02** | **0.016** |
| Wave X Exponent X Age | 0.03 | -0.02 – 0.08 | 0.253 | 0.01 | -0.02 – 0.03 | 0.709 | 0.00 | -0.02 – 0.03 | 0.898 |
| **Random Effects** | | | | | | | | | |
| σ^2^ | 0.46 | | | 0.15 | | | 0.13 | | |
| τ_00_ | 0.41 _M2ID_ | | | 0.23 _M2ID_ | | | 0.20 _M2ID_ | | |
| ICC | 0.47 | | | 0.61 | | | 0.60 | | |
| N | 234 _M2ID_ | | | 235 _M2ID_ | | | 234 _M2ID_ | | |
| Observations | 451 | | | 458 | | | 443 | | |
| Marginal R^2^ / Conditional R^2^ | 0.190 / 0.569 | | | 0.323 / 0.735 | | | 0.334 / 0.735 | | |

| **Table S8:** Complex multilevel models to test hypothesis 7b and 7d. Sex, race, and lag in 2 parts as covariates. | | | | | | | | | | | |
| --- | --- | --- | --- | --- | --- | --- | --- | --- | --- | --- | --- |
|  | **Episodic Memory** | | | **Executive Functioning** | | | | **BTACT** | | | |
| *Predictors* | *Estimates* | *CI* | *p* | *Estimates* | | *CI* | *p* | *Estimates* | | *CI* | *p* |
| (Intercept) | **-0.32** | **-0.54 – -0.11** | **0.003** | 0.02 | | -0.12 – 0.16 | 0.812 | -0.04 | | -0.17 – 0.09 | 0.575 |
| MIDUS Wave | **-0.25** | **-0.48 – -0.03** | **0.027** | **-0.39** | | **-0.52 – -0.27** | **<0.001** | **-0.37** | | **-0.48 – -0.25** | **<0.001** |
| Alpha Peak | 0.09 | -0.04 – 0.22 | 0.174 | 0.02 | | -0.07 – 0.10 | 0.675 | 0.03 | | -0.05 – 0.11 | 0.513 |
| Age | **-0.01** | **-0.03 – -0.00** | **0.016** | **-0.02** | | **-0.03 – -0.01** | **<0.001** | **-0.02** | | **-0.03 – -0.01** | **<0.001** |
| Sex | **0.62** | **0.37 – 0.87** | **<0.001** | 0.16 | | -0.00 – 0.32 | 0.053 | **0.24** | | **0.09 – 0.39** | **0.002** |
| Race | -0.17 | -0.53 – 0.20 | 0.370 | **-0.41** | | **-0.65 – -0.17** | **0.001** | **-0.36** | | **-0.59 – -0.14** | **0.001** |
| M2 Cog to Neuro Lag | 0.08 | -0.12 – 0.29 | 0.427 | 0.05 | | -0.08 – 0.19 | 0.434 | 0.06 | | -0.07 – 0.19 | 0.355 |
| Neuro to M3 Cog Lag | -0.01 | -0.19 – 0.16 | 0.895 | -0.01 | | -0.12 – 0.11 | 0.897 | -0.01 | | -0.12 – 0.10 | 0.859 |
| Wave X Alpha Peak | 0.03 | -0.10 – 0.17 | 0.637 | 0.06 | | -0.02 – 0.13 | 0.153 | 0.05 | | -0.02 – 0.12 | 0.166 |
| Wave X Age | -0.01 | -0.02 – 0.00 | 0.068 | -0.00 | | -0.01 – 0.01 | 0.891 | -0.00 | | -0.01 – 0.00 | 0.535 |
| Alpha Peak X Age | -0.00 | -0.01 – 0.01 | 0.989 | 0.00 | | -0.00 – 0.01 | 0.335 | 0.00 | | -0.00 – 0.01 | 0.420 |
| Wave X Sex | 0.19 | -0.07 – 0.46 | 0.146 | -0.10 | | -0.25 – 0.04 | 0.166 | -0.05 | | -0.19 – 0.09 | 0.477 |
| Wave X Race | 0.04 | -0.34 – 0.43 | 0.820 | 0.03 | | -0.18 – 0.25 | 0.776 | 0.01 | | -0.20 – 0.22 | 0.921 |
| Wave X M2 Cog to Neuro Lag | -0.15 | -0.37 – 0.07 | 0.186 | **-0.14** | | **-0.27 – -0.02** | **0.025** | **-0.15** | | **-0.27 – -0.03** | **0.013** |
| Wave X Neuro to M3 Cog Lag | -0.05 | -0.24 – 0.13 | 0.568 | **-0.14** | | **-0.24 – -0.03** | **0.010** | **-0.12** | | **-0.22 – -0.03** | **0.014** |
| Wave X Alpha Peak X Age | 0.00 | -0.01 – 0.02 | 0.632 | -0.00 | | -0.01 – 0.00 | 0.173 | -0.00 | | -0.01 – 0.00 | 0.323 |
| **Random Effects** | | | | | | | | | | | |
| σ^2^ | | 0.47 | | | 0.14 | | | | 0.13 | | |
| τ_00_ | | 0.39 _M2ID_ | | | 0.23 _M2ID_ | | | | 0.20 _M2ID_ | | |
| ICC | | 0.46 | | | 0.61 | | | | 0.61 | | |
| N | | 234 _M2ID_ | | | 235 _M2ID_ | | | | 234 _M2ID_ | | |
| Observations | | 451 | | | 458 | | | | 443 | | |
| Marginal R^2^ / Conditional R^2^ | | 0.198 / 0.565 | | | 0.324 / 0.738 | | | | 0.337 / 0.739 | | |

| **Table S9:** Complex multilevel model, exploratory analyses. Sex, race, and lag in 2 parts as covariates. | | | | | | | | | |
| --- | --- | --- | --- | --- | --- | --- | --- | --- | --- |
|  | **Episodic Memory** | | | **Executive Functioning** | | | **BTACT** | | |
| *Predictors* | *Estimates* | *CI* | *p* | *Estimates* | *CI* | *p* | *Estimates* | *CI* | *p* |
| Intercept (M2) | **-0.33** | **-0.54 – -0.12** | **0.002** | -0.00 | -0.14 – 0.14 | 0.986 | -0.05 | -0.18 – 0.08 | 0.423 |
| MIDUS Wave | **-0.26** | **-0.48 – -0.03** | **0.025** | **-0.37** | **-0.49 – -0.24** | **<0.001** | **-0.34** | **-0.46 – -0.23** | **<0.001** |
| Exponent | 0.14 | -0.34 – 0.63 | 0.562 | 0.18 | -0.14 – 0.50 | 0.279 | 0.17 | -0.13 – 0.47 | 0.262 |
| Alpha Peak | 0.10 | -0.03 – 0.23 | 0.135 | 0.03 | -0.06 – 0.11 | 0.532 | 0.04 | -0.04 – 0.12 | 0.382 |
| Age | **-0.01** | **-0.03 – -0.00** | **0.036** | **-0.02** | **-0.03 – -0.01** | **<0.001** | **-0.02** | **-0.03 – -0.01** | **<0.001** |
| Sex | **0.63** | **0.38 – 0.88** | **<0.001** | **0.17** | **0.01 – 0.33** | **0.042** | **0.25** | **0.10 – 0.40** | **0.002** |
| Race | -0.17 | -0.54 – 0.19 | 0.350 | **-0.42** | **-0.65 – -0.18** | **0.001** | **-0.37** | **-0.59 – -0.15** | **0.001** |
| M2 Cog to Neuro Lag | 0.08 | -0.13 – 0.29 | 0.468 | 0.04 | -0.09 – 0.18 | 0.528 | 0.05 | -0.08 – 0.18 | 0.443 |
| Neuro to M3 Cog Lag | -0.02 | -0.19 – 0.16 | 0.850 | -0.02 | -0.14 – 0.09 | 0.717 | -0.02 | -0.13 – 0.09 | 0.686 |
| Wave X Exponent | -0.04 | -0.55 – 0.48 | 0.888 | -0.10 | -0.38 – 0.18 | 0.481 | -0.05 | -0.33 – 0.22 | 0.703 |
| Wave X Alpha Peak | 0.02 | -0.12 – 0.16 | 0.753 | 0.05 | -0.03 – 0.12 | 0.237 | 0.04 | -0.03 – 0.12 | 0.230 |
| Exponent X Alpha Peak | -0.05 | -0.50 – 0.41 | 0.843 | -0.12 | -0.41 – 0.18 | 0.445 | -0.11 | -0.39 – 0.17 | 0.437 |
| Wave X Age | -0.01 | -0.03 – 0.00 | 0.066 | -0.00 | -0.01 – 0.01 | 0.844 | -0.00 | -0.01 – 0.01 | 0.558 |
| Wave X Sex | 0.19 | -0.08 – 0.45 | 0.162 | -0.11 | -0.25 – 0.04 | 0.138 | -0.06 | -0.20 – 0.08 | 0.428 |
| Wave X Race | 0.05 | -0.33 – 0.44 | 0.788 | 0.03 | -0.18 – 0.25 | 0.752 | 0.01 | -0.19 – 0.22 | 0.886 |
| Wave X M2 Cog to Neuro Lag | -0.14 | -0.36 – 0.08 | 0.222 | **-0.13** | **-0.25 – -0.00** | **0.045** | **-0.14** | **-0.26 – -0.02** | **0.024** |
| Wave X Neuro to M3 Cog Lag | -0.05 | -0.23 – 0.14 | 0.599 | **-0.12** | **-0.22 – -0.01** | **0.028** | **-0.11** | **-0.21 – -0.01** | **0.033** |
| Wave X Exponent X Alpha Peak | 0.11 | -0.37 – 0.58 | 0.657 | 0.25 | -0.02 – 0.52 | 0.067 | 0.24 | -0.02 – 0.49 | 0.066 |
| **Random Effects** | | | | | | | | | |
| σ^2^ | 0.47 | | | 0.14 | | | 0.13 | | |
| τ_00_ | 0.39 _M2ID_ | | | 0.23 _M2ID_ | | | 0.20 _M2ID_ | | |
| ICC | 0.46 | | | 0.62 | | | 0.61 | | |
| N | 234 _M2ID_ | | | 235 _M2ID_ | | | 234 _M2ID_ | | |
| Observations | 451 | | | 458 | | | 443 | | |
| Marginal R^2^ / Conditional R^2^ | 0.198 / 0.565 | | | 0.326 / 0.741 | | | 0.340 / 0.743 | | |

| **Table S10:** Examining the Wave X Aperiodic Exponent X Individual Peak Alpha Frequency interaction from analyses described in table S9 (complex multilevel models with sex, race, and lag in two parts as covariates) through the slope of the change in cognitive function over waves for each BTACT measure at high and low levels of each EEG metric. | | | | | | |
| --- | --- | --- | --- | --- | --- | --- |
| **Change in BTACT Episodic Memory Factor from M2 to M3** | | | | | | |
|  | *Slope of IAPF* | | *95% CI* | | *p-value* | |
| Low Exponent (-1 SD) | -0.01 | | -0.20 – 0.19 | | 0.939 | |
| High Exponent (+1 SD) | 0.05 | | -0.14 – 0.24 | | 0.586 | |
|  | *Slope of Exponent* | | *95% CI* | | *p-value* | |
| Low IAPF (-1 SD) | -0.14 | | -0.73 –0.45 | | 0.636 | |
| High IAPF (+1 SD) | 0.07 | | -0.72 – 0.85 | | 0.865 | |
| **Change in BTACT Executive Functioning Factor from M2 to M3** | | | | | | |
|  | | *Slope of IAPF* | | *95% CI* | | *p-value* |
| Low Exponent (-1 SD) | | -0.02 | | -0.13 – 0.09 | | 0.683 |
| High Exponent (+1 SD) | | **0.12** | | **0.01 – 0.22** | | **0.030** |
|  | | *Slope of Exponent* | | *95% CI* | | *p-value* |
| Low IAPF (-1 SD) | | **-0.35** | | **-0.68 – -0.01** | | **0.043** |
| High IAPF (+1 SD) | | 0.14 | | -0.29 – 0.57 | | 0.515 |
| **Change in BTACT Overall Composite from M2 to M3** | | | | | | |
|  | | *Slope of IAPF* | | *95% CI* | | *p-value* |
| Low Exponent (-1 SD) | | -0.02 | | -0.13 – 0.08 | | 0.696 |
| High Exponent (+1 SD) | | **0.11** | | **0.01 – 0.21** | | **0.028** |
|  | | *Slope of Exponent* | | *95% CI* | | *p-value* |
| Low IAPF (-1 SD) | | -0.28 | | -0.60 – 0.03 | | 0.081 |
| High IAPF (+1 SD) | | 0.18 | | -0.23 – 0.59 | | 0.393 |

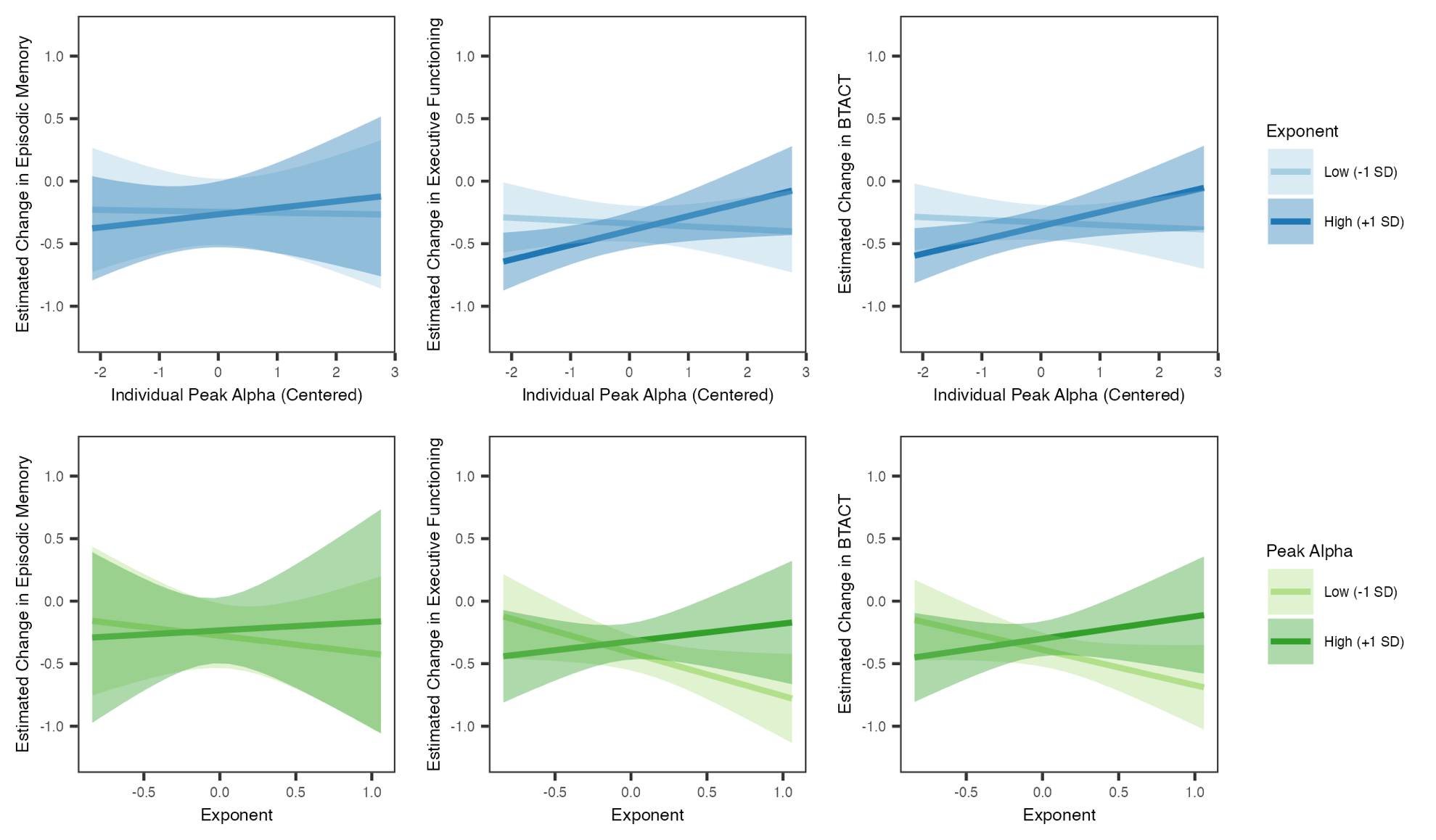

Figure S1. Wave by aperiodic exponent by individual peak alpha frequency interaction plot for models described in Table S9 (complex multilevel models with sex, race, and lag in two parts as covariates).

**Analyses Parallel to Manuscript: Simplified analyses with sex, race, and total lag (1 part)**

| **Table S11:** Simple multilevel models to test Hypotheses 1 and 2. Sex, race, and lag in 1 part as covariates. | | | | | | | | | |
| --- | --- | --- | --- | --- | --- | --- | --- | --- | --- |
|  | **Episodic Memory** | | | **Executive Functioning** | | | **BTACT** | | |
| *Predictors* | *Estimates* | *CI* | *p* | *Estimates* | *CI* | *p* | *Estimates* | *CI* | *p* |
| Intercept (M2) | **-0.39** | **-0.58 – -0.20** | **<0.001** | 0.03 | -0.10 – 0.16 | 0.609 | -0.03 | -0.15 – 0.09 | 0.586 |
| MIDUS Wave | **-0.13** | **-0.26 – -0.01** | **0.040** | **-0.43** | **-0.50 – -0.36** | **<0.001** | **-0.38** | **-0.45 – -0.32** | **<0.001** |
| Age | **-0.02** | **-0.03 – -0.01** | **0.001** | **-0.02** | **-0.03 – -0.02** | **<0.001** | **-0.02** | **-0.03 – -0.02** | **<0.001** |
| Sex | **0.73** | **0.52 – 0.95** | **<0.001** | 0.12 | -0.03 – 0.26 | 0.107 | **0.23** | **0.09 – 0.36** | **0.001** |
| Race | -0.17 | -0.48 – 0.14 | 0.276 | **-0.42** | **-0.64 – -0.21** | **<0.001** | **-0.39** | **-0.59 – -0.19** | **<0.001** |
| Lag between Waves | -0.06 | -0.20 – 0.09 | 0.451 | -0.09 | -0.19 – 0.01 | 0.087 | -0.09 | -0.18 – 0.01 | 0.080 |
| MIDUS Wave X Age | -0.01 | -0.02 – 0.00 | 0.088 | 0.00 | -0.01 – 0.01 | 0.990 | -0.00 | -0.01 – 0.01 | 0.704 |
| **Random Effects** | | | | | | | | | |
| σ^2^ | 0.46 | | | 0.15 | | | 0.13 | | |
| τ_00_ | 0.40 _M2ID_ | | | 0.23 _M2ID_ | | | 0.20 _M2ID_ | | |
| ICC | 0.46 | | | 0.60 | | | 0.60 | | |
| N | 234 _M2ID_ | | | 235 _M2ID_ | | | 234 _M2ID_ | | |
| Observations | 451 | | | 458 | | | 443 | | |
| Marginal R^2^ / Conditional R^2^ | 0.182 / 0.561 | | | 0.305 / 0.720 | | | 0.317 / 0.724 | | |

| **Table S12:** Simple multilevel models to test Hypotheses 3 and 4. Sex, race, and lag in 1 part as covariates. | | | | | | | | | |
| --- | --- | --- | --- | --- | --- | --- | --- | --- | --- |
|  | **Episodic Memory** | | | **Executive Functioning** | | | **BTACT** | | |
| *Predictors* | *Estimates* | *CI* | *p* | *Estimates* | *CI* | *p* | *Estimates* | *CI* | *p* |
| Intercept (M2) | **-0.37** | **-0.57 – -0.17** | **<0.001** | 0.06 | -0.08 – 0.20 | 0.412 | -0.01 | -0.14 – 0.12 | 0.872 |
| MIDUS Wave | **-0.13** | **-0.26 – -0.00** | **0.048** | **-0.44** | **-0.51 – -0.36** | **<0.001** | **-0.38** | **-0.45 – -0.31** | **<0.001** |
| Exponent | 0.29 | -0.16 – 0.74 | 0.201 | **0.45** | **0.14 – 0.76** | **0.005** | **0.43** | **0.14 – 0.73** | **0.004** |
| Sex | **0.71** | **0.49 – 0.93** | **<0.001** | 0.09 | -0.07 – 0.25 | 0.252 | **0.20** | **0.05 – 0.36** | **0.008** |
| Race | -0.19 | -0.51 – 0.13 | 0.250 | **-0.45** | **-0.68 – -0.22** | **<0.001** | **-0.42** | **-0.64 – -0.19** | **<0.001** |
| Lag between Waves | -0.03 | -0.18 – 0.12 | 0.714 | -0.06 | -0.17 – 0.05 | 0.288 | -0.06 | -0.16 – 0.05 | 0.291 |
| Wave x Exponent | -0.02 | -0.48 – 0.43 | 0.917 | -0.22 | -0.48 – 0.04 | 0.101 | -0.17 | -0.42 – 0.08 | 0.175 |
| **Random Effects** | | | | | | | | | |
| σ^2^ | 0.47 | | | 0.15 | | | 0.13 | | |
| τ_00_ | 0.45 _M2ID_ | | | 0.29 _M2ID_ | | | 0.26 _M2ID_ | | |
| ICC | 0.49 | | | 0.66 | | | 0.66 | | |
| N | 234 _M2ID_ | | | 235 _M2ID_ | | | 234 _M2ID_ | | |
| Observations | 451 | | | 458 | | | 443 | | |
| Marginal R^2^ / Conditional R^2^ | 0.123 / 0.553 | | | 0.200 / 0.725 | | | 0.198 / 0.726 | | |

| **Table S13:** Simple multilevel models to test Hypotheses 5 and 6. Sex, race, and lag in 1 part as covariates. | | | | | | | | | |
| --- | --- | --- | --- | --- | --- | --- | --- | --- | --- |
|  | **Episodic Memory** | | | **Executive Functioning** | | | **BTACT** | | |
| *Predictors* | *Estimates* | *CI* | *p* | *Estimates* | *CI* | *p* | *Estimates* | *CI* | *p* |
| Intercept (M2) | **-0.36** | **-0.55 – -0.16** | **<0.001** | 0.07 | -0.07 – 0.21 | 0.338 | 0.00 | -0.13 – 0.13 | 0.995 |
| MIDUS Wave | **-0.13** | **-0.26 – -0.00** | **0.049** | **-0.44** | **-0.51 – -0.36** | **<0.001** | **-0.38** | **-0.45 – -0.32** | **<0.001** |
| Individual Alpha Peak Frequency | **0.13** | **0.00 – 0.26** | **0.046** | 0.07 | -0.02 – 0.16 | 0.109 | 0.08 | -0.00 – 0.16 | 0.060 |
| Sex | **0.68** | **0.46 – 0.89** | **<0.001** | 0.06 | -0.09 – 0.22 | 0.421 | **0.17** | **0.02 – 0.32** | **0.025** |
| Race | -0.15 | -0.47 – 0.16 | 0.341 | **-0.43** | **-0.66 – -0.19** | **<0.001** | **-0.38** | **-0.61 – -0.16** | **0.001** |
| Lag between Waves | -0.02 | -0.17 – 0.13 | 0.828 | -0.05 | -0.16 – 0.06 | 0.398 | -0.04 | -0.15 – 0.06 | 0.415 |
| Wave x Alpha Freq | 0.06 | -0.07 – 0.19 | 0.338 | 0.06 | -0.01 – 0.14 | 0.091 | 0.07 | -0.00 – 0.14 | 0.063 |
| **Random Effects** | | | | | | | | | |
| σ^2^ | 0.47 | | | 0.15 | | | 0.13 | | |
| τ_00_ | 0.43 _M2ID_ | | | 0.29 _M2ID_ | | | 0.25 _M2ID_ | | |
| ICC | 0.48 | | | 0.66 | | | 0.66 | | |
| N | 234 _M2ID_ | | | 235 _M2ID_ | | | 234 _M2ID_ | | |
| Observations | 451 | | | 458 | | | 443 | | |
| Marginal R^2^ / Conditional R^2^ | 0.142 / 0.554 | | | 0.202 / 0.725 | | | 0.204 / 0.730 | | |

| **Table S14:** Simple multilevel models to test Hypotheses 7a and 7c. Sex, race, and lag in 1 part as covariates. | | | | | | | | | |
| --- | --- | --- | --- | --- | --- | --- | --- | --- | --- |
|  | **Episodic Memory** | | | **Executive Functioning** | | | **BTACT** | | |
| *Predictors* | *Estimates* | *CI* | *p* | *Estimates* | *CI* | *p* | *Estimates* | *CI* | *p* |
| Intercept (M2) | **-0.40** | **-0.59 – -0.20** | **<0.001** | 0.05 | -0.09 – 0.18 | 0.488 | -0.02 | -0.15 – 0.10 | 0.721 |
| MIDUS Wave | -0.11 | -0.24 – 0.03 | 0.118 | **-0.43** | **-0.50 – -0.35** | **<0.001** | **-0.38** | **-0.45 – -0.31** | **<0.001** |
| Exponent | 0.11 | -0.35 – 0.56 | 0.650 | 0.20 | -0.10 – 0.50 | 0.181 | 0.20 | -0.08 – 0.48 | 0.171 |
| Age | **-0.02** | **-0.03 – -0.01** | **0.003** | **-0.02** | **-0.03 – -0.01** | **<0.001** | **-0.02** | **-0.03 – -0.01** | **<0.001** |
| Sex | **0.73** | **0.52 – 0.95** | **<0.001** | 0.12 | -0.03 – 0.26 | 0.116 | **0.23** | **0.09 – 0.37** | **0.001** |
| Race | -0.18 | -0.49 – 0.13 | 0.266 | **-0.43** | **-0.65 – -0.22** | **<0.001** | **-0.40** | **-0.60 – -0.20** | **<0.001** |
| Lag between Waves | -0.05 | -0.20 – 0.09 | 0.474 | -0.09 | -0.19 – 0.01 | 0.093 | -0.08 | -0.18 – 0.01 | 0.085 |
| Wave X Exponent | -0.15 | -0.62 – 0.33 | 0.539 | -0.24 | -0.51 – 0.03 | 0.087 | -0.20 | -0.46 – 0.06 | 0.139 |
| Wave X Age | -0.01 | -0.02 – 0.00 | 0.112 | -0.00 | -0.01 – 0.01 | 0.683 | -0.00 | -0.01 – 0.00 | 0.466 |
| Exponent X Age | -0.01 | -0.05 – 0.04 | 0.745 | 0.01 | -0.02 – 0.04 | 0.495 | 0.01 | -0.02 – 0.04 | 0.556 |
| Wave X Exponent X Age | 0.03 | -0.02 – 0.08 | 0.187 | 0.01 | -0.02 – 0.03 | 0.690 | 0.00 | -0.02 – 0.03 | 0.774 |
| **Random Effects** | | | | | | | | | |
| σ^2^ | 0.46 | | | 0.15 | | | 0.13 | | |
| τ_00_ | 0.41 _M2ID_ | | | 0.23 _M2ID_ | | | 0.20 _M2ID_ | | |
| ICC | 0.47 | | | 0.60 | | | 0.60 | | |
| N | 234 _M2ID_ | | | 235 _M2ID_ | | | 234 _M2ID_ | | |
| Observations | 451 | | | 458 | | | 443 | | |
| Marginal R^2^ / Conditional R^2^ | 0.183 / 0.565 | | | 0.309 / 0.723 | | | 0.319 / 0.726 | | |

| **Table S15:** Simple multilevel models to test Hypotheses 7b and 7d. Sex, race, and lag in 1 part as covariates. | | | | | | | | | | | | |
| --- | --- | --- | --- | --- | --- | --- | --- | --- | --- | --- | --- | --- |
|  | **Episodic Memory** | | | | | **Executive Functioning** | | | | **BTACT** | | |
| *Predictors* | *Estimates* | | *CI* | *p* | | *Estimates* | *CI* | *p* | | *Estimates* | *CI* | *p* |
| Intercept (M2) | **-0.38** | | **-0.57 – -0.19** | **<0.001** | | 0.05 | -0.08 – 0.18 | 0.476 | | -0.02 | -0.14 – 0.10 | 0.744 |
| MIDUS Wave | -0.13 | | -0.26 – 0.00 | 0.054 | | **-0.44** | **-0.52 – -0.37** | **<0.001** | | **-0.39** | **-0.46 – -0.32** | **<0.001** |
| Alpha Freq | 0.09 | | -0.04 – 0.21 | 0.194 | | 0.01 | -0.07 – 0.10 | 0.763 | | 0.02 | -0.06 – 0.10 | 0.603 |
| Age | **-0.02** | | **-0.03 – -0.01** | **0.005** | | **-0.02** | **-0.03 – -0.02** | **<0.001** | | **-0.02** | **-0.03 – -0.02** | **<0.001** |
| Sex | **0.72** | | **0.51 – 0.93** | **<0.001** | | 0.12 | -0.03 – 0.26 | 0.121 | | **0.22** | **0.09 – 0.36** | **0.002** |
| Race | -0.16 | | -0.47 – 0.15 | 0.307 | | **-0.42** | **-0.64 – -0.21** | **<0.001** | | **-0.39** | **-0.59 – -0.18** | **<0.001** |
| Lag between Waves | -0.05 | | -0.19 – 0.10 | 0.534 | | -0.08 | -0.19 – 0.02 | 0.116 | | -0.08 | -0.17 – 0.02 | 0.111 |
| Wave x Alpha Freq | 0.04 | | -0.10 – 0.17 | 0.564 | | 0.06 | -0.01 – 0.14 | 0.106 | | 0.06 | -0.01 – 0.13 | 0.100 |
| Wave X Age | -0.01 | | -0.02 – 0.00 | 0.146 | | 0.00 | -0.01 – 0.01 | 0.774 | | -0.00 | -0.01 – 0.01 | 0.968 |
| Alpha Freq X Age | 0.00 | | -0.01 – 0.01 | 0.744 | | 0.00 | -0.00 – 0.01 | 0.257 | | 0.00 | -0.00 – 0.01 | 0.288 |
| Wave X Alpha Freq X Age | 0.00 | | -0.01 – 0.01 | 0.870 | | -0.00 | -0.01 – 0.00 | 0.303 | | -0.00 | -0.01 – 0.00 | 0.409 |
| **Random Effects** | | | | | | | | | | | | |
| σ^2^ | 0.47 | | | | | 0.15 | | | | 0.13 | | |
| τ_00_ | 0.39 _M2ID_ | | | | | 0.23 _M2ID_ | | | | 0.20 _M2ID_ | | |
| ICC | 0.46 | | | | | 0.60 | | | | 0.60 | | |
| N | 234 _M2ID_ | | | | | 235 _M2ID_ | | | | 234 _M2ID_ | | |
| Observations | 451 | | | | | 458 | | | | 443 | | |
| Marginal R^2^ / Conditional R^2^ | | 0.191 / 0.560 | | | 0.310 / 0.725 | | | | 0.323 / 0.729 | | | |

| **Table S16:** Simple multilevel models exploratory analyses. Sex, race, and lag in 1 part as covariates. | | | | | | | | | |
| --- | --- | --- | --- | --- | --- | --- | --- | --- | --- |
|  | **Episodic Memory** | | | **Executive Functioning** | | | **BTACT** | | |
| *Predictors* | *Estimates* | *CI* | *p* | *Estimates* | *CI* | *p* | *Estimates* | *CI* | *p* |
| Intercept (M2) | **-0.39** | **-0.58 – -0.20** | **<0.001** | 0.03 | -0.10 – 0.16 | 0.677 | -0.04 | -0.16 – 0.08 | 0.526 |
| MIDUS Wave | -0.12 | -0.25 – 0.01 | 0.061 | **-0.42** | **-0.49 – -0.35** | **<0.001** | **-0.37** | **-0.44 – -0.30** | **<0.001** |
| Exponent | 0.09 | -0.39 – 0.57 | 0.706 | 0.18 | -0.14 – 0.50 | 0.267 | 0.17 | -0.12 – 0.47 | 0.254 |
| Alpha Peak | 0.08 | -0.05 – 0.21 | 0.229 | 0.02 | -0.06 – 0.11 | 0.571 | 0.03 | -0.05 – 0.11 | 0.449 |
| Age | **-0.02** | **-0.03 – -0.01** | **<0.001** | **-0.02** | **-0.03 – -0.02** | **<0.001** | **-0.02** | **-0.03 – -0.02** | **<0.001** |
| Sex | **0.73** | **0.51 – 0.94** | **<0.001** | 0.12 | -0.03 – 0.27 | 0.108 | **0.23** | **0.09 – 0.37** | **0.001** |
| Race | -0.16 | -0.47 – 0.15 | 0.301 | **-0.43** | **-0.64 – -0.21** | **<0.001** | **-0.39** | **-0.59 – -0.19** | **<0.001** |
| Lag between Waves | -0.05 | -0.20 – 0.10 | 0.496 | -0.09 | -0.19 – 0.01 | 0.090 | -0.08 | -0.18 – 0.01 | 0.085 |
| Wave X Exponent | 0.07 | -0.42 – 0.55 | 0.781 | -0.11 | -0.37 – 0.16 | 0.438 | -0.05 | -0.31 – 0.20 | 0.692 |
| Wave X Alpha | 0.06 | -0.07 – 0.19 | 0.369 | 0.05 | -0.03 – 0.12 | 0.206 | 0.05 | -0.02 – 0.12 | 0.152 |
| Exponent X Alpha | -0.08 | -0.53 – 0.37 | 0.737 | -0.16 | -0.45 – 0.14 | 0.303 | -0.15 | -0.43 – 0.13 | 0.288 |
| Wave X Exponent X Alpha | 0.15 | -0.31 – 0.62 | 0.514 | **0.31** | **0.04 – 0.57** | **0.024** | **0.29** | **0.04 – 0.54** | **0.021** |
| **Random Effects** | | | | | | | | | |
| σ^2^ | 0.47 | | | 0.15 | | | 0.13 | | |
| τ_00_ | 0.39 _M2ID_ | | | 0.23 _M2ID_ | | | 0.20 _M2ID_ | | |
| ICC | 0.45 | | | 0.61 | | | 0.61 | | |
| N | 234 _M2ID_ | | | 235 _M2ID_ | | | 234 _M2ID_ | | |
| Observations | 451 | | | 458 | | | 443 | | |
| Marginal R^2^ / Conditional R^2^ | 0.189 / 0.557 | | | 0.313 / 0.734 | | | 0.327 / 0.737 | | |

| **Table S17:** Examining the Wave X Aperiodic Exponent X Individual Peak Alpha Frequency interaction from analyses described in Table S16 (simple multilevel model, Sex, race, and lag in 1 part as covariates) through the slope of the change in cognitive function over waves for each BTACT measure at high and low levels of each EEG metric. | | | | | | |
| --- | --- | --- | --- | --- | --- | --- |
| **Change in BTACT Episodic Memory Factor from M2 to M3** | | | | | | |
|  | *Slope of IAPF* | | *95% CI* | | *p-value* | |
| Low Exponent (-1 SD) | 0.02 | | -0.17 – 0.21 | | 0.858 | |
| High Exponent (+1 SD) | 0.10 | | -0.07 – 0.28 | | 0.253 | |
|  | *Slope of Exponent* | | *95% CI* | | *p-value* | |
| Low IAPF (-1 SD) | -0.08 | | -0.64 –0.48 | | 0.771 | |
| High IAPF (+1 SD) | 0.22 | | -0.54 – 0.98 | | 0.568 | |
| **Change in BTACT Executive Functioning Factor from M2 to M3** | | | | | | |
|  | | *Slope of IAPF* | | *95% CI* | | *p-value* |
| Low Exponent (-1 SD) | | -0.04 | | -0.15 – 0.07 | | 0.501 |
| High Exponent (+1 SD) | | **0.13** | | **0.03 – 0.23** | | **0.009** |
|  | | *Slope of Exponent* | | *95% CI* | | *p-value* |
| Low IAPF (-1 SD) | | **-0.40** | | **-0.73 – -0.08** | | **0.014** |
| High IAPF (+1 SD) | | 0.19 | | -0.22 – 0.61 | | 0.362 |
| **Change in BTACT Overall Composite from M2 to M3** | | | | | | |
|  | | *Slope of IAPF* | | *95% CI* | | *p-value* |
| Low Exponent (-1 SD) | | -0.03 | | -0.13 – 0.07 | | 0.559 |
| High Exponent (+1 SD) | | **0.13** | | **0.04 – 0.23** | | **0.006** |
|  | | *Slope of Exponent* | | *95% CI* | | *p-value* |
| Low IAPF (-1 SD) | | **-0.34** | | **-0.65 – -0.03** | | **0.030** |
| High IAPF (+1 SD) | | 0.24 | | -0.16 – 0.63 | | 0.242 |

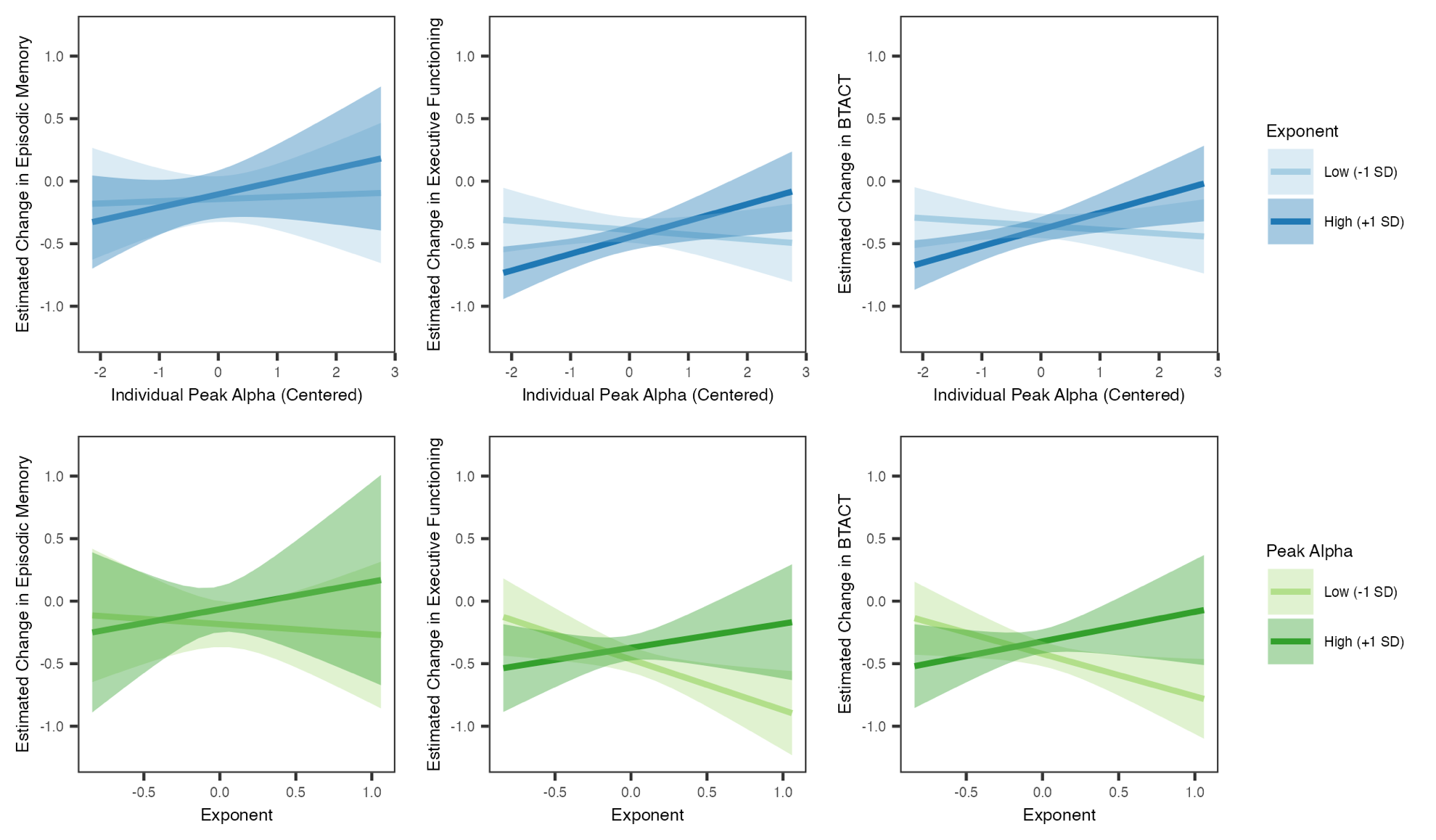

Figure S2. Wave by aperiodic exponent by individual peak alpha frequency interaction plot for models described in Table S16 (simple multilevel model, Sex, race, and lag in 1 part as covariates).

**Analyses Parallel to Preregistration: Complex analyses with sex, education, and lag split into 2 parts**

| **Table S18:** Complex multilevel models to test Hypotheses 1 and 2. Sex, education, and lag in 2 parts as covariates. | | | | | | | | | |
| --- | --- | --- | --- | --- | --- | --- | --- | --- | --- |
|  | **Episodic Memory** | | | **Executive Functioning** | | | **BTACT** | | |
| *Predictors* | *Estimates* | *CI* | *p* | *Estimates* | *CI* | *p* | *Estimates* | *CI* | *p* |
| Intercept (M2) | **-0.37** | **-0.56 – -0.18** | **<0.001** | -0.09 | -0.22 – 0.03 | 0.141 | **-0.13** | **-0.25 – -0.02** | **0.025** |
| MIDUS Wave | **-0.24** | **-0.44 – -0.04** | **0.018** | **-0.37** | **-0.48 – -0.26** | **<0.001** | **-0.35** | **-0.46 – -0.24** | **<0.001** |
| Age | **-0.02** | **-0.03 – -0.01** | **0.005** | **-0.02** | **-0.03 – -0.01** | **<0.001** | **-0.02** | **-0.03 – -0.01** | **<0.001** |
| Sex | **0.64** | **0.40 – 0.89** | **<0.001** | 0.16 | -0.00 – 0.32 | 0.054 | **0.24** | **0.09 – 0.39** | **0.002** |
| Education | **0.16** | **0.01 – 0.31** | **0.035** | **0.16** | **0.07 – 0.26** | **0.001** | **0.15** | **0.07 – 0.24** | **0.001** |
| M2 Cog to Neuro Lag | 0.03 | -0.15 – 0.21 | 0.742 | -0.05 | -0.17 – 0.07 | 0.401 | -0.04 | -0.15 – 0.08 | 0.533 |
| Neuro to M3 Cog Lag | -0.06 | -0.20 – 0.07 | 0.367 | **-0.13** | **-0.21 – -0.04** | **0.005** | **-0.11** | **-0.20 – -0.03** | **0.007** |
| Wave X Age | **-0.01** | **-0.03 – -0.00** | **0.038** | -0.00 | -0.01 – 0.01 | 0.634 | -0.00 | -0.01 – 0.00 | 0.319 |
| Wave X Sex | 0.18 | -0.08 – 0.44 | 0.165 | -0.10 | -0.25 – 0.04 | 0.171 | -0.05 | -0.20 – 0.09 | 0.454 |
| Wave X Education | -0.01 | -0.19 – 0.18 | 0.955 | -0.06 | -0.16 – 0.05 | 0.282 | -0.04 | -0.14 – 0.06 | 0.419 |
| Wave X M2 Cog to Neuro Lag | -0.14 | -0.34 – 0.05 | 0.140 | **-0.15** | **-0.26 – -0.04** | **0.006** | **-0.17** | **-0.27 – -0.06** | **0.002** |
| Wave X Neuro to M3 Cog Lag | -0.05 | -0.19 – 0.09 | 0.464 | **-0.14** | **-0.22 – -0.06** | **0.001** | **-0.13** | **-0.21 – -0.06** | **0.001** |
| **Random Effects** | | | | | | | | | |
| σ^2^ | 0.46 | | | 0.15 | | | 0.13 | | |
| τ_00_ | 0.38 _M2ID_ | | | 0.22 _M2ID_ | | | 0.19 _M2ID_ | | |
| ICC | 0.45 | | | 0.60 | | | 0.58 | | |
| N | 234 _M2ID_ | | | 235 _M2ID_ | | | 234 _M2ID_ | | |
| Observations | 451 | | | 458 | | | 443 | | |
| Marginal R^2^ / Conditional R^2^ | 0.202 / 0.564 | | | 0.312 / 0.723 | | | 0.330 / 0.722 | | |

| **Table S19:** Complex multilevel models to test Hypotheses 3 and 4. Sex, education, and lag in 2 parts as covariates. | | | | | | | | | |
| --- | --- | --- | --- | --- | --- | --- | --- | --- | --- |
|  | **Episodic Memory** | | | **Executive Functioning** | | | **BTACT** | | |
| *Predictors* | *Estimates* | *CI* | *p* | *Estimates* | *CI* | *p* | *Estimates* | *CI* | *p* |
| Intercept (M2) | **-0.36** | **-0.56 – -0.16** | **<0.001** | -0.08 | -0.21 – 0.05 | 0.246 | -0.12 | -0.25 – 0.01 | 0.063 |
| MIDUS Wave | **-0.23** | **-0.43 – -0.02** | **0.029** | **-0.37** | **-0.48 – -0.26** | **<0.001** | **-0.35** | **-0.46 – -0.24** | **<0.001** |
| Exponent | 0.21 | -0.24 – 0.65 | 0.366 | **0.35** | **0.04 – 0.65** | **0.027** | **0.33** | **0.04 – 0.62** | **0.024** |
| Sex | **0.62** | **0.37 – 0.88** | **<0.001** | 0.14 | -0.04 – 0.31 | 0.123 | **0.22** | **0.05 – 0.38** | **0.009** |
| Education | **0.16** | **0.01 – 0.32** | **0.042** | **0.15** | **0.06 – 0.25** | **0.002** | **0.14** | **0.05 – 0.24** | **0.002** |
| M2 Cog to Neuro Lag | 0.07 | -0.11 – 0.26 | 0.439 | 0.00 | -0.12 – 0.13 | 0.968 | 0.02 | -0.10 – 0.14 | 0.789 |
| Neuro to M3 Cog Lag | -0.04 | -0.18 – 0.10 | 0.562 | **-0.10** | **-0.20 – -0.01** | **0.034** | **-0.09** | **-0.18 – -0.00** | **0.044** |
| Wave X Exponent | 0.04 | -0.43 – 0.51 | 0.872 | -0.16 | -0.42 – 0.10 | 0.224 | -0.11 | -0.36 – 0.14 | 0.396 |
| Wave X Sex | 0.17 | -0.10 – 0.43 | 0.217 | -0.11 | -0.26 – 0.03 | 0.131 | -0.06 | -0.21 – 0.08 | 0.387 |
| Wave X Education | 0.00 | -0.18 – 0.19 | 0.964 | -0.05 | -0.16 – 0.05 | 0.321 | -0.04 | -0.14 – 0.06 | 0.482 |
| Wave X M2 Cog to Neuro Lag | -0.10 | -0.29 – 0.09 | 0.313 | **-0.14** | **-0.25 – -0.03** | **0.011** | **-0.14** | **-0.25 – -0.04** | **0.007** |
| Wave X Neuro to M3 Cog Lag | -0.03 | -0.17 – 0.11 | 0.683 | **-0.13** | **-0.21 – -0.05** | **0.002** | **-0.12** | **-0.20 – -0.04** | **0.003** |
| **Random Effects** | | | | | | | | | |
| σ^2^ | 0.47 | | | 0.15 | | | 0.13 | | |
| τ_00_ | 0.43 _M2ID_ | | | 0.27 _M2ID_ | | | 0.24 _M2ID_ | | |
| ICC | 0.48 | | | 0.65 | | | 0.64 | | |
| N | 234 _M2ID_ | | | 235 _M2ID_ | | | 234 _M2ID_ | | |
| Observations | 451 | | | 458 | | | 443 | | |
| Marginal R^2^ / Conditional R^2^ | 0.147 / 0.552 | | | 0.214 / 0.724 | | | 0.218 / 0.720 | | |

| **Table S20:** Complex multilevel models to test Hypotheses 5 and 6. Sex, education, and lag in 2 parts as covariates. | | | | | | | | |  |
| --- | --- | --- | --- | --- | --- | --- | --- | --- | --- |
| **Episodic Memory** | | | **Executive Functioning** | | | **BTACT** | | |  |
| *Predictors* | *Estimates* | *CI* | *p* | *Estimates* | *CI* | *p* | *Estimates* | *CI* | *p* |
| Intercept (M2) | **-0.35** | **-0.54 – -0.15** | **<0.001** | -0.06 | -0.20 – 0.07 | 0.338 | -0.10 | -0.23 – 0.02 | 0.105 |
| MIDUS Wave | **-0.22** | **-0.43 – -0.02** | **0.032** | **-0.37** | **-0.48 – -0.26** | **<0.001** | **-0.35** | **-0.46 – -0.24** | **<0.001** |
| Alpha Peak | 0.12 | -0.00 – 0.25 | 0.054 | 0.07 | -0.02 – 0.15 | 0.124 | 0.08 | -0.00 – 0.16 | 0.060 |
| Sex | **0.60** | **0.35 – 0.85** | **<0.001** | 0.11 | -0.06 – 0.28 | 0.197 | **0.19** | **0.03 – 0.35** | **0.020** |
| Education | **0.16** | **0.01 – 0.32** | **0.041** | **0.16** | **0.07 – 0.26** | **0.001** | **0.15** | **0.06 – 0.24** | **0.001** |
| M2 Cog to Neuro Lag | 0.09 | -0.09 – 0.27 | 0.320 | 0.03 | -0.10 – 0.15 | 0.672 | 0.04 | -0.08 – 0.16 | 0.486 |
| Neuro to M3 Cog Lag | -0.02 | -0.16 – 0.11 | 0.730 | -0.08 | -0.17 – 0.01 | 0.084 | -0.07 | -0.16 – 0.02 | 0.115 |
| Wave X Alpha Peak | 0.07 | -0.06 – 0.20 | 0.301 | 0.07 | -0.00 – 0.14 | 0.060 | 0.07 | -0.00 – 0.14 | 0.050 |
| Wave X Sex | 0.16 | -0.10 – 0.42 | 0.234 | -0.11 | -0.26 – 0.03 | 0.133 | -0.06 | -0.20 – 0.08 | 0.380 |
| Wave X Education | 0.01 | -0.18 – 0.19 | 0.946 | -0.05 | -0.16 – 0.05 | 0.292 | -0.04 | -0.14 – 0.06 | 0.450 |
| Wave X M2 Cog to Neuro Lag | -0.09 | -0.28 – 0.10 | 0.338 | **-0.15** | **-0.25 – -0.04** | **0.007** | **-0.15** | **-0.25 – -0.05** | **0.005** |
| Wave X Neuro to M3 Cog Lag | -0.02 | -0.16 – 0.12 | 0.749 | **-0.13** | **-0.21 – -0.05** | **0.001** | **-0.12** | **-0.20 – -0.04** | **0.002** |
| **Random Effects** | | | | | | | | | |
| σ^2^ | 0.47 | | | 0.15 | | | 0.13 | | |
| τ_00_ | 0.41 _M2ID_ | | | 0.27 _M2ID_ | | | 0.23 _M2ID_ | | |
| ICC | 0.46 | | | 0.65 | | | 0.64 | | |
| N | 234 _M2ID_ | | | 235 _M2ID_ | | | 234 _M2ID_ | | |
| Observations | 451 | | | 458 | | | 443 | | |
| Marginal R^2^ / Conditional R^2^ | 0.168 / 0.554 | | | 0.225 / 0.727 | | | 0.234 / 0.726 | | |

| **Table S21:** Complex multilevel models to test Hypotheses 7a and 7c. Sex, education, and lag in 2 parts as covariates. | | | | | | | | | |
| --- | --- | --- | --- | --- | --- | --- | --- | --- | --- |
|  | **Episodic Memory** | | | **Executive Functioning** | | | **BTACT** | | |
| *Predictors* | *Estimates* | *CI* | *p* | *Estimates* | *CI* | *p* | *Estimates* | *CI* | *p* |
| Intercept (M2) | **-0.38** | **-0.58 – -0.18** | **<0.001** | -0.09 | -0.22 – 0.04 | 0.184 | **-0.13** | **-0.25 – -0.01** | **0.038** |
| MIDUS Wave | -0.21 | -0.42 – 0.00 | 0.050 | **-0.36** | **-0.48 – -0.24** | **<0.001** | **-0.35** | **-0.46 – -0.23** | **<0.001** |
| Exponent | 0.05 | -0.41 – 0.50 | 0.842 | 0.13 | -0.16 – 0.43 | 0.376 | 0.12 | -0.15 – 0.40 | 0.377 |
| Age | **-0.02** | **-0.03 – -0.00** | **0.007** | **-0.02** | **-0.03 – -0.01** | **<0.001** | **-0.02** | **-0.03 – -0.01** | **<0.001** |
| Sex | **0.65** | **0.40 – 0.90** | **<0.001** | 0.16 | -0.00 – 0.32 | 0.053 | **0.24** | **0.09 – 0.40** | **0.002** |
| Education | **0.16** | **0.01 – 0.31** | **0.037** | **0.16** | **0.06 – 0.25** | **0.001** | **0.15** | **0.06 – 0.24** | **0.001** |
| M2 Cog to Neuro Lag | 0.03 | -0.16 – 0.21 | 0.767 | -0.06 | -0.18 – 0.07 | 0.367 | -0.04 | -0.15 – 0.07 | 0.488 |
| Neuro to M3 Cog Lag | -0.06 | -0.20 – 0.07 | 0.357 | **-0.13** | **-0.22 – -0.04** | **0.004** | **-0.12** | **-0.20 – -0.04** | **0.005** |
| Wave X Exponent | -0.09 | -0.57 – 0.39 | 0.704 | -0.19 | -0.47 – 0.08 | 0.161 | -0.15 | -0.41 – 0.12 | 0.278 |
| Wave X Age | -0.01 | -0.03 – 0.00 | 0.058 | -0.00 | -0.01 – 0.00 | 0.454 | -0.00 | -0.01 – 0.00 | 0.227 |
| Exponent X Age | -0.01 | -0.05 – 0.04 | 0.747 | 0.01 | -0.02 – 0.04 | 0.626 | 0.01 | -0.02 – 0.03 | 0.668 |
| Wave X Sex | 0.17 | -0.10 – 0.43 | 0.215 | -0.11 | -0.26 – 0.04 | 0.136 | -0.06 | -0.20 – 0.08 | 0.396 |
| Wave X Education | -0.01 | -0.19 – 0.17 | 0.935 | -0.05 | -0.16 – 0.05 | 0.307 | -0.04 | -0.14 – 0.06 | 0.450 |
| Wave X M2 Cog to Neuro Lag | -0.14 | -0.33 – 0.06 | 0.169 | **-0.15** | **-0.26 – -0.04** | **0.010** | **-0.16** | **-0.27 – -0.05** | **0.003** |
| Wave X Neuro to M3 Cog Lag | -0.05 | -0.19 – 0.10 | 0.530 | **-0.13** | **-0.21 – -0.05** | **0.002** | **-0.13** | **-0.20 – -0.05** | **0.002** |
| Wave X Exponent X Age | 0.03 | -0.02 – 0.08 | 0.263 | 0.00 | -0.02 – 0.03 | 0.730 | 0.00 | -0.03 – 0.03 | 0.950 |
| **Random Effects** | | | | | | | | | |
| σ^2^ | 0.46 | | | 0.15 | | | 0.13 | | |
| τ_00_ | 0.39 _M2ID_ | | | 0.22 _M2ID_ | | | 0.19 _M2ID_ | | |
| ICC | 0.46 | | | 0.60 | | | 0.59 | | |
| N | 234 _M2ID_ | | | 235 _M2ID_ | | | 234 _M2ID_ | | |
| Observations | 451 | | | 458 | | | 443 | | |
| Marginal R^2^ / Conditional R^2^ | 0.202 / 0.566 | | | 0.313 / 0.725 | | | 0.330 / 0.723 | | |

| **Table S22:** Complex multilevel models to test Hypotheses 7b and 7d. Sex, education, and lag in 2 parts as covariates. | | | | | | | | | |
| --- | --- | --- | --- | --- | --- | --- | --- | --- | --- |
|  | **Episodic Memory** | | | **Executive Functioning** | | | **BTACT** | | |
| *Predictors* | *Estimates* | *CI* | *p* | *Estimates* | *CI* | *p* | *Estimates* | *CI* | *p* |
| Intercept (M2) | **-0.37** | **-0.56 – -0.18** | **<0.001** | -0.09 | -0.21 – 0.04 | 0.175 | **-0.13** | **-0.24 – -0.01** | **0.035** |
| MIDUS Wave | **-0.23** | **-0.44 – -0.03** | **0.025** | **-0.38** | **-0.49 – -0.26** | **<0.001** | **-0.35** | **-0.47 – -0.24** | **<0.001** |
| Alpha Peak | 0.08 | -0.04 – 0.21 | 0.202 | 0.01 | -0.07 – 0.10 | 0.743 | 0.02 | -0.05 – 0.10 | 0.542 |
| Age | **-0.01** | **-0.03 – -0.00** | **0.015** | **-0.02** | **-0.03 – -0.01** | **<0.001** | **-0.02** | **-0.03 – -0.01** | **<0.001** |
| Sex | **0.63** | **0.38 – 0.87** | **<0.001** | 0.16 | -0.00 – 0.32 | 0.052 | **0.24** | **0.09 – 0.39** | **0.002** |
| Education | **0.16** | **0.01 – 0.31** | **0.038** | **0.16** | **0.07 – 0.26** | **0.001** | **0.16** | **0.07 – 0.25** | **0.001** |
| M2 Cog to Neuro Lag | 0.04 | -0.14 – 0.22 | 0.653 | -0.05 | -0.17 – 0.07 | 0.397 | -0.03 | -0.15 – 0.08 | 0.548 |
| Neuro to M3 Cog Lag | -0.05 | -0.19 – 0.08 | 0.437 | **-0.12** | **-0.21 – -0.03** | **0.007** | **-0.11** | **-0.19 – -0.03** | **0.009** |
| Wave X Alpha Peak | 0.04 | -0.10 – 0.18 | 0.561 | 0.06 | -0.01 – 0.14 | 0.096 | 0.06 | -0.02 – 0.13 | 0.119 |
| Wave X Age | -0.01 | -0.02 – 0.00 | 0.076 | -0.00 | -0.01 – 0.01 | 0.898 | -0.00 | -0.01 – 0.00 | 0.548 |
| Alpha Peak X Age | -0.00 | -0.01 – 0.01 | 0.902 | 0.00 | -0.01 – 0.01 | 0.481 | 0.00 | -0.01 – 0.01 | 0.555 |
| Wave X Sex | 0.18 | -0.08 – 0.44 | 0.173 | -0.12 | -0.26 – 0.03 | 0.122 | -0.07 | -0.21 – 0.08 | 0.362 |
| Wave X Education | -0.01 | -0.19 – 0.18 | 0.952 | -0.05 | -0.16 – 0.05 | 0.312 | -0.04 | -0.14 – 0.06 | 0.458 |
| Wave X M2 Cog to Neuro Lag | -0.14 | -0.33 – 0.05 | 0.155 | **-0.14** | **-0.25 – -0.03** | **0.012** | **-0.15** | **-0.26 – -0.05** | **0.004** |
| Wave X Neuro to M3 Cog Lag | -0.05 | -0.19 – 0.10 | 0.528 | **-0.13** | **-0.21 – -0.05** | **0.001** | **-0.13** | **-0.20 – -0.05** | **0.001** |
| Wave X Alpha Peak X Age | 0.00 | -0.01 – 0.02 | 0.655 | -0.00 | -0.01 – 0.00 | 0.193 | -0.00 | -0.01 – 0.00 | 0.318 |
| **Random Effects** | | | | | | | | | |
| σ^2^ | 0.47 | | | 0.15 | | | 0.13 | | |
| τ_00_ | 0.38 _M2ID_ | | | 0.22 _M2ID_ | | | 0.19 _M2ID_ | | |
| ICC | 0.45 | | | 0.60 | | | 0.59 | | |
| N | 234 _M2ID_ | | | 235 _M2ID_ | | | 234 _M2ID_ | | |
| Observations | 451 | | | 458 | | | 443 | | |
| Marginal R^2^ / Conditional R^2^ | 0.210 / 0.564 | | | 0.318 / 0.729 | | | 0.337 / 0.727 | | |

| **Table S23:** Complex multilevel models, exploratory analyses. Sex, education, and lag in 2 parts as covariates. | | | | | | | | | |
| --- | --- | --- | --- | --- | --- | --- | --- | --- | --- |
|  | **Episodic Memory** | | | **Executive Functioning** | | | **BTACT** | | |
| *Predictors* | *Estimates* | *CI* | *p* | *Estimates* | *CI* | *p* | *Estimates* | *CI* | *p* |
| Intercept (M2) | **-0.37** | **-0.56 – -0.18** | **<0.001** | -0.10 | -0.23 – 0.02 | 0.110 | **-0.14** | **-0.26 – -0.02** | **0.019** |
| MIDUS Wave | **-0.23** | **-0.44 – -0.03** | **0.027** | **-0.35** | **-0.46 – -0.23** | **<0.001** | **-0.33** | **-0.44 – -0.22** | **<0.001** |
| Exponent | 0.11 | -0.37 – 0.59 | 0.656 | 0.12 | -0.20 – 0.44 | 0.448 | 0.12 | -0.17 – 0.42 | 0.411 |
| Alpha Peak | 0.09 | -0.04 – 0.22 | 0.165 | 0.02 | -0.07 – 0.11 | 0.629 | 0.03 | -0.05 – 0.11 | 0.436 |
| Age | **-0.01** | **-0.03 – -0.00** | **0.032** | **-0.02** | **-0.03 – -0.01** | **<0.001** | **-0.02** | **-0.03 – -0.01** | **<0.001** |
| Sex | **0.64** | **0.39 – 0.88** | **<0.001** | **0.17** | **0.00 – 0.33** | **0.044** | **0.25** | **0.09 – 0.40** | **0.002** |
| Education | **0.16** | **0.01 – 0.31** | **0.040** | **0.16** | **0.07 – 0.26** | **0.001** | **0.16** | **0.07 – 0.24** | **0.001** |
| M2 Cog to Neuro Lag | 0.03 | -0.15 – 0.22 | 0.712 | -0.06 | -0.18 – 0.06 | 0.319 | -0.04 | -0.16 – 0.07 | 0.446 |
| Neuro to M3 Cog Lag | -0.06 | -0.20 – 0.08 | 0.404 | **-0.14** | **-0.23 – -0.04** | **0.004** | **-0.12** | **-0.21 – -0.04** | **0.005** |
| Wave X Exponent | -0.01 | -0.52 – 0.51 | 0.984 | -0.07 | -0.35 – 0.22 | 0.640 | -0.02 | -0.30 – 0.25 | 0.872 |
| Wave X Alpha Peak | 0.03 | -0.11 – 0.17 | 0.664 | 0.06 | -0.02 – 0.14 | 0.150 | 0.05 | -0.02 – 0.13 | 0.162 |
| Exponent X Alpha Peak | -0.03 | -0.48 – 0.41 | 0.878 | -0.10 | -0.40 – 0.19 | 0.486 | -0.10 | -0.37 – 0.18 | 0.484 |
| Wave X Age | -0.01 | -0.03 – 0.00 | 0.082 | -0.00 | -0.01 – 0.01 | 0.911 | -0.00 | -0.01 – 0.01 | 0.627 |
| Wave X Sex | 0.18 | -0.09 – 0.44 | 0.187 | -0.12 | -0.27 – 0.02 | 0.103 | -0.07 | -0.21 – 0.07 | 0.328 |
| Wave X Race | 0.00 | -0.18 – 0.19 | 0.971 | -0.05 | -0.15 – 0.06 | 0.387 | -0.03 | -0.13 – 0.07 | 0.557 |
| Wave X M2 Cog to Neuro Lag | -0.13 | -0.32 – 0.07 | 0.207 | **-0.12** | **-0.24 – -0.01** | **0.027** | **-0.14** | **-0.24 – -0.03** | **0.011** |
| Wave X Neuro to M3 Cog Lag | -0.04 | -0.18 – 0.11 | 0.615 | **-0.11** | **-0.19 – -0.03** | **0.007** | **-0.11** | **-0.19 – -0.03** | **0.007** |
| Wave X Exponent X Alpha Peak | 0.15 | -0.33 – 0.63 | 0.535 | **0.27** | **0.00 – 0.54** | **0.048** | **0.27** | **0.01 – 0.52** | **0.042** |
| **Random Effects** | | | | | | | | | |
| σ^2^ | 0.47 | | | 0.15 | | | 0.13 | | |
| τ_00_ | 0.38 _M2ID_ | | | 0.22 _M2ID_ | | | 0.19 _M2ID_ | | |
| ICC | 0.45 | | | 0.61 | | | 0.59 | | |
| N | 234 _M2ID_ | | | 235 _M2ID_ | | | 234 _M2ID_ | | |
| Observations | 451 | | | 458 | | | 443 | | |
| Marginal R^2^ / Conditional R^2^ | 0.210 / 0.564 | | | 0.320 / 0.732 | | | 0.340 / 0.731 | | |

| **Table S24:** Examining the Wave X Aperiodic Exponent X Individual Peak Alpha Frequency interaction from analyses described in Table S23 (complex multilevel model, Sex, education, and lag in 2 parts as covariates) through the slope of the change in cognitive function over waves for each BTACT measure at high and low levels of each EEG metric. | | | | | | |
| --- | --- | --- | --- | --- | --- | --- |
| **Change in BTACT Episodic Memory Factor from M2 to M3** | | | | | | |
|  | *Slope of IAPF* | | *95% CI* | | *p-value* | |
| Low Exponent (-1 SD) | -0.01 | | -0.21 – 0.18 | | 0.911 | |
| High Exponent (+1 SD) | 0.07 | | -0.12 – 0.26 | | 0.450 | |
|  | *Slope of Exponent* | | *95% CI* | | *p-value* | |
| Low IAPF (-1 SD) | -0.15 | | -0.74 –0.44 | | 0.612 | |
| High IAPF (+1 SD) | 0.14 | | -0.65 – 0.93 | | 0.723 | |
| **Change in BTACT Executive Functioning Factor from M2 to M3** | | | | | | |
|  | | *Slope of IAPF* | | *95% CI* | | *p-value* |
| Low Exponent (-1 SD) | | -0.02 | | -0.13 – 0.09 | | 0.742 |
| High Exponent (+1 SD) | | **0.13** | | **0.03 – 0.24** | | **0.014** |
|  | | *Slope of Exponent* | | *95% CI* | | *p-value* |
| Low IAPF (-1 SD) | | -0.34 | | -0.67 – 0.00 | | 0.053 |
| High IAPF (+1 SD) | | 0.20 | | -0.23 – 0.63 | | 0.367 |
| **Change in BTACT Overall Composite from M2 to M3** | | | | | | |
|  | | *Slope of IAPF* | | *95% CI* | | *p-value* |
| Low Exponent (-1 SD) | | -0.02 | | -0.13 – 0.09 | | 0.700 |
| High Exponent (+1 SD) | | **0.13** | | **0.03 – 0.23** | | **0.013** |
|  | | *Slope of Exponent* | | *95% CI* | | *p-value* |
| Low IAPF (-1 SD) | | -0.28 | | -0.61 – 0.04 | | 0.087 |
| High IAPF (+1 SD) | | 0.24 | | -0.18 – 0.65 | | 0.263 |

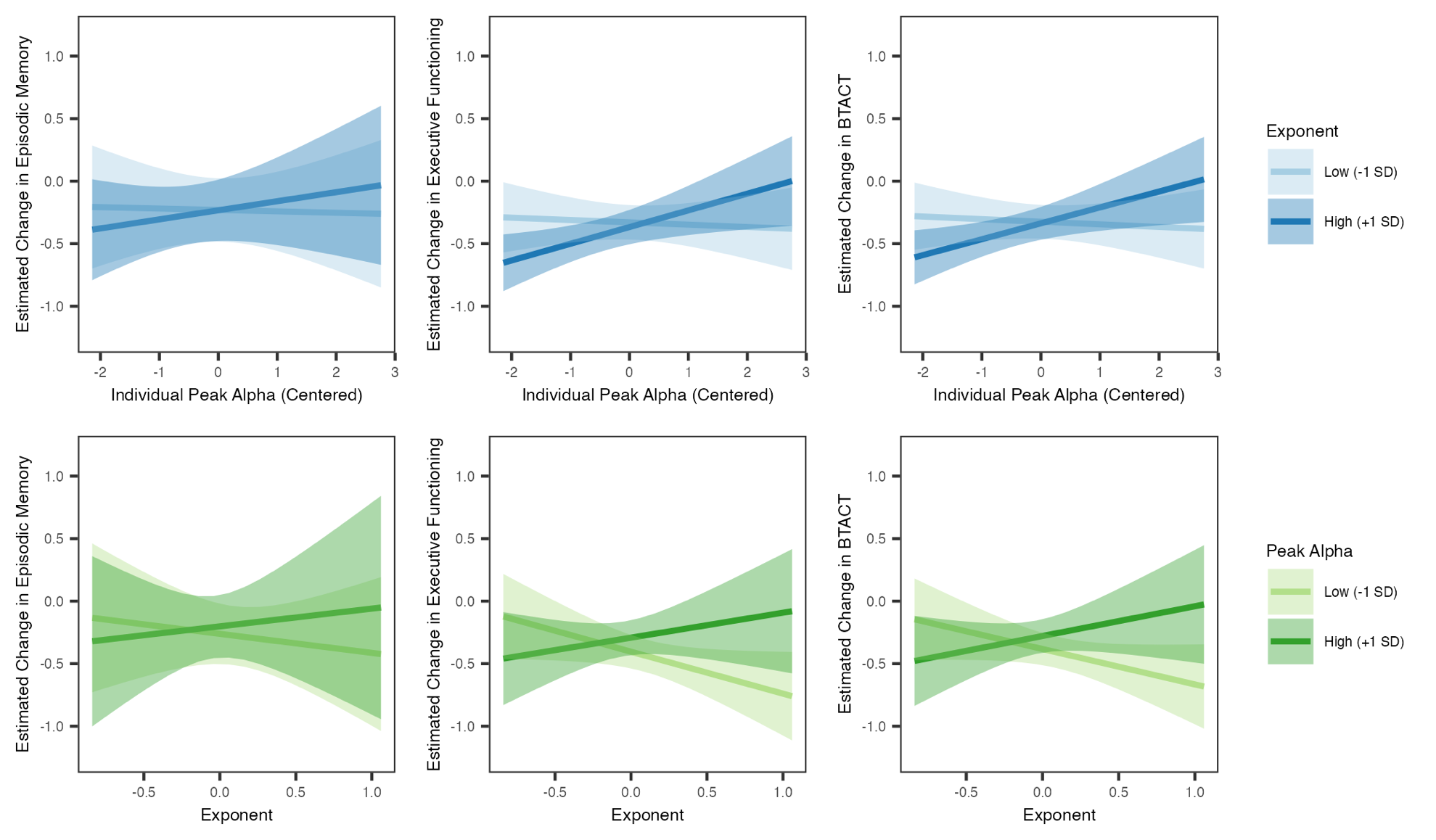

Figure S3. Wave by aperiodic exponent by individual peak alpha frequency interaction plot for models described in Table S23 (complex multilevel model, Sex, education, and lag in 2 parts as covariates).

**Analyses Parallel to Manuscript: Simplified analyses to account for genetic dependencies with family ID and total lag**

| **Table S25:** Simple multilevel models with participants nested within families to test Hypotheses 1 and 2. Lag in 1 part as covariate. | | | | | | | | | |
| --- | --- | --- | --- | --- | --- | --- | --- | --- | --- |
|  | **Episodic Memory** | | | **Executive Functioning** | | | **BTACT** | | |
| *Predictors* | *Estimates* | *CI* | *p* | *Estimates* | *CI* | *p* | *Estimates* | *CI* | *p* |
| Intercept (M2) | 0.01 | -0.12 – 0.14 | 0.914 | -0.01 | -0.09 – 0.08 | 0.890 | 0.00 | -0.07 – 0.08 | 0.943 |
| MIDUS Wave | **-0.14** | **-0.26 – -0.01** | **0.037** | **-0.43** | **-0.50 – -0.36** | **<0.001** | **-0.38** | **-0.45 – -0.31** | **<0.001** |
| Age | **-0.02** | **-0.03 – -0.00** | **0.008** | **-0.02** | **-0.03 – -0.02** | **<0.001** | **-0.02** | **-0.03 – -0.02** | **<0.001** |
| Total Lag | -0.10 | -0.22 – 0.02 | 0.114 | **-0.22** | **-0.30 – -0.14** | **<0.001** | **-0.20** | **-0.28 – -0.13** | **<0.001** |
| Wave X Age | -0.01 | -0.02 – 0.00 | 0.097 | -0.00 | -0.01 – 0.01 | 0.982 | -0.00 | -0.01 – 0.01 | 0.693 |
| **Random Effects** | | | | | | | | | |
| σ^2^ | 0.46 | | | 0.15 | | | 0.13 | | |
| τ_00_ | 0.53 _M2ID:M2FAMNUM_ | | | 0.25 _M2ID:M2FAMNUM_ | | | 0.22 _M2ID:M2FAMNUM_ | | |
| ICC | 0.53 | | | 0.62 | | | 0.63 | | |
| N | 234 _M2ID_ | | | 235 _M2ID_ | | | 234 _M2ID_ | | |
|  | 151 _M2FAMNUM_ | | | 151 _M2FAMNUM_ | | | 151 _M2FAMNUM_ | | |
| Observations | 451 | | | 458 | | | 443 | | |
| Marginal R^2^ / Conditional R^2^ | 0.060 / 0.559 | | | 0.265 / 0.720 | | | 0.261 / 0.723 | | |

| **Table S26:** Simple multilevel models with participants nested within families to test Hypotheses 3 and 4. Lag in 1 part as covariate. | | | | | | | | | |
| --- | --- | --- | --- | --- | --- | --- | --- | --- | --- |
|  | **Episodic Memory** | | | **Executive Functioning** | | | **BTACT** | | |
| *Predictors* | *Estimates* | *CI* | *p* | *Estimates* | *CI* | *p* | *Estimates* | *CI* | *p* |
| Intercept (M2) | 0.01 | -0.12 – 0.14 | 0.910 | -0.00 | -0.09 – 0.08 | 0.921 | 0.00 | -0.08 – 0.09 | 0.929 |
| MIDUS Wave | **-0.13** | **-0.26 – -0.00** | **0.044** | **-0.44** | **-0.51 – -0.36** | **<0.001** | **-0.38** | **-0.45 – -0.31** | **<0.001** |
| Exponent | 0.15 | -0.32 – 0.63 | 0.531 | **0.40** | **0.09 – 0.72** | **0.013** | **0.37** | **0.07 – 0.67** | **0.016** |
| Total Lag | -0.07 | -0.20 – 0.05 | 0.251 | **-0.20** | **-0.29 – -0.11** | **<0.001** | **-0.18** | **-0.26 – -0.10** | **<0.001** |
| Wave X Exponent | -0.03 | -0.49 – 0.43 | 0.893 | -0.21 | -0.47 – 0.05 | 0.106 | -0.17 | -0.42 – 0.08 | 0.176 |
| **Random Effects** | | | | | | | | | |
| σ^2^ | 0.47 | | | 0.15 | | | 0.13 | | |
| τ_00_ | 0.57 _M2ID:M2FAMNUM_ | | | 0.31 _M2ID:M2FAMNUM_ | | | 0.28 _M2ID:M2FAMNUM_ | | |
| ICC | 0.55 | | | 0.67 | | | 0.68 | | |
| N | 234 _M2ID_ | | | 235 _M2ID_ | | | 234 _M2ID_ | | |
|  | 151 _M2FAMNUM_ | | | 151 _M2FAMNUM_ | | | 151 _M2FAMNUM_ | | |
| Observations | 451 | | | 458 | | | 443 | | |
| Marginal R^2^ / Conditional R^2^ | 0.009 / 0.552 | | | 0.157 / 0.724 | | | 0.142 / 0.725 | | |

| **Table S27:** Simple multilevel models with participants nested within families to test Hypotheses 5 and 6. Lag in 1 part as covariate. | | | | | | | | | |
| --- | --- | --- | --- | --- | --- | --- | --- | --- | --- |
|  | **Episodic Memory** | | | **Executive Functioning** | | | **BTACT** | | |
| *Predictors* | *Estimates* | *CI* | *p* | *Estimates* | *CI* | *p* | *Estimates* | *CI* | *p* |
| Intercept (M2) | 0.01 | -0.12 – 0.14 | 0.900 | -0.00 | -0.09 – 0.08 | 0.917 | 0.00 | -0.08 – 0.09 | 0.922 |
| MIDUS Wave | **-0.13** | **-0.26 – -0.00** | **0.045** | **-0.44** | **-0.51 – -0.36** | **<0.001** | **-0.38** | **-0.45 – -0.32** | **<0.001** |
| Alpha Peak | **0.15** | **0.01 – 0.28** | **0.031** | 0.08 | -0.01 – 0.17 | 0.086 | **0.09** | **0.00 – 0.17** | **0.039** |
| Total Lag | -0.06 | -0.18 – 0.07 | 0.367 | **-0.18** | **-0.27 – -0.09** | **<0.001** | **-0.16** | **-0.24 – -0.08** | **<0.001** |
| Wave X Alpha Peak | 0.06 | -0.07 – 0.19 | 0.355 | 0.06 | -0.01 – 0.14 | 0.093 | 0.07 | -0.00 – 0.14 | 0.066 |
| **Random Effects** | | | | | | | | | |
| σ^2^ | 0.47 | | | 0.15 | | | 0.13 | | |
| τ_00_ | 0.54 _M2ID:M2FAMNUM_ | | | 0.30 _M2ID:M2FAMNUM_ | | | 0.27 _M2ID:M2FAMNUM_ | | |
| ICC | 0.54 | | | 0.67 | | | 0.68 | | |
| N | 234 _M2ID_ | | | 235 _M2ID_ | | | 234 _M2ID_ | | |
|  | 151 _M2FAMNUM_ | | | 151 _M2FAMNUM_ | | | 151 _M2FAMNUM_ | | |
| Observations | 451 | | | 458 | | | 443 | | |
| Marginal R^2^ / Conditional R^2^ | 0.038 / 0.554 | | | 0.165 / 0.724 | | | 0.160 / 0.729 | | |

| **Table S28:** Simple multilevel models with participants nested within families to test Hypotheses 7a and 7c. Lag in 1 part as covariate. | | | | | | | | | |
| --- | --- | --- | --- | --- | --- | --- | --- | --- | --- |
|  | **Episodic Memory** | | | **Executive Functioning** | | | **BTACT** | | |
| *Predictors* | *Estimates* | *CI* | *p* | *Estimates* | *CI* | *p* | *Estimates* | *CI* | *p* |
| Intercept (M2) | 0.01 | -0.13 – 0.15 | 0.880 | 0.00 | -0.08 – 0.09 | 0.936 | 0.01 | -0.07 – 0.09 | 0.768 |
| MIDUS Wave | -0.11 | -0.24 – 0.03 | 0.111 | **-0.43** | **-0.50 – -0.35** | **<0.001** | **-0.38** | **-0.45 – -0.31** | **<0.001** |
| Exponent | -0.03 | -0.51 – 0.46 | 0.914 | 0.15 | -0.15 – 0.46 | 0.324 | 0.13 | -0.16 – 0.42 | 0.382 |
| Age | **-0.02** | **-0.03 – -0.00** | **0.011** | **-0.02** | **-0.03 – -0.01** | **<0.001** | **-0.02** | **-0.03 – -0.01** | **<0.001** |
| Total Lag | -0.09 | -0.22 – 0.03 | 0.142 | **-0.22** | **-0.30 – -0.14** | **<0.001** | **-0.20** | **-0.28 – -0.13** | **<0.001** |
| Wave X Exponent | -0.16 | -0.63 – 0.32 | 0.518 | -0.23 | -0.51 – 0.04 | 0.091 | -0.20 | -0.46 – 0.06 | 0.140 |
| Wave X Age | -0.01 | -0.02 – 0.00 | 0.123 | -0.00 | -0.01 – 0.01 | 0.659 | -0.00 | -0.01 – 0.00 | 0.458 |
| Exponent X Age | 0.00 | -0.05 – 0.05 | 0.863 | 0.01 | -0.02 – 0.04 | 0.524 | 0.01 | -0.02 – 0.04 | 0.493 |
| Wave X Exponent X Age | 0.03 | -0.02 – 0.08 | 0.182 | 0.00 | -0.02 – 0.03 | 0.725 | 0.00 | -0.02 – 0.03 | 0.816 |
| **Random Effects** | | | | | | | | | |
| σ^2^ | 0.46 | | | 0.15 | | | 0.13 | | |
| τ_00_ | 0.53 _M2ID:M2FAMNUM_ | | | 0.25 _M2ID:M2FAMNUM_ | | | 0.22 _M2ID:M2FAMNUM_ | | |
| ICC | 0.53 | | | 0.62 | | | 0.63 | | |
| N | 234 _M2ID_ | | | 235 _M2ID_ | | | 234 _M2ID_ | | |
|  | 151 _M2FAMNUM_ | | | 151 _M2FAMNUM_ | | | 151 _M2FAMNUM_ | | |
| Observations | 451 | | | 458 | | | 443 | | |
| Marginal R^2^ / Conditional R^2^ | 0.064 / 0.564 | | | 0.267 / 0.723 | | | 0.263 / 0.725 | | |

| **Table S29:** Simple multilevel models with participants nested within families to test Hypotheses 7b and 7d. Lag in 1 part as covariate. | | | | | | | | | |
| --- | --- | --- | --- | --- | --- | --- | --- | --- | --- |
|  | **Episodic Memory** | | | **Executive Functioning** | | | **BTACT** | | |
| *Predictors* | *Estimates* | *CI* | *p* | *Estimates* | *CI* | *p* | *Estimates* | *CI* | *p* |
| Intercept (M2) | 0.01 | -0.12 – 0.14 | 0.863 | 0.00 | -0.08 – 0.09 | 0.909 | 0.01 | -0.07 – 0.09 | 0.756 |
| MIDUS Wave | -0.13 | -0.26 – 0.00 | 0.051 | **-0.44** | **-0.52 – -0.37** | **<0.001** | **-0.39** | **-0.46 – -0.32** | **<0.001** |
| Alpha | 0.11 | -0.03 – 0.25 | 0.110 | 0.02 | -0.07 – 0.11 | 0.631 | 0.03 | -0.05 – 0.12 | 0.414 |
| Age | **-0.01** | **-0.03 – -0.00** | **0.031** | **-0.02** | **-0.03 – -0.02** | **<0.001** | **-0.02** | **-0.03 – -0.01** | **<0.001** |
| Total Lag | -0.08 | -0.21 – 0.04 | 0.182 | **-0.21** | **-0.29 – -0.13** | **<0.001** | **-0.19** | **-0.27 – -0.12** | **<0.001** |
| Wave X Alpha | 0.04 | -0.10 – 0.17 | 0.582 | 0.06 | -0.01 – 0.14 | 0.111 | 0.06 | -0.01 – 0.13 | 0.104 |
| Wave X Age | -0.01 | -0.02 – 0.00 | 0.157 | 0.00 | -0.01 – 0.01 | 0.803 | -0.00 | -0.01 – 0.01 | 0.958 |
| Alpha X Age | 0.00 | -0.01 – 0.01 | 0.812 | 0.00 | -0.00 – 0.01 | 0.311 | 0.00 | -0.00 – 0.01 | 0.346 |
| Wave X Alpha X Age | 0.00 | -0.01 – 0.01 | 0.879 | -0.00 | -0.01 – 0.00 | 0.302 | -0.00 | -0.01 – 0.00 | 0.416 |
| **Random Effects** | | | | | | | | | |
| σ^2^ | 0.47 | | | 0.15 | | | 0.13 | | |
| τ_00_ | 0.52 _M2ID:M2FAMNUM_ | | | 0.25 _M2ID:M2FAMNUM_ | | | 0.22 _M2ID:M2FAMNUM_ | | |
| ICC | 0.52 | | | 0.62 | | | 0.63 | | |
| N | 234 _M2ID_ | | | 235 _M2ID_ | | | 234 _M2ID_ | | |
|  | 151 _M2FAMNUM_ | | | 151 _M2FAMNUM_ | | | 151 _M2FAMNUM_ | | |
| Observations | 451 | | | 458 | | | 443 | | |
| Marginal R^2^ / Conditional R^2^ | 0.074 / 0.559 | | | 0.271 / 0.725 | | | 0.269 / 0.729 | | |

| **Table S30:** Simple multilevel models with participants nested within families, exploratory analyses. Lag in 1 part as covariate. | | | | | | | | | |
| --- | --- | --- | --- | --- | --- | --- | --- | --- | --- |
|  | **Episodic Memory** | | | **Executive Functioning** | | | **BTACT** | | |
| *Predictors* | *Estimates* | *CI* | *p* | *Estimates* | *CI* | *p* | *Estimates* | *CI* | *p* |
| Intercept (M2) | 0.01 | -0.12 – 0.14 | 0.923 | -0.01 | -0.09 – 0.07 | 0.776 | -0.00 | -0.08 – 0.08 | 0.946 |
| MIDUS Wave | -0.13 | -0.26 – 0.00 | 0.050 | **-0.42** | **-0.49 – -0.35** | **<0.001** | **-0.37** | **-0.44 – -0.30** | **<0.001** |
| Exponent | 0.06 | -0.46 – 0.58 | 0.819 | 0.13 | -0.20 – 0.46 | 0.429 | 0.12 | -0.19 – 0.43 | 0.446 |
| Alpha Peak | 0.12 | -0.03 – 0.26 | 0.107 | 0.03 | -0.06 – 0.12 | 0.509 | 0.04 | -0.04 – 0.13 | 0.330 |
| Age | **-0.01** | **-0.03 – -0.00** | **0.046** | **-0.02** | **-0.03 – -0.01** | **<0.001** | **-0.02** | **-0.03 – -0.01** | **<0.001** |
| Total Lag | -0.09 | -0.21 – 0.04 | 0.182 | **-0.22** | **-0.30 – -0.14** | **<0.001** | **-0.20** | **-0.28 – -0.12** | **<0.001** |
| Wave X Exponent | -0.08 | -0.59 – 0.44 | 0.775 | -0.10 | -0.38 – 0.19 | 0.500 | -0.06 | -0.33 – 0.22 | 0.677 |
| Wave X Alpha Freq | 0.03 | -0.11 – 0.17 | 0.705 | 0.05 | -0.03 – 0.13 | 0.226 | 0.05 | -0.03 – 0.12 | 0.194 |
| Exponent X Alpha Freq | -0.04 | -0.52 – 0.44 | 0.884 | -0.15 | -0.46 – 0.15 | 0.327 | -0.14 | -0.43 – 0.15 | 0.336 |
| Wave X Age | -0.01 | -0.02 – 0.00 | 0.139 | 0.00 | -0.01 – 0.01 | 0.940 | -0.00 | -0.01 – 0.01 | 0.894 |
| Wave X Exponent X Alpha Freq | 0.12 | -0.34 – 0.59 | 0.600 | **0.31** | **0.04 – 0.57** | **0.023** | **0.29** | **0.04 – 0.54** | **0.021** |
| **Random Effects** | | | | | | | | | |
| σ^2^ | 0.47 | | | 0.15 | | | 0.13 | | |
| τ_00_ | 0.52 _M2ID:M2FAMNUM_ | | | 0.25 _M2ID:M2FAMNUM_ | | | 0.23 _M2ID:M2FAMNUM_ | | |
| ICC | 0.52 | | | 0.63 | | | 0.64 | | |
| N | 234 _M2ID_ | | | 235 _M2ID_ | | | 234 _M2ID_ | | |
|  | 151 _M2FAMNUM_ | | | 151 _M2FAMNUM_ | | | 151 _M2FAMNUM_ | | |
| Observations | 451 | | | 458 | | | 443 | | |
| Marginal R^2^ / Conditional R^2^ | 0.074 / 0.560 | | | 0.273 / 0.733 | | | 0.272 / 0.736 | | |

| **Table S31:** Examining the Wave X Aperiodic Exponent X Individual Peak Alpha Frequency interaction from analyses described in Table S30 (simple multilevel model, participant nested within family, lag in 1 part as covariate) through the slope of the change in cognitive function over waves for each BTACT measure at high and low levels of each EEG metric. | | | | | | |
| --- | --- | --- | --- | --- | --- | --- |
| **Change in BTACT Episodic Memory Factor from M2 to M3** | | | | | | |
|  | *Slope of IAPF* | | *95% CI* | | *p-value* | |
| Low Exponent (-1 SD) | -0.01 | | -0.20 – 0.19 | | 0.936 | |
| High Exponent (+1 SD) | 0.06 | | -0.12 – 0.25 | | 0.515 | |
|  | *Slope of Exponent* | | *95% CI* | | *p-value* | |
| Low IAPF (-1 SD) | -0.20 | | -0.77 – 0.38 | | 0.502 | |
| High IAPF (+1 SD) | 0.05 | | -0.74 –0.83 | | 0.907 | |
| **Change in BTACT Executive Functioning Factor from M2 to M3** | | | | | | |
|  | | *Slope of IAPF* | | *95% CI* | | *p-value* |
| Low Exponent (-1 SD) | | -0.04 | | -0.15 – 0.07 | | 0.502 |
| High Exponent (+1 SD) | | **0.13** | | **0.03 – 0.24** | | **0.012** |
|  | | *Slope of Exponent* | | *95% CI* | | *p-value* |
| Low IAPF (-1 SD) | | **-0.40** | | **-0.73– -0.07** | | **0.019** |
| High IAPF (+1 SD) | | 0.20 | | -0.23 – 0.64 | | 0.355 |
| **Change in BTACT Overall Composite from M2 to M3** | | | | | | |
|  | | *Slope of IAPF* | | *95% CI* | | *p-value* |
| Low Exponent (-1 SD) | | -0.03 | | -0.14 – 0.07 | | 0.538 |
| High Exponent (+1 SD) | | **0.13** | | **0.03 – 0.23** | | **0.009** |
|  | | *Slope of Exponent* | | *95% CI* | | *p-value* |
| Low IAPF (-1 SD) | | **-0.35** | | **-0.66 – -0.03** | | **0.032** |
| High IAPF (+1 SD) | | 0.23 | | -0.18 – 0.64 | | 0.274 |

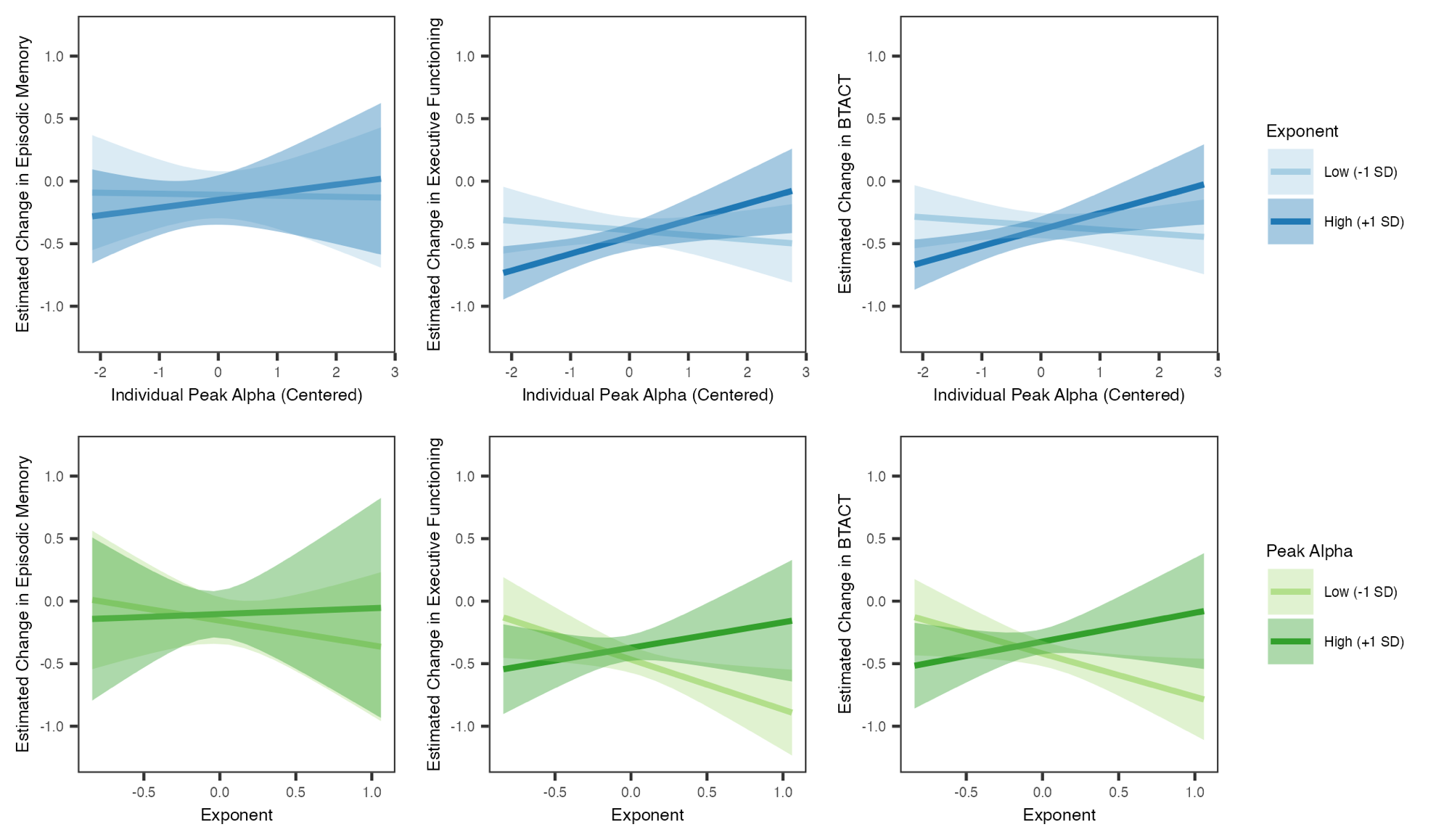

Figure S4. Wave by aperiodic exponent by individual peak alpha frequency interaction plot for models described in Table S30 (simple multilevel model, participant nested within family, lag in 1 part as covariate).

**Analyses Parallel to Preregistration: Complex analyses to account for genetic dependencies with family ID and lag split into 2 parts**

| **Table S32:** Complex multilevel models with participants nested within families to test Hypotheses 1 and 2. Lag in 2 parts as covariates. | | | | | | | | | |
| --- | --- | --- | --- | --- | --- | --- | --- | --- | --- |
|  | **Episodic Memory** | | | **Executive Functioning** | | | **BTACT** | | |
| *Predictors* | *Estimates* | *CI* | *p* | *Estimates* | *CI* | *p* | *Estimates* | *CI* | *p* |
| Intercept (M2) | 0.01 | -0.12 – 0.14 | 0.905 | -0.00 | -0.08 – 0.08 | 0.925 | 0.00 | -0.07 – 0.08 | 0.910 |
| MIDUS Wave | **-0.14** | **-0.26 – -0.01** | **0.036** | **-0.43** | **-0.50 – -0.36** | **<0.001** | **-0.38** | **-0.45 – -0.32** | **<0.001** |
| Age | **-0.01** | **-0.03 – -0.00** | **0.022** | **-0.02** | **-0.03 – -0.01** | **<0.001** | **-0.02** | **-0.03 – -0.01** | **<0.001** |
| M2 Cog to Neuro Lag | 0.04 | -0.15 – 0.24 | 0.659 | -0.05 | -0.18 – 0.07 | 0.404 | -0.03 | -0.15 – 0.08 | 0.590 |
| Neuro to M3 Cog Lag | -0.06 | -0.20 – 0.08 | 0.419 | **-0.13** | **-0.22 – -0.04** | **0.004** | **-0.12** | **-0.20 – -0.03** | **0.007** |
| Wave X Age | **-0.01** | **-0.02 – -0.00** | **0.048** | -0.00 | -0.01 – 0.00 | 0.558 | -0.00 | -0.01 – 0.00 | 0.293 |
| Wave X M2 Cog to Neuro Lag | -0.14 | -0.33 – 0.05 | 0.151 | **-0.15** | **-0.26 – -0.04** | **0.007** | **-0.17** | **-0.27 – -0.06** | **0.002** |
| Wave X Neuro to M3 Cog Lag | -0.05 | -0.19 – 0.09 | 0.513 | **-0.13** | **-0.21 – -0.05** | **0.001** | **-0.13** | **-0.20 – -0.05** | **0.001** |
| **Random Effects** | | | | | | | | | |
| σ^2^ | 0.46 | | | 0.15 | | | 0.13 | | |
| τ_00_ | 0.52 _M2ID:M2FAMNUM_ | | | 0.24 _M2ID:M2FAMNUM_ | | | 0.22 _M2ID:M2FAMNUM_ | | |
| ICC | 0.53 | | | 0.63 | | | 0.63 | | |
| N | 234 _M2ID_ | | | 235 _M2ID_ | | | 234 _M2ID_ | | |
|  | 151 _M2FAMNUM_ | | | 151 _M2FAMNUM_ | | | 151 _M2FAMNUM_ | | |
| Observations | 451 | | | 458 | | | 443 | | |
| Marginal R^2^ / Conditional R^2^ | 0.068 / 0.563 | | | 0.284 / 0.732 | | | 0.282 / 0.736 | | |

| **Table S33:** Complex multilevel models with participants nested within families to test Hypotheses 3 and 4. Lag in 2 parts as covariates. | | | | | | | | | |
| --- | --- | --- | --- | --- | --- | --- | --- | --- | --- |
|  | **Episodic Memory** | | | **Executive Functioning** | | | **BTACT** | | |
| *Predictors* | *Estimates* | *CI* | *p* | *Estimates* | *CI* | *p* | *Estimates* | *CI* | *p* |
| Intercept (M2) | 0.01 | -0.12 – 0.14 | 0.901 | -0.00 | -0.09 – 0.08 | 0.947 | 0.01 | -0.08 – 0.09 | 0.902 |
| MIDUS Wave | **-0.13** | **-0.26 – -0.00** | **0.044** | **-0.44** | **-0.51 – -0.37** | **<0.001** | **-0.38** | **-0.45 – -0.31** | **<0.001** |
| Exponent | 0.11 | -0.36 – 0.59 | 0.642 | **0.34** | **0.03 – 0.65** | **0.031** | **0.31** | **0.01 – 0.60** | **0.040** |
| M2 Cog to Neuro Lag | 0.09 | -0.11 – 0.28 | 0.399 | 0.00 | -0.13 – 0.13 | 0.993 | 0.02 | -0.10 – 0.14 | 0.764 |
| Neuro to M3 Cog Lag | -0.04 | -0.19 – 0.11 | 0.611 | **-0.11** | **-0.20 – -0.01** | **0.027** | **-0.09** | **-0.19 – -0.00** | **0.042** |
| Wave X Exponent | -0.01 | -0.47 – 0.46 | 0.976 | -0.16 | -0.42 – 0.10 | 0.228 | -0.11 | -0.36 – 0.14 | 0.375 |
| Wave X M2 Cog to Neuro Lag | -0.09 | -0.29 – 0.10 | 0.332 | **-0.14** | **-0.24 – -0.03** | **0.013** | **-0.14** | **-0.24 – -0.04** | **0.006** |
| Wave X Neuro to M3 Cog Lag | -0.02 | -0.17 – 0.12 | 0.746 | **-0.12** | **-0.20 – -0.04** | **0.003** | **-0.12** | **-0.19 – -0.04** | **0.003** |
| **Random Effects** | | | | | | | | | |
| σ^2^ | 0.47 | | | 0.15 | | | 0.13 | | |
| τ_00_ | 0.56 _M2ID:M2FAMNUM_ | | | 0.30 _M2ID:M2FAMNUM_ | | | 0.27 _M2ID:M2FAMNUM_ | | |
| ICC | 0.54 | | | 0.67 | | | 0.68 | | |
| N | 234 _M2ID_ | | | 235 _M2ID_ | | | 234 _M2ID_ | | |
|  | 151 _M2FAMNUM_ | | | 151 _M2FAMNUM_ | | | 151 _M2FAMNUM_ | | |
| Observations | 451 | | | 458 | | | 443 | | |
| Marginal R^2^ / Conditional R^2^ | 0.022 / 0.554 | | | 0.190 / 0.733 | | | 0.179 / 0.735 | | |

| **Table S34:** Complex multilevel models with participants nested within families to test Hypotheses 5 and 6. Lag in 2 parts as covariates. | | | | | | | | | |
| --- | --- | --- | --- | --- | --- | --- | --- | --- | --- |
|  | **Episodic Memory** | | | **Executive Functioning** | | | **BTACT** | | |
| *Predictors* | *Estimates* | *CI* | *p* | *Estimates* | *CI* | *p* | *Estimates* | *CI* | *p* |
| Intercept (M2) | 0.01 | -0.12 – 0.14 | 0.890 | -0.00 | -0.09 – 0.08 | 0.946 | 0.01 | -0.08 – 0.09 | 0.892 |
| MIDUS Wave | **-0.13** | **-0.26 – -0.00** | **0.045** | **-0.44** | **-0.51 – -0.37** | **<0.001** | **-0.38** | **-0.45 – -0.32** | **<0.001** |
| Alpha Peak | **0.14** | **0.01 – 0.28** | **0.034** | 0.08 | -0.01 – 0.17 | 0.078 | **0.09** | **0.01 – 0.17** | **0.036** |
| M2 Cog to Neuro Lag | 0.10 | -0.09 – 0.29 | 0.314 | 0.02 | -0.10 – 0.15 | 0.704 | 0.04 | -0.08 – 0.16 | 0.487 |
| Neuro to M3 Cog Lag | -0.02 | -0.17 – 0.12 | 0.741 | -0.09 | -0.18 – 0.01 | 0.066 | -0.07 | -0.16 – 0.01 | 0.102 |
| Wave X Alpha Peak | 0.06 | -0.07 – 0.19 | 0.343 | 0.06 | -0.02 – 0.13 | 0.121 | 0.06 | -0.01 – 0.13 | 0.082 |
| Wave X M2 Cog to Neuro Lag | -0.09 | -0.28 – 0.10 | 0.342 | **-0.14** | **-0.25 – -0.04** | **0.008** | **-0.15** | **-0.25 – -0.05** | **0.004** |
| Wave X Neuro to M3 Cog Lag | -0.02 | -0.16 – 0.12 | 0.786 | **-0.13** | **-0.20 – -0.05** | **0.002** | **-0.12** | **-0.19 – -0.04** | **0.002** |
| **Random Effects** | | | | | | | | | |
| σ^2^ | 0.47 | | | 0.14 | | | 0.13 | | |
| τ_00_ | 0.53 _M2ID:M2FAMNUM_ | | | 0.29 _M2ID:M2FAMNUM_ | | | 0.26 _M2ID:M2FAMNUM_ | | |
| ICC | 0.53 | | | 0.67 | | | 0.67 | | |
| N | 234 _M2ID_ | | | 235 _M2ID_ | | | 234 _M2ID_ | | |
|  | 151 _M2FAMNUM_ | | | 151 _M2FAMNUM_ | | | 151 _M2FAMNUM_ | | |
| Observations | 451 | | | 458 | | | 443 | | |
| Marginal R^2^ / Conditional R^2^ | 0.051 / 0.555 | | | 0.201 / 0.735 | | | 0.198 / 0.739 | | |

| **Table S35:** Complex multilevel models with participants nested within families to test Hypotheses 7a and 7c. Lag in 2 parts as covariates. | | | | | | | | | |
| --- | --- | --- | --- | --- | --- | --- | --- | --- | --- |
|  | **Episodic Memory** | | | **Executive Functioning** | | | **BTACT** | | |
| *Predictors* | *Estimates* | *CI* | *p* | *Estimates* | *CI* | *p* | *Estimates* | *CI* | *p* |
| Intercept (M2) | 0.01 | -0.12 – 0.15 | 0.864 | 0.01 | -0.08 – 0.09 | 0.884 | 0.01 | -0.07 – 0.10 | 0.720 |
| MIDUS Wave | -0.11 | -0.24 – 0.02 | 0.106 | **-0.43** | **-0.51 – -0.36** | **<0.001** | **-0.38** | **-0.46 – -0.31** | **<0.001** |
| Exponent | -0.04 | -0.52 – 0.45 | 0.875 | 0.13 | -0.18 – 0.43 | 0.414 | 0.10 | -0.19 – 0.39 | 0.491 |
| Age | **-0.01** | **-0.03 – -0.00** | **0.029** | **-0.02** | **-0.03 – -0.01** | **<0.001** | **-0.02** | **-0.03 – -0.01** | **<0.001** |
| M2 Cog to Neuro Lag | 0.05 | -0.15 – 0.24 | 0.649 | -0.06 | -0.18 – 0.07 | 0.378 | -0.03 | -0.15 – 0.08 | 0.561 |
| Neuro to M3 Cog Lag | -0.06 | -0.20 – 0.09 | 0.437 | **-0.14** | **-0.23 – -0.04** | **0.004** | **-0.12** | **-0.21 – -0.03** | **0.006** |
| Wave X Exponent | -0.14 | -0.62 – 0.33 | 0.557 | -0.19 | -0.46 – 0.07 | 0.157 | -0.15 | -0.41 – 0.11 | 0.253 |
| Wave X Age | -0.01 | -0.02 – 0.00 | 0.067 | -0.00 | -0.01 – 0.00 | 0.372 | -0.00 | -0.01 – 0.00 | 0.199 |
| Exponent X Age | 0.01 | -0.04 – 0.05 | 0.839 | 0.01 | -0.02 – 0.04 | 0.472 | 0.01 | -0.02 – 0.04 | 0.444 |
| Wave X M2 Cog to Neuro Lag | -0.13 | -0.32 – 0.06 | 0.191 | **-0.14** | **-0.25 – -0.04** | **0.010** | **-0.16** | **-0.26 – -0.06** | **0.003** |
| Wave X Neuro to M3 Cog Lag | -0.04 | -0.18 – 0.10 | 0.607 | **-0.13** | **-0.21 – -0.05** | **0.002** | **-0.12** | **-0.20 – -0.05** | **0.002** |
| Wave X Exponent X Age | 0.03 | -0.02 – 0.08 | 0.197 | 0.00 | -0.02 – 0.03 | 0.836 | 0.00 | -0.03 – 0.03 | 0.990 |
| **Random Effects** | | | | | | | | | |
| σ^2^ | 0.46 | | | 0.15 | | | 0.13 | | |
| τ_00_ | 0.53 _M2ID:M2FAMNUM_ | | | 0.25 _M2ID:M2FAMNUM_ | | | 0.22 _M2ID:M2FAMNUM_ | | |
| ICC | 0.53 | | | 0.63 | | | 0.63 | | |
| N | 234 _M2ID_ | | | 235 _M2ID_ | | | 234 _M2ID_ | | |
|  | 151 _M2FAMNUM_ | | | 151 _M2FAMNUM_ | | | 151 _M2FAMNUM_ | | |
| Observations | 451 | | | 458 | | | 443 | | |
| Marginal R^2^ / Conditional R^2^ | 0.072 / 0.567 | | | 0.286 / 0.734 | | | 0.283 / 0.737 | | |

| **Table S36:** Complex multilevel models with participants nested within families to test Hypotheses 7b and 7d. Lag in 2 parts as covariates. | | | | | | | | | |
| --- | --- | --- | --- | --- | --- | --- | --- | --- | --- |
|  | **Episodic Memory** | | | **Executive Functioning** | | | **BTACT** | | |
| *Predictors* | *Estimates* | *CI* | *p* | *Estimates* | *CI* | *p* | *Estimates* | *CI* | *p* |
| Intercept (M2) | 0.01 | -0.13 – 0.14 | 0.918 | 0.00 | -0.08 – 0.09 | 0.916 | 0.01 | -0.07 – 0.09 | 0.783 |
| MIDUS Wave | -0.13 | -0.26 – 0.00 | 0.058 | **-0.45** | **-0.52 – -0.37** | **<0.001** | **-0.39** | **-0.46 – -0.32** | **<0.001** |
| Alpha Peak | 0.11 | -0.02 – 0.25 | 0.107 | 0.03 | -0.06 – 0.11 | 0.512 | 0.04 | -0.04 – 0.12 | 0.332 |
| Age | -0.01 | -0.02 – 0.00 | 0.068 | **-0.02** | **-0.03 – -0.01** | **<0.001** | **-0.02** | **-0.03 – -0.01** | **<0.001** |
| M2 Cog to Neuro Lag | 0.06 | -0.14 – 0.25 | 0.557 | -0.05 | -0.18 – 0.07 | 0.414 | -0.03 | -0.15 – 0.09 | 0.626 |
| Neuro to M3 Cog Lag | -0.05 | -0.19 – 0.10 | 0.513 | **-0.13** | **-0.22 – -0.04** | **0.006** | **-0.11** | **-0.20 – -0.03** | **0.011** |
| Wave X Alpha Peak | 0.04 | -0.10 – 0.17 | 0.602 | 0.05 | -0.03 – 0.12 | 0.200 | 0.05 | -0.02 – 0.12 | 0.188 |
| Wave X Age | -0.01 | -0.02 – 0.00 | 0.090 | -0.00 | -0.01 – 0.01 | 0.724 | -0.00 | -0.01 – 0.00 | 0.469 |
| Alpha Peak X Age | -0.00 | -0.01 – 0.01 | 0.922 | 0.00 | -0.00 – 0.01 | 0.430 | 0.00 | -0.01 – 0.01 | 0.531 |
| Wave X M2 Cog to Neuro Lag | -0.14 | -0.33 – 0.06 | 0.165 | **-0.14** | **-0.25 – -0.03** | **0.011** | **-0.15** | **-0.26 – -0.05** | **0.004** |
| Wave X Neuro to M3 Cog Lag | -0.04 | -0.18 – 0.10 | 0.576 | **-0.13** | **-0.21 – -0.05** | **0.001** | **-0.13** | **-0.20 – -0.05** | **0.001** |
| Wave X Alpha Peak X Age | 0.00 | -0.01 – 0.02 | 0.656 | -0.00 | -0.01 – 0.00 | 0.182 | -0.00 | -0.01 – 0.00 | 0.340 |
| **Random Effects** | | | | | | | | | |
| σ^2^ | 0.47 | | | 0.14 | | | 0.13 | | |
| τ_00_ | 0.51 _M2ID:M2FAMNUM_ | | | 0.25 _M2ID:M2FAMNUM_ | | | 0.22 _M2ID:M2FAMNUM_ | | |
| ICC | 0.52 | | | 0.63 | | | 0.63 | | |
| N | 234 _M2ID_ | | | 235 _M2ID_ | | | 234 _M2ID_ | | |
|  | 151 _M2FAMNUM_ | | | 151 _M2FAMNUM_ | | | 151 _M2FAMNUM_ | | |
| Observations | 451 | | | 458 | | | 443 | | |
| Marginal R^2^ / Conditional R^2^ | 0.082 / 0.563 | | | 0.289 / 0.737 | | | 0.290 / 0.740 | | |

| **Table S37:** Complex multilevel models with participants nested within families, exploratory analyses. Lag in 2 parts as covariates. | | | | | | | | | |
| --- | --- | --- | --- | --- | --- | --- | --- | --- | --- |
|  | **Episodic Memory** | | | **Executive Functioning** | | | **BTACT** | | |
| *Predictors* | *Estimates* | *CI* | *p* | *Estimates* | *CI* | *p* | *Estimates* | *CI* | *p* |
| Intercept (M2) | 0.01 | -0.12 – 0.14 | 0.904 | -0.01 | -0.09 – 0.07 | 0.842 | 0.00 | -0.08 – 0.08 | 0.985 |
| MIDUS Wave | **-0.13** | **-0.26 – -0.00** | **0.047** | **-0.42** | **-0.49 – -0.35** | **<0.001** | **-0.37** | **-0.44 – -0.31** | **<0.001** |
| Exponent | 0.06 | -0.46 – 0.57 | 0.829 | 0.13 | -0.20 – 0.45 | 0.440 | 0.11 | -0.19 – 0.42 | 0.466 |
| Alpha Peak | 0.12 | -0.02 – 0.26 | 0.098 | 0.04 | -0.05 – 0.12 | 0.424 | 0.05 | -0.04 – 0.13 | 0.269 |
| Age | -0.01 | -0.02 – 0.00 | 0.101 | **-0.02** | **-0.03 – -0.01** | **<0.001** | **-0.02** | **-0.03 – -0.01** | **<0.001** |
| M2 Cog to Neuro Lag | 0.05 | -0.14 – 0.25 | 0.590 | -0.06 | -0.19 – 0.06 | 0.338 | -0.04 | -0.16 – 0.08 | 0.528 |
| Neuro to M3 Cog Lag | -0.05 | -0.20 – 0.10 | 0.499 | **-0.14** | **-0.23 – -0.05** | **0.003** | **-0.12** | **-0.21 – -0.04** | **0.006** |
| Wave X Exponent | -0.07 | -0.59 – 0.44 | 0.781 | -0.09 | -0.37 – 0.19 | 0.542 | -0.04 | -0.31 – 0.23 | 0.752 |
| Wave X Alpha Freq | 0.02 | -0.12 – 0.16 | 0.740 | 0.04 | -0.04 – 0.12 | 0.295 | 0.04 | -0.03 – 0.12 | 0.256 |
| Exponent X Alpha Freq | -0.02 | -0.50 – 0.46 | 0.941 | -0.11 | -0.41 – 0.20 | 0.494 | -0.10 | -0.38 – 0.19 | 0.507 |
| Wave X Age | -0.01 | -0.03 – 0.00 | 0.080 | -0.00 | -0.01 – 0.01 | 0.701 | -0.00 | -0.01 – 0.00 | 0.504 |
| Wave X M2 Cog to Neuro Lag | -0.12 | -0.32 – 0.07 | 0.215 | **-0.13** | **-0.24 – -0.02** | **0.024** | **-0.14** | **-0.24 – -0.04** | **0.009** |
| Wave X Neuro to M3 Cog Lag | -0.03 | -0.18 – 0.11 | 0.653 | **-0.11** | **-0.19 – -0.03** | **0.007** | **-0.11** | **-0.18 – -0.03** | **0.006** |
| Wave X Exponent X Alpha Freq | 0.11 | -0.37 – 0.58 | 0.659 | 0.24 | -0.02 – 0.51 | 0.072 | 0.23 | -0.02 – 0.48 | 0.068 |
| **Random Effects** | | | | | | | | | |
| σ^2^ | 0.47 | | | 0.14 | | | 0.12 | | |
| τ_00_ | 0.52 _M2ID:M2FAMNUM_ | | | 0.25 _M2ID:M2FAMNUM_ | | | 0.22 _M2ID:M2FAMNUM_ | | |
| ICC | 0.52 | | | 0.63 | | | 0.64 | | |
| N | 234 _M2ID_ | | | 235 _M2ID_ | | | 234 _M2ID_ | | |
|  | 147 _M2FAMNUM_ | | | 147 _M2FAMNUM_ | | | 147 _M2FAMNUM_ | | |
| Observations | 451 | | | 458 | | | 443 | | |
| Marginal R^2^ / Conditional R^2^ | 0.082 / 0.563 | | | 0.290 / 0.740 | | | 0.292 / 0.744 | | |

| **Table S38:** Examining the Wave X Aperiodic Exponent X Individual Peak Alpha Frequency interaction from analyses described in Table S37 (complex multilevel model, participant nested within family, lag in 2 parts as covariates) through the slope of the change in cognitive function over waves for each BTACT measure at high and low levels of each EEG metric. | | | | | | |
| --- | --- | --- | --- | --- | --- | --- |
| **Change in BTACT Episodic Memory Factor from M2 to M3** | | | | | | |
|  | *Slope of IAPF* | | *95% CI* | | *p-value* | |
| Low Exponent (-1 SD) | -0.01 | | -0.20 – 0.19 | | 0.950 | |
| High Exponent (+1 SD) | 0.07 | | -0.11 – 0.26 | | 0.438 | |
|  | *Slope of Exponent* | | *95% CI* | | *p-value* | |
| Low IAPF (-1 SD) | -0.18 | | -0.76 –0.41 | | 0.552 | |
| High IAPF (+1 SD) | 0.05 | | -0.13 – 0.24 | | 0.578 | |
| **Change in BTACT Executive Functioning Factor from M2 to M3** | | | | | | |
|  | | *Slope of IAPF* | | *95% CI* | | *p-value* |
| Low Exponent (-1 SD) | | -0.03 | | -0.14 – 0.08 | | 0.629 |
| High Exponent (+1 SD) | | **0.11** | | **0.01 – 0.21** | | **0.040** |
|  | | *Slope of Exponent* | | *95% CI* | | *p-value* |
| Low IAPF (-1 SD) | | -0.33 | | -0.66– 0.01 | | 0.055 |
| High IAPF (+1 SD) | | 0.15 | | -0.28 – 0.58 | | 0.487 |
| **Change in BTACT Overall Composite from M2 to M3** | | | | | | |
|  | | *Slope of IAPF* | | *95% CI* | | *p-value* |
| Low Exponent (-1 SD) | | -0.02 | | -0.13 – 0.08 | | 0.670 |
| High Exponent (+1 SD) | | 0.11 | | 0.01 – 0.21 | | **0.033** |
|  | | *Slope of Exponent* | | *95% CI* | | *p-value* |
| Low IAPF (-1 SD) | | -0.27 | | -0.59 – -0.04 | | 0.092 |
| High IAPF (+1 SD) | | 0.18 | | -0.22 – 0.59 | | 0.374 |

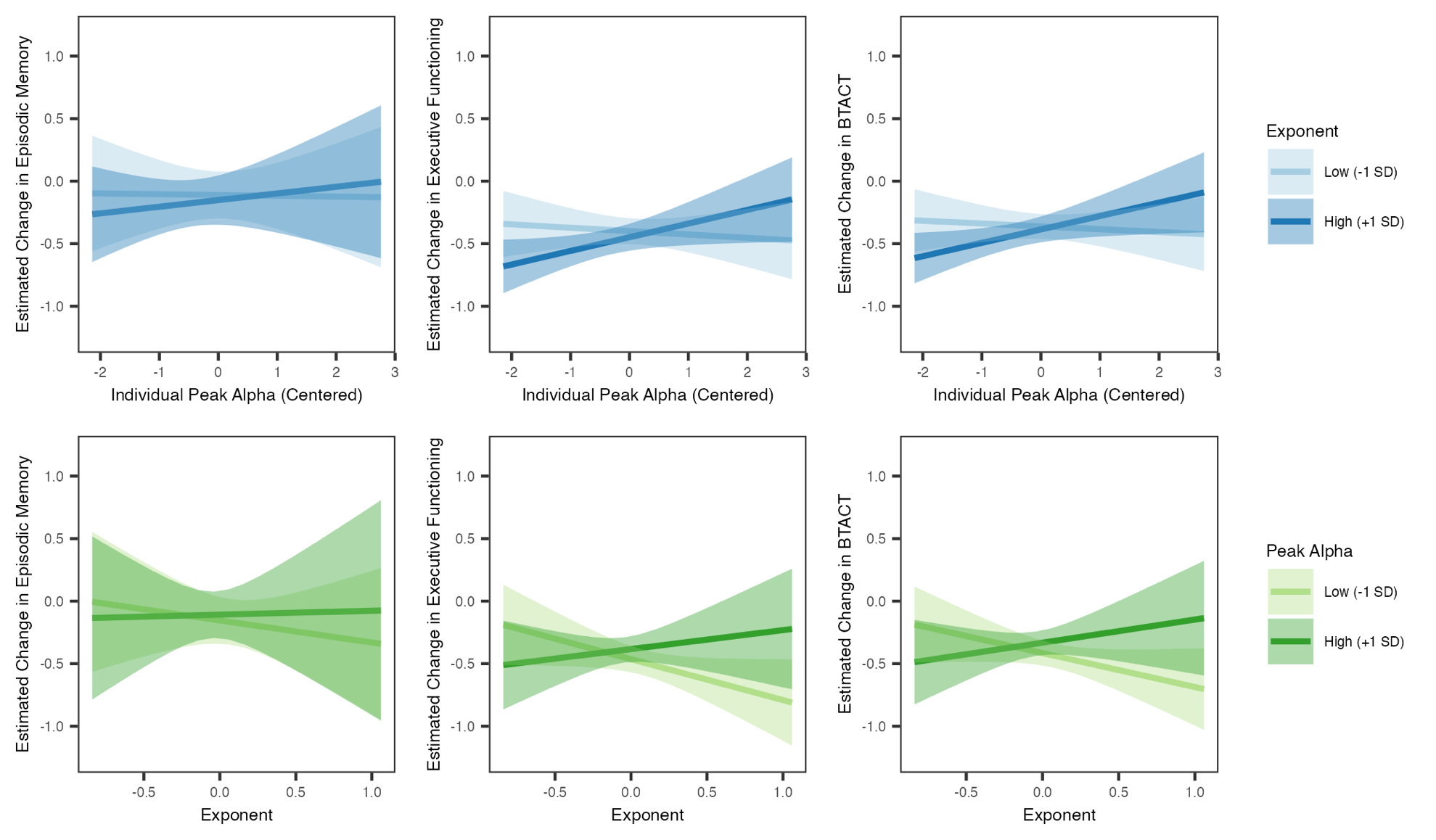

Figure S5. Wave by aperiodic exponent by individual peak alpha frequency interaction plot for models described in Table S30 (complex multilevel model, participant nested within family, lag in 2 parts as covariates).

**Analyses With No Covariates**

| **Table S38: M**ultilevel models with no covariates to test Hypotheses 1 and 2. | | | | | | | | | |
| --- | --- | --- | --- | --- | --- | --- | --- | --- | --- |
|  | **Episodic Memory** | | | **Executive Functioning** | | | **BTACT** | | |
| *Predictors* | *Estimates* | *CI* | *p* | *Estimates* | *CI* | *p* | *Estimates* | *CI* | *p* |
| Intercept (M2) | 0.01 | -0.12 – 0.14 | 0.915 | -0.01 | -0.09 – 0.08 | 0.899 | 0.00 | -0.08 – 0.08 | 0.939 |
| MIDUS Wave | **-0.14** | **-0.26 – -0.01** | **0.037** | **-0.43** | **-0.51 – -0.36** | **<0.001** | **-0.39** | **-0.45 – -0.32** | **<0.001** |
| Age | **-0.02** | **-0.03 – -0.00** | **0.012** | **-0.02** | **-0.03 – -0.01** | **<0.001** | **-0.02** | **-0.03 – -0.01** | **<0.001** |
| MIDUS Wave X Age | -0.01 | -0.02 – 0.00 | 0.099 | -0.00 | -0.01 – 0.01 | 0.979 | -0.00 | -0.01 – 0.01 | 0.712 |
| **Random Effects** | | | | | | | | | |
| σ^2^ | 0.46 | | | 0.15 | | | 0.13 | | |
| τ_00_ | 0.53 _M2ID_ | | | 0.29 _M2ID_ | | | 0.25 _M2ID_ | | |
| ICC | 0.53 | | | 0.65 | | | 0.66 | | |
| N | 234 _M2ID_ | | | 235 _M2ID_ | | | 234 _M2ID_ | | |
| Observations | 451 | | | 458 | | | 443 | | |
| Marginal R^2^ / Conditional R^2^ | 0.052 / 0.558 | | | 0.191 / 0.719 | | | 0.189 / 0.721 | | |

| **Table S39: M**ultilevel models with no covariates to test Hypotheses 3 and 4. | | | | | | | | | |
| --- | --- | --- | --- | --- | --- | --- | --- | --- | --- |
|  | **Episodic Memory** | | | **Executive Functioning** | | | **BTACT** | | |
| *Predictors* | *Estimates* | *CI* | *p* | *Estimates* | *CI* | *p* | *Estimates* | *CI* | *p* |
| Intercept (M2) | 0.01 | -0.12 – 0.14 | 0.911 | -0.00 | -0.09 – 0.09 | 0.923 | 0.00 | -0.08 – 0.09 | 0.927 |
| MIDUS Wave | **-0.13** | **-0.26 – -0.00** | **0.044** | **-0.44** | **-0.51 – -0.36** | **<0.001** | **-0.38** | **-0.45 – -0.32** | **<0.001** |
| Exponent | 0.12 | -0.35 – 0.59 | 0.624 | 0.31 | -0.01 – 0.63 | 0.059 | 0.29 | -0.02 – 0.59 | 0.067 |
| Wave X Exponent | -0.03 | -0.49 – 0.43 | 0.889 | -0.21 | -0.47 – 0.05 | 0.111 | -0.17 | -0.42 – 0.08 | 0.177 |
| **Random Effects** | | | | | | | | | |
| σ^2^ | 0.47 | | | 0.15 | | | 0.13 | | |
| τ_00_ | 0.57 _M2ID_ | | | 0.34 _M2ID_ | | | 0.31 _M2ID_ | | |
| ICC | 0.55 | | | 0.69 | | | 0.70 | | |
| N | 234 _M2ID_ | | | 235 _M2ID_ | | | 234 _M2ID_ | | |
| Observations | 451 | | | 458 | | | 443 | | |
| Marginal R^2^ / Conditional R^2^ | 0.005 / 0.551 | | | 0.096 / 0.722 | | | 0.085 / 0.724 | | |

| **Table S40: M**ultilevel models with no covariates to test Hypotheses 5 and 6. | | | | | | | | | |
| --- | --- | --- | --- | --- | --- | --- | --- | --- | --- |
|  | **Episodic Memory** | | | **Executive Functioning** | | | **BTACT** | | |
| *Predictors* | *Estimates* | *CI* | *p* | *Estimates* | *CI* | *p* | *Estimates* | *CI* | *p* |
| Intercept (M2) | 0.01 | -0.12 – 0.14 | 0.900 | -0.00 | -0.09 – 0.08 | 0.921 | 0.00 | -0.08 – 0.09 | 0.918 |
| MIDUS Wave | **-0.13** | **-0.26 – -0.00** | **0.045** | **-0.44** | **-0.51 – -0.37** | **<0.001** | **-0.39** | **-0.45 – -0.32** | **<0.001** |
| Alpha Peak | **0.15** | **0.02 – 0.28** | **0.028** | 0.09 | -0.00 – 0.18 | 0.061 | **0.10** | **0.01 – 0.19** | **0.027** |
| Wave X Alpha Peak | 0.06 | -0.07 – 0.19 | 0.353 | 0.06 | -0.01 – 0.14 | 0.092 | 0.07 | -0.00 – 0.14 | 0.062 |
| **Random Effects** | | | | | | | | | |
| σ^2^ | 0.47 | | | 0.15 | | | 0.13 | | |
| τ_00_ | 0.54 _M2ID_ | | | 0.33 _M2ID_ | | | 0.29 _M2ID_ | | |
| ICC | 0.54 | | | 0.69 | | | 0.69 | | |
| N | 234 _M2ID_ | | | 235 _M2ID_ | | | 234 _M2ID_ | | |
| Observations | 451 | | | 458 | | | 443 | | |
| Marginal R^2^ / Conditional R^2^ | 0.035 / 0.552 | | | 0.115 / 0.723 | | | 0.114 / 0.728 | | |

| **Table S41: M**ultilevel models with no covariates to test Hypotheses 7a and 7c. | | | | | | | | | |
| --- | --- | --- | --- | --- | --- | --- | --- | --- | --- |
|  | **Episodic Memory** | | | **Executive Functioning** | | | **BTACT** | | |
| *Predictors* | *Estimates* | *CI* | *p* | *Estimates* | *CI* | *p* | *Estimates* | *CI* | *p* |
| Intercept (M2) | 0.01 | -0.13 – 0.15 | 0.873 | 0.01 | -0.08 – 0.10 | 0.903 | 0.01 | -0.07 – 0.10 | 0.745 |
| MIDUS Wave | -0.11 | -0.24 – 0.03 | 0.112 | **-0.43** | **-0.51 – -0.35** | **<0.001** | **-0.38** | **-0.46 – -0.31** | **<0.001** |
| Exponent | -0.06 | -0.54 – 0.42 | 0.803 | 0.07 | -0.25 – 0.39 | 0.654 | 0.05 | -0.25 – 0.36 | 0.729 |
| Age | **-0.02** | **-0.03 – -0.00** | **0.015** | **-0.02** | **-0.03 – -0.01** | **<0.001** | **-0.02** | **-0.03 – -0.01** | **<0.001** |
| Wave X Exponent | -0.16 | -0.63 – 0.32 | 0.515 | -0.23 | -0.50 – 0.04 | 0.096 | -0.20 | -0.46 – 0.06 | 0.141 |
| Wave X Age | -0.01 | -0.02 – 0.00 | 0.126 | -0.00 | -0.01 – 0.01 | 0.668 | -0.00 | -0.01 – 0.00 | 0.490 |
| Exponent X Age | 0.01 | -0.04 – 0.05 | 0.841 | 0.01 | -0.02 – 0.05 | 0.460 | 0.01 | -0.02 – 0.04 | 0.440 |
| Wave X Exponent X Age | 0.03 | -0.02 – 0.08 | 0.175 | 0.01 | -0.02 – 0.03 | 0.708 | 0.00 | -0.02 – 0.03 | 0.737 |
| **Random Effects** | | | | | | | | | |
| σ^2^ | 0.46 | | | 0.15 | | | 0.13 | | |
| τ_00_ | 0.54 _M2ID_ | | | 0.29 _M2ID_ | | | 0.26 _M2ID_ | | |
| ICC | 0.54 | | | 0.65 | | | 0.66 | | |
| N | 234 _M2ID_ | | | 235 _M2ID_ | | | 234 _M2ID_ | | |
| Observations | 451 | | | 458 | | | 443 | | |
| Marginal R^2^ / Conditional R^2^ | 0.058 / 0.563 | | | 0.194 / 0.722 | | | 0.192 / 0.724 | | |

| **Table S42: M**ultilevel models with no covariates to test Hypotheses 7b and 7d. | | | | | | | | | |
| --- | --- | --- | --- | --- | --- | --- | --- | --- | --- |
|  | **Episodic Memory** | | | **Executive Functioning** | | | **BTACT** | | |
| *Predictors* | *Estimates* | *CI* | *p* | *Estimates* | *CI* | *p* | *Estimates* | *CI* | *p* |
| Intercept (M2) | 0.01 | -0.12 – 0.15 | 0.839 | 0.01 | -0.08 – 0.10 | 0.819 | 0.02 | -0.07 – 0.10 | 0.674 |
| MIDUS Wave | -0.13 | -0.26 – 0.00 | 0.051 | **-0.44** | **-0.52 – -0.37** | **<0.001** | **-0.39** | **-0.46 – -0.32** | **<0.001** |
| Alpha | 0.12 | -0.02 – 0.26 | 0.085 | 0.04 | -0.05 – 0.13 | 0.366 | 0.05 | -0.03 – 0.14 | 0.219 |
| MIDUS Wave X Alpha | **-0.01** | **-0.03 – -0.00** | **0.046** | **-0.02** | **-0.03 – -0.01** | **<0.001** | **-0.02** | **-0.03 – -0.01** | **<0.001** |
| MIDUS Wave X Age | 0.04 | -0.10 – 0.17 | 0.574 | 0.06 | -0.01 – 0.14 | 0.110 | 0.06 | -0.01 – 0.14 | 0.093 |
| Alpha X Age | -0.01 | -0.02 – 0.00 | 0.161 | 0.00 | -0.01 – 0.01 | 0.807 | -0.00 | -0.01 – 0.01 | 0.989 |
| Wave x Alpha | 0.00 | -0.01 – 0.02 | 0.715 | 0.01 | -0.00 – 0.01 | 0.146 | 0.01 | -0.00 – 0.01 | 0.169 |
| Wave X Alpha X Age | 0.00 | -0.01 – 0.01 | 0.872 | -0.00 | -0.01 – 0.00 | 0.309 | -0.00 | -0.01 – 0.00 | 0.443 |
| **Random Effects** | | | | | | | | | |
| σ^2^ | 0.47 | | | 0.15 | | | 0.13 | | |
| τ_00_ | 0.52 _M2ID_ | | | 0.28 _M2ID_ | | | 0.25 _M2ID_ | | |
| ICC | 0.53 | | | 0.65 | | | 0.66 | | |
| N | 234 _M2ID_ | | | 235 _M2ID_ | | | 234 _M2ID_ | | |
| Observations | 451 | | | 458 | | | 443 | | |
| Marginal R^2^ / Conditional R^2^ | 0.069 / 0.558 | | | 0.203 / 0.724 | | | 0.206 / 0.728 | | |

| **Table S43: M**ultilevel models with no covariates, exploratory analyses. | | | | | | | | | |
| --- | --- | --- | --- | --- | --- | --- | --- | --- | --- |
|  | **Episodic Memory** | | | **Executive Functioning** | | | **BTACT** | | |
| *Predictors* | *Estimates* | *CI* | *p* | *Estimates* | *CI* | *p* | *Estimates* | *CI* | *p* |
| Intercept (M2) | 0.01 | -0.12 – 0.14 | 0.905 | -0.01 | -0.09 – 0.08 | 0.895 | 0.00 | -0.08 – 0.08 | 0.943 |
| MIDUS Wave | -0.13 | -0.26 – 0.00 | 0.057 | **-0.42** | **-0.49 – -0.35** | **<0.001** | **-0.37** | **-0.44 – -0.30** | **<0.001** |
| Exponent | -0.02 | -0.53 – 0.50 | 0.949 | 0.11 | -0.23 – 0.45 | 0.525 | 0.09 | -0.23 – 0.42 | 0.560 |
| Alpha Peak | 0.11 | -0.03 – 0.24 | 0.139 | 0.04 | -0.05 – 0.13 | 0.364 | 0.05 | -0.03 – 0.14 | 0.234 |
| Age | **-0.02** | **-0.03 – -0.01** | **0.003** | **-0.02** | **-0.03 – -0.01** | **<0.001** | **-0.02** | **-0.03 – -0.01** | **<0.001** |
| Wave X Exponent | 0.06 | -0.43 – 0.54 | 0.822 | -0.10 | -0.37 – 0.17 | 0.456 | -0.05 | -0.31 – 0.20 | 0.679 |
| Wave X Alpha Freq | 0.06 | -0.07 – 0.19 | 0.387 | 0.05 | -0.03 – 0.12 | 0.211 | 0.05 | -0.02 – 0.12 | 0.150 |
| Exponent X Alpha Freq | 0.00 | -0.47 – 0.48 | 0.986 | -0.02 | -0.33 – 0.30 | 0.918 | -0.02 | -0.31 – 0.28 | 0.901 |
| Wave X Exponent X Alpha Freq | 0.15 | -0.31 – 0.62 | 0.524 | **0.30** | **0.04 – 0.57** | **0.024** | **0.29** | **0.04 – 0.54** | **0.022** |
| **Random Effects** | | | | | | | | | |
| σ^2^ | 0.47 | | | 0.15 | | | 0.13 | | |
| τ_00_ | 0.52 _M2ID_ | | | 0.29 _M2ID_ | | | 0.26 _M2ID_ | | |
| ICC | 0.52 | | | 0.66 | | | 0.67 | | |
| N | 234 _M2ID_ | | | 235 _M2ID_ | | | 234 _M2ID_ | | |
| Observations | 451 | | | 458 | | | 443 | | |
| Marginal R^2^ / Conditional R^2^ | 0.066 / 0.555 | | | 0.204 / 0.733 | | | 0.207 / 0.736 | | |

| **Table S44:** Examining the Wave X Aperiodic Exponent X Individual Peak Alpha Frequency interaction from analyses described in Table S43 (no covariates) through the slope of the change in cognitive function over waves for each BTACT measure at high and low levels of each EEG metric. | | | | | | |
| --- | --- | --- | --- | --- | --- | --- |
| **Change in BTACT Episodic Memory Factor from M2 to M3** | | | | | | |
|  | *Slope of IAPF* | | *95% CI* | | *p-value* | |
| Low Exponent (-1 SD) | 0.02 | | -0.18 – 0.21 | | 0.867 | |
| High Exponent (+1 SD) | 0.10 | | -0.08 – 0.28 | | 0.269 | |
|  | *Slope of Exponent* | | *95% CI* | | *p-value* | |
| Low IAPF (-1 SD) | -0.09 | | -0.65 – 0.47 | | 0.747 | |
| High IAPF (+1 SD) | 0.20 | | -0.55 – 0.96 | | 0.598 | |
| **Change in BTACT Executive Functioning Factor from M2 to M3** | | | | | | |
|  | | *Slope of IAPF* | | *95% CI* | | *p-value* |
| Low Exponent (-1 SD) | | -0.05 | | -0.15 – 0.07 | | 0.498 |
| High Exponent (+1 SD) | | **0.13** | | **0.03 – 0.23** | | **0.010** |
|  | | *Slope of Exponent* | | *95% CI* | | *p-value* |
| Low IAPF (-1 SD) | | **-0.40** | | **-0.72– -0.08** | | **0.016** |
| High IAPF (+1 SD) | | 0.20 | | -0.22 – 0.61 | | 0.353 |
| **Change in BTACT Overall Composite from M2 to M3** | | | | | | |
|  | | *Slope of IAPF* | | *95% CI* | | *p-value* |
| Low Exponent (-1 SD) | | -0.03 | | -0.13 – 0.07 | | 0.575 |
| High Exponent (+1 SD) | | **0.13** | | **0.04 – 0.23** | | **0.006** |
|  | | *Slope of Exponent* | | *95% CI* | | *p-value* |
| **Low IAPF (-1 SD)** | | **-0.34** | | **-0.65 – -0.03** | | **0.030** |
| High IAPF (+1 SD) | | 0.23 | | -0.17 – 0.63 | | 0.254 |

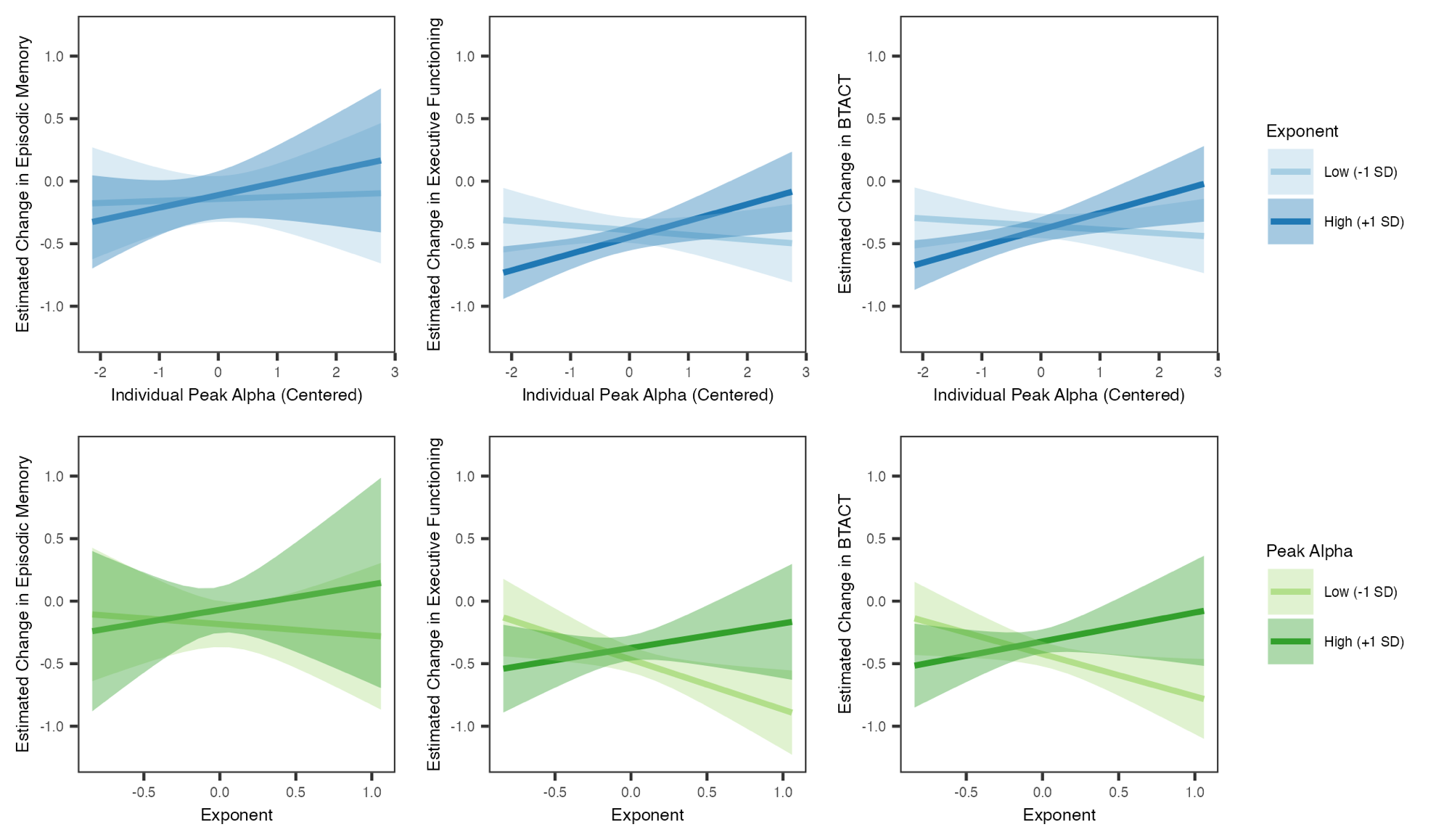

Figure S6. Wave by aperiodic exponent by individual peak alpha frequency interaction plot for models described in Table S43 (no covariates).

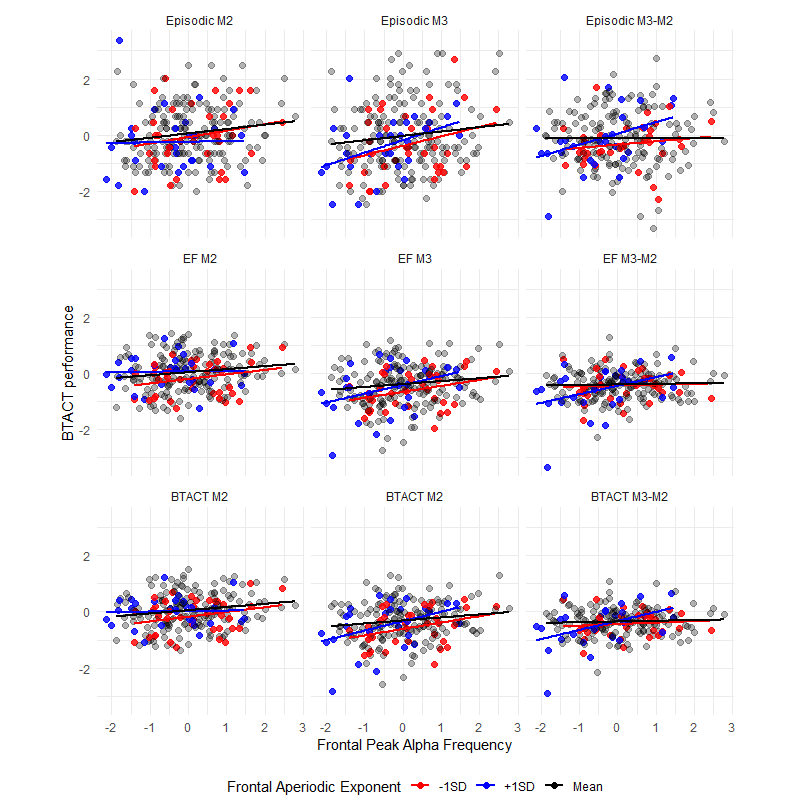
Figure S7. Composite and domain specific BTACT performance as a function of function of frontal peak alpha frequency, and low (-1SD) high (+1SD), and mean frontal aperiodic exponent. The association between frontal peak alpha frequency and change in cognitive performance is strongest in participants with +1SD aperiodic exponents, with decline occurring in those participants who also have slower than average frontal peak alpha frequency.

**Offset Models Parallel to Manuscript**

Next, we examined the relationships between aperiodic offset and cognitive decline in a parallel manner to the aperiodic exponent slope analyses reported in the main manuscript. There was a significant Wave by Offset by Individual Peak Alpha Frequency interaction for the BTACT composite (*p* = 0.026; see Table S47). As shown in Table S48 and Figure S7, it is similar in shape such that individuals with “matching” offsets and individual peak alpha frequencies (i.e., high on both or low on both) show less decline than individuals with “mismatched” metrics, particularly for individuals with high offsets. Given the exploratory nature of this analysis and the weakness of the effect compared to the analyses with the aperiodic exponent, we are hesitant to interpret this effect and believe additional research is needed.

| **Table S45:** Simple multilevel models with offset parallel to models testing hypotheses 3 and 4. Sex education, and lag in one part as covariates. | | | | | | | | | |
| --- | --- | --- | --- | --- | --- | --- | --- | --- | --- |
|  | **Episodic Memory** | | | **Executive Functioning** | | | **BTACT** | | |
| *Predictors* | *Estimates* | *CI* | *p* | *Estimates* | *CI* | *p* | *Estimates* | *CI* | *p* |
| Intercept (M2) | **-0.40** | **-0.58 – -0.22** | **<0.001** | -0.04 | -0.17 – 0.09 | 0.522 | -0.10 | -0.22 – 0.02 | 0.114 |
| MIDUS Wave | -0.13 | -0.26 – 0.00 | 0.051 | **-0.43** | **-0.50 – -0.36** | **<0.001** | **-0.38** | **-0.45 – -0.31** | **<0.001** |
| Offset | 0.00 | -0.26 – 0.26 | 0.986 | 0.11 | -0.08 – 0.29 | 0.250 | 0.10 | -0.08 – 0.27 | 0.272 |
| Sex | **0.69** | **0.47 – 0.91** | **<0.001** | 0.07 | -0.09 – 0.23 | 0.391 | **0.18** | **0.03 – 0.33** | **0.021** |
| Education | **0.18** | **0.05 – 0.30** | **0.008** | **0.13** | **0.05 – 0.22** | **0.002** | **0.13** | **0.05 – 0.21** | **0.002** |
| Lag between Waves | -0.07 | -0.19 – 0.05 | 0.259 | **-0.19** | **-0.27 – -0.10** | **<0.001** | **-0.17** | **-0.25 – -0.09** | **<0.001** |
| Wave x Offset | 0.01 | -0.26 – 0.27 | 0.969 | -0.08 | -0.23 – 0.07 | 0.308 | -0.09 | -0.24 – 0.07 | 0.269 |
| **Random Effects** | | | | | | | | | |
| σ^2^ | 0.47 | | | 0.16 | | | 0.14 | | |
| τ_00_ | 0.44 _M2ID_ | | | 0.29 _M2ID_ | | | 0.25 _M2ID_ | | |
| ICC | 0.48 | | | 0.65 | | | 0.65 | | |
| N | 234 _M2ID_ | | | 235 _M2ID_ | | | 234 _M2ID_ | | |
| Observations | 451 | | | 458 | | | 443 | | |
| Marginal R^2^ / Conditional R^2^ | 0.131 / 0.550 | | | 0.168 / 0.707 | | | 0.168 / 0.708 | | |

| **Table S46:** Simple multilevel models with offset parallel to models testing hypotheses 7a and 7c. Sex education, and lag in 1 part as covariates. | | | | | | | | | |
| --- | --- | --- | --- | --- | --- | --- | --- | --- | --- |
|  | **Episodic Memory** | | | **Executive Functioning** | | | **BTACT** | | |
| *Predictors* | *Estimates* | *CI* | *p* | *Estimates* | *CI* | *p* | *Estimates* | *CI* | *p* |
| Intercept (M2) | **-0.42** | **-0.60 – -0.24** | **<0.001** | -0.07 | -0.19 – 0.05 | 0.241 | **-0.13** | **-0.24 – -0.01** | **0.027** |
| MIDUS Wave | **-0.13** | **-0.26 – -0.00** | **0.045** | **-0.43** | **-0.50 – -0.35** | **<0.001** | **-0.39** | **-0.46 – -0.31** | **<0.001** |
| Offset | -0.06 | -0.31 – 0.20 | 0.668 | 0.05 | -0.12 – 0.22 | 0.554 | 0.04 | -0.12 – 0.20 | 0.614 |
| Age | **-0.02** | **-0.03 – -0.01** | **0.002** | **-0.02** | **-0.03 – -0.02** | **<0.001** | **-0.02** | **-0.03 – -0.02** | **<0.001** |
| Sex | **0.72** | **0.51 – 0.94** | **<0.001** | 0.12 | -0.03 – 0.26 | 0.123 | **0.23** | **0.09 – 0.36** | **0.001** |
| Education | **0.17** | **0.04 – 0.29** | **0.008** | **0.13** | **0.05 – 0.22** | **0.001** | **0.14** | **0.06 – 0.21** | **<0.001** |
| Lag between Waves | -0.10 | -0.21 – 0.02 | 0.093 | **-0.22** | **-0.29 – -0.14** | **<0.001** | **-0.20** | **-0.27 – -0.13** | **<0.001** |
| Wave x Exponent | -0.01 | -0.28 – 0.26 | 0.954 | -0.09 | -0.25 – 0.07 | 0.260 | -0.08 | -0.24 – 0.07 | 0.275 |
| Wave X Age | -0.01 | -0.02 – 0.00 | 0.097 | -0.00 | -0.01 – 0.01 | 0.915 | -0.00 | -0.01 – 0.00 | 0.604 |
| Offset X Age | 0.01 | -0.02 – 0.03 | 0.608 | -0.00 | -0.02 – 0.01 | 0.758 | 0.00 | -0.01 – 0.02 | 0.921 |
| Wave X Offset X Age | -0.00 | -0.03 – 0.03 | 0.988 | 0.00 | -0.01 – 0.02 | 0.786 | -0.01 | -0.02 – 0.01 | 0.278 |
| **Random Effects** | | | | | | | | | |
| σ^2^ | 0.47 | | | 0.16 | | | 0.14 | | |
| τ_00_ | 0.39 _M2ID_ | | | 0.22 _M2ID_ | | | 0.19 _M2ID_ | | |
| ICC | 0.45 | | | 0.59 | | | 0.57 | | |
| N | 234 _M2ID_ | | | 235 _M2ID_ | | | 234 _M2ID_ | | |
| Observations | 451 | | | 458 | | | 443 | | |
| Marginal R^2^ / Conditional R^2^ | 0.195 / 0.558 | | | 0.292 / 0.708 | | | 0.312 / 0.707 | | |

| **Table S47:** Simple multilevel models with offset parallel to exploratory models. Sex education, and lag in 1 part as covariates. | | | | | | | | | |
| --- | --- | --- | --- | --- | --- | --- | --- | --- | --- |
|  | **Episodic Memory** | | | **Executive Functioning** | | | **BTACT** | | |
| *Predictors* | *Estimates* | *CI* | *p* | *Estimates* | *CI* | *p* | *Estimates* | *CI* | *p* |
| Intercept (M2) | **-0.44** | **-0.62 – -0.27** | **<0.001** | -0.06 | -0.18 – 0.06 | 0.312 | **-0.12** | **-0.23 – -0.01** | **0.034** |
| MIDUS Wave | -0.10 | -0.23 – 0.03 | 0.136 | **-0.41** | **-0.49 – -0.34** | **<0.001** | **-0.36** | **-0.43 – -0.29** | **<0.001** |
| Offset | -0.09 | -0.36 – 0.19 | 0.539 | 0.06 | -0.12 – 0.24 | 0.518 | 0.04 | -0.13 – 0.21 | 0.628 |
| Alpha Peak | 0.07 | -0.06 – 0.20 | 0.302 | 0.02 | -0.07 – 0.10 | 0.694 | 0.03 | -0.06 – 0.11 | 0.543 |
| Age | **-0.02** | **-0.03 – -0.01** | **<0.001** | **-0.02** | **-0.03 – -0.02** | **<0.001** | **-0.02** | **-0.03 – -0.02** | **<0.001** |
| Sex | **0.73** | **0.52 – 0.93** | **<0.001** | **0.10** | **-0.04 – 0.25** | 0.158 | **0.21** | **0.08 – 0.35** | **0.002** |
| Education | **0.17** | **0.05 – 0.30** | **0.006** | **0.14** | **0.06 – 0.22** | **0.001** | **0.14** | **0.07 – 0.22** | **<0.001** |
| Lag between Waves | -0.10 | -0.21 – 0.02 | 0.097 | **-0.21** | **-0.29 – -0.13** | **<0.001** | **-0.19** | **-0.27 – -0.12** | **<0.001** |
| Wave X Offset | 0.15 | -0.14 – 0.44 | 0.316 | -0.01 | -0.18 – 0.15 | 0.888 | 0.00 | -0.16 – 0.16 | 0.960 |
| Wave X Alpha Freq | 0.07 | -0.06 – 0.21 | 0.301 | 0.06 | -0.02 – 0.14 | 0.116 | 0.06 | -0.01 – 0.13 | 0.108 |
| Offset X Alpha Freq | -0.19 | -0.45 – 0.07 | 0.154 | 0.01 | -0.16 – 0.18 | 0.907 | -0.02 | -0.18 – 0.14 | 0.769 |
| Wave X Offset X Alpha Freq | 0.25 | -0.02 – 0.52 | 0.066 | 0.14 | -0.02 – 0.30 | 0.081 | **0.17** | **0.02 – 0.32** | **0.026** |
| **Random Effects** | | | | | | | | | |
| σ^2^ | 0.46 | | | 0.15 | | | 0.13 | | |
| τ_00_ | 0.38 _M2ID_ | | | 0.22 _M2ID_ | | | 0.19 _M2ID_ | | |
| ICC | 0.45 | | | 0.59 | | | 0.59 | | |
| N | 234 _M2ID_ | | | 235 _M2ID_ | | | 234 _M2ID_ | | |
| Observations | 451 | | | 458 | | | 443 | | |
| Marginal R^2^ / Conditional R^2^ | 0.205 / 0.564 | | | 0.301 / 0.717 | | | 0.321 / 0.722 | | |

| **Table S48:** Examining the Wave X Aperiodic Offset X Individual Peak Alpha Frequency interaction from analyses described in Table S47 (simple model with aperiodic offset instead of exponent. Sex, education, and lag in 1 part as covariates) through the slope of the change in cognitive function over waves for each BTACT measure at high and low levels of each EEG metric. | | | | | | |
| --- | --- | --- | --- | --- | --- | --- |
| **Change in BTACT Episodic Memory Factor from M2 to M3** | | | | | | |
|  | *Slope of IAPF* | | *95% CI* | | *p-value* | |
| Low Offset (-1 SD) | -0.05 | | -0.24 – 0.14 | | 0.602 | |
| High Offset (+1 SD) | **0.19** | | **0.01 – 0.37** | | **0.037** | |
|  | *Slope of Offset* | | *95% CI* | | *p-value* | |
| Low IAPF (-1 SD) | -0.10 | | -0.42 – 0.23 | | 0.555 | |
| High IAPF (+1 SD) | 0.40 | | -0.06 – 0.85 | | 0.085 | |
| **Change in BTACT Executive Functioning Factor from M2 to M3** | | | | | | |
|  | | *Slope of IAPF* | | *95% CI* | | *p-value* |
| Low Offset (-1 SD) | | -0.01 | | -0.12 – 0.11 | | 0.907 |
| High Offset (+1 SD) | | **0.13** | | **0.02 – 0.24** | | **0.016** |
|  | | *Slope of Offset* | | *95% CI* | | *p-value* |
| Low IAPF (-1 SD) | | -0.15 | | -0.34 – 0.04 | | 0.123 |
| High IAPF (+1 SD) | | 0.13 | | -0.13 – 0.38 | | 0.333 |
| **Change in BTACT Overall Composite from M2 to M3** | | | | | | |
|  | | *Slope of IAPF* | | *95% CI* | | *p-value* |
| Low Offset (-1 SD) | | -0.02 | | -0.13 – 0.08 | | 0.676 |
| High Offset (+1 SD) | | **0.14** | | **0.04 – 0.24** | | **0.005** |
|  | | *Slope of Offset* | | *95% CI* | | *p-value* |
| Low IAPF (-1 SD) | | -0.16 | | -0.35 – 0.02 | | 0.084 |
| High IAPF (+1 SD) | | 0.17 | | -0.08 – 0.42 | | 0.173 |

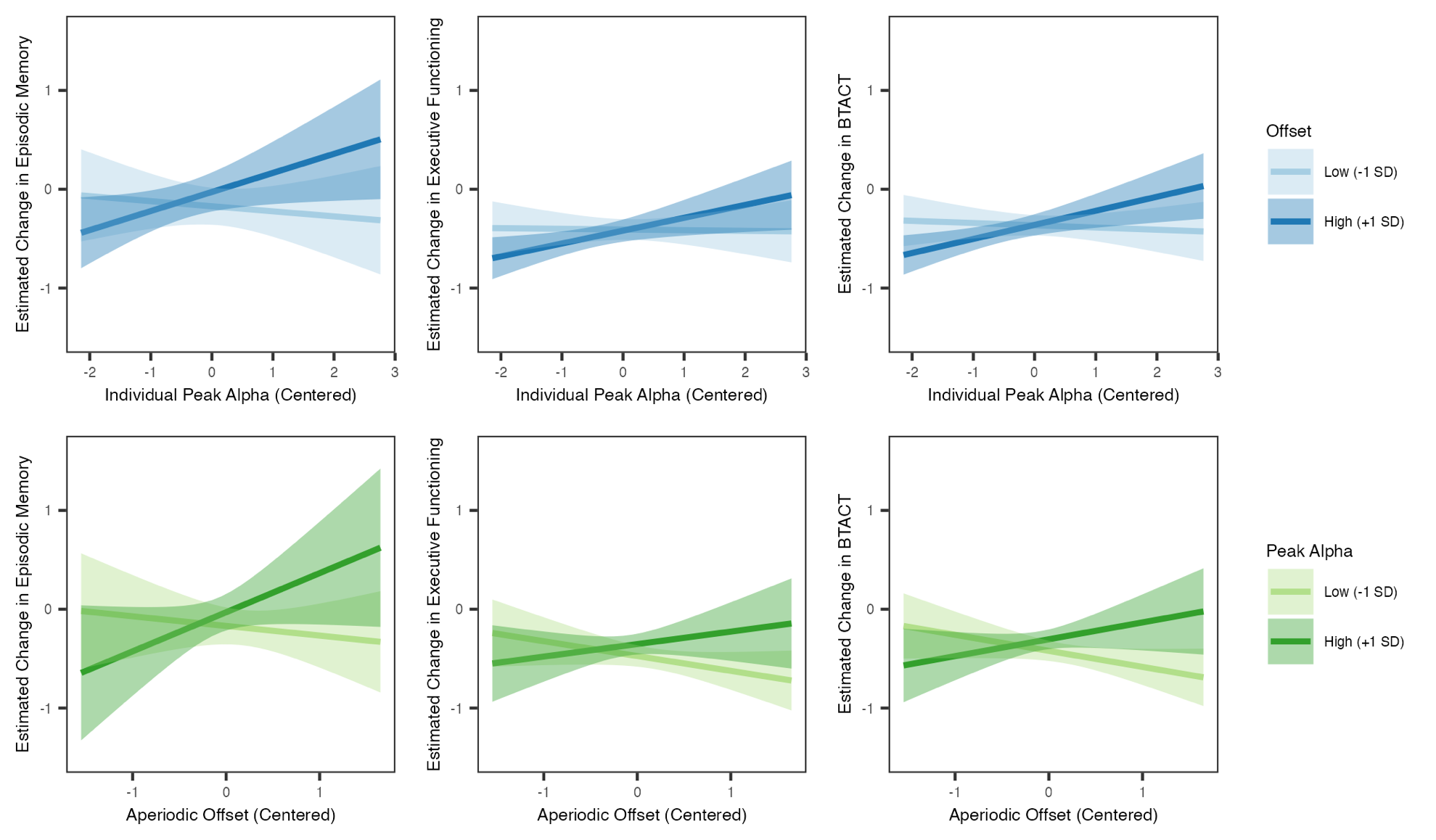

Figure S8. Wave by aperiodic offset by individual peak alpha frequency interaction plot for models described in Table S47 (simple model with aperiodic offset instead of exponent. Sex, education, and lag in 1 part as covariates).

**Parietal Composite Model**

Next, we examined the exploratory aperiodic exponent by individual alpha peak frequency by wave interaction using a parietal composite of P3/Pz/P4 (using sensors 38, 43, 52, 53, 54, 60, 61, 62, 79, 80, 86, 87, 88, 93, and 94), and include a correlation table of the parietal composites with the cognitive measures (Table S49). Overall results are nearly identical to the frontal composite (BTACT composite *p* = 0.009), but the overall interaction remained non -significant for episodic memory (*p* = 0.090).

| **Table S49:** Correlations and descriptive statistics with parietal composite. | | | | | | | | | | |
| --- | --- | --- | --- | --- | --- | --- | --- | --- | --- | --- |
|  | Mean (SD)  n | 2.  Total Lag | 3. M2 Episodic Memory | 4. M3 Episodic Memory | 5. M2 Executive Function | 6. M3 Executive Function | 7. M2 BTACT Composite | 8. M3 BTACT Composite | 9. Aperiodic Exponent (parietal) | 10.  IAPF (parietal) |
| 1. Age (MIDUS 2 Neuro) | 55.10 (10.71) *n* = 235 | -0.12  [-0.25, 0.01]  *p* = 0.064  *n* = 335 | **-0.18**  **[-0.30,-0.05]**  ***p* = 0.006**  *n* = 228 | **-0.24**  **[-0.36,-0.11]**  ***p* < 0.001**  *n* = 223 | **-0.35**  **[-0.46,-0.23]**  ***p* < 0.001**  *n* = 228 | **-0.32**  **[-0.43,-0.20]**  ***p* < 0.001**  *n* = 230 | **-0.36**  **[-0.47,-0.24]**  ***p* < 0.001**  *n* = 224 | **-0.34**  **[-0.45,-0.21]**  ***p* < 0.001**  *n* = 219 | **-0.25**  **[-0.36,-0.12]**  ***p* < 0.001**  *n* = 235 | **-0.16**  **[-0.28,-0.04]**  ***p* = 0.013**  *n* = 235 |
| 2. Total Lag (M2 to M3 Cognitive Projects, yrs) | 9.71 (0.92) *n* = 235 | - | -0.06  [-0.19, 0.07]  *p* = 0.345  *n* = 228 | -0.07  [-0.20, 0.06]  *p* = 0.314  *n* = 223 | **-0.17**  **[-0.29,-0.04]**  ***p* = 0.011**  *n* = 228 | **-0.30**  **[-0.41,-0.17]**  ***p* < 0.001**  *n* = 230 | **-0.16**  **[-0.29,-0.03]**  ***p* = 0.015**  *n* = 224 | **-0.29**  **[-0.41,-0.17**]  ***p* < 0.001**  *n* = 219 | 0.11  [-0.02, 0.23]  *p* = 0.102  *n* = 235 | -0.05  [-0.18, 0.08]  *p* = 0.418  *n* = 224 |
| 3. Episodic Memory MIDUS 2 | 0.00 (0.93) *n* = 228 |  | - | **0.56**  **[0.46, 0.64]**  ***p* < 0.001**  *n* = 217 | **0.36**  **[0.24, 0.47]**  ***p* < 0.001**  *n* = 224 | **0.32**  **[0.19, 0.43]**  ***p* < 0.001**  *n* = 223 | **0.56**  **[0.47, 0.65]**  ***p* < 0.001**  *n* = 224 | **0.41**  **[0.29, 0.52]**  ***p* < 0.001**  *n* = 213 | 0.01  [-0.12, 0.14]  *p* = 0.865  *n* = 228 | **0.17**  **[0.04, 0.29]**  ***p* = 0.011**  *n* = 228 |
| 4. Episodic Memory MIDUS 3 | -0.11 (1.10) *n* = 223 |  |  | - | **0.26**  **[0.13, 0.38]**  ***p* < 0.001**  *n* = 217 | **0.37**  **[0.25, 0.48]**  ***p* < 0.001**  *n* = 219 | **0.39**  **[0.27, 0.50]**  ***p* < 0.001**  *n* = 213 | **0.59**  **[0.49, 0.67]**  ***p* < 0.001**  *n* = 219 | < 0.001  [-0.13, 0.13]  *p* = 0.996  *n* = 223 | **0.23**  **[0.11, 0.35]**  ***p* < 0.001**  *n* = 223 |
| 5. Executive Function MIDUS 2 | 0.01 (0.65) *n* = 228 |  |  |  | - | **0.69**  **[0.62, 0.76]**  ***p* < 0.001**  *n* = 223 | **0.97**  **[0.96, 0.98]**  ***p* < 0.001**  *n* = 222 | **0.68**  **[0.60, 0.74]**  ***p* < 0.001**  *n* = 211 | 0.10  [-0.03, 0.23]  *p* = 0.123  *n* = 228 | **0.16**  **[0.03, 0.28]**  ***p* = 0.016**  ***n* = 228** |
| 6. Executive Function MIDUS 3 | -0.44 (0.74) *n* = 230 |  |  |  |  | - | **0.69**  **[0.61, 0.75]**  ***p* < 0.001**  *n* = 219 | **0.97**  **[0.96, 0.98]**  ***p* < 0.001**  *n* = 219 | 0.01  [-0.12, 0.14]  *p* = 0.849  *n* = 230 | **0.20**  **[0.07, 0.32]**  ***p* = 0.003**  *n* = 230 |
| 7. BTACT Composite MIDUS 2 | 0.01 (0.62) *n* = 224 |  |  |  |  |  | - | **0.70**  **[0.63, 0.77]**  ***p* < 0.001**  *n* = 209 | 0.10  [-0.03, 0.23]  *p* = 0.141  *n* = 224 | **0.19**  **[0.06, 0.31]**  ***p* = 0.004**  *n* = 224 |
| 8. BTACT Composite MIDUS 3 | -0.37 (0.71) *n* = 219 |  |  |  |  |  |  | - | 0.01  [-0.12, 0.14]  *p* = 0.854  *n* = 219 | **0.22**  **[0.09, 0.35]**  ***p* = 0.001**  *n* = 219 |
| 9. Aperiodic Exponent (parietal) | 1.23 (0.27) *n* = 235 |  |  |  |  |  |  |  | - | **-0.28**  **[-0.39-0.15]**  ***p* < 0.001**  *n* = 235 |
| 10. IAPF (parietal) | 9.81 (0.99) *n* = 235 |  |  |  |  |  |  |  |  | - |

| **Table S50:** Multilevel models examine the interaction between aperiodic exponent and individual peak alpha frequency over parietal pooling | | | | | | | | | |
| --- | --- | --- | --- | --- | --- | --- | --- | --- | --- |
|  | **Episodic Memory** | | | **Executive Functioning** | | | **BTACT** | | |
| *Predictors* | *Estimates* | *CI* | *p* | *Estimates* | *CI* | *p* | *Estimates* | *CI* | *p* |
| Intercept (M2) | **-0.44** | **-0.61 – -0.26** | **<0.001** | -0.06 | -0.18 – 0.06 | 0.354 | **-0.12** | **-0.23 – -0.00** | **0.044** |
| MIDUS Wave | -0.10 | -0.23 – 0.03 | 0.126 | **-0.40** | **-0.48 – -0.33** | **<0.001** | **-0.36** | **-0.43 – -0.29** | **<0.001** |
| Exponent | 0.01 | -0.48 – 0.50 | 0.969 | 0.15 | -0.17 – 0.48 | 0.357 | 0.15 | -0.16 – 0.45 | 0.341 |
| Individual Alpha Peak Frequency | 0.09 | -0.04 – 0.22 | 0.166 | 0.04 | -0.04 – 0.13 | 0.314 | 0.06 | -0.02 – 0.14 | 0.164 |
| Age | **-0.02** | **-0.03 – -0.01** | **<0.001** | **-0.02** | **-0.03 – -0.02** | **<0.001** | **-0.02** | **-0.03 – -0.02** | **<0.001** |
| Sex | **0.72** | **0.51 – 0.93** | **<0.001** | 0.10 | -0.05 – 0.24 | 0.181 | **0.21** | **0.07 – 0.34** | **0.003** |
| Education | **0.17** | **0.05 – 0.30** | **0.007** | **0.15** | **0.07 – 0.23** | **<0.001** | **0.14** | **0.07 – 0.22** | **<0.001** |
| Lag between Waves | -0.09 | -0.20 – 0.02 | 0.117 | **-0.20** | **-0.28 – -0.12** | **<0.001** | **-0.18** | **-0.26 – -0.11** | **<0.001** |
| Wave X Exponent | 0.21 | -0.29 – 0.71 | 0.416 | -0.07 | -0.36 – 0.22 | 0.623 | -0.04 | -0.31 – 0.24 | 0.779 |
| Wave X IPAF | 0.12 | -0.01 – 0.25 | 0.079 | 0.05 | -0.02 – 0.13 | 0.157 | 0.05 | -0.02 – 0.13 | 0.149 |
| Exponent X IPAF | -0.22 | -0.65 – 0.21 | 0.310 | 0.03 | -0.25 – 0.32 | 0.819 | -0.01 | -0.27 – 0.25 | 0.950 |
| Wave X Exponent X IPAF | 0.39 | -0.06 – 0.83 | 0.090 | **0.35** | **0.10 – 0.61** | **0.007** | **0.32** | **0.08 – 0.57** | **0.009** |
| **Random Effects** |  | | |  | | |  | | |
| σ^2^ | 0.46 | | | 0.15 | | | 0.13 | | |
| τ_00_ | 0.37 _M2ID_ | | | 0.22 _M2ID_ | | | 0.19 _M2ID_ | | |
| ICC | 0.45 | | | 0.60 | | | 0.59 | | |
| N | 234 _M2ID_ | | | 235 _M2ID_ | | | 234 _M2ID_ | | |
| Observations | 451 | | | 458 | | | 443 | | |
| Marginal R^2^ / Conditional R^2^ | 0.216 / 0.565 | | | 0.313 / 0.722 | | | 0.332 / 0.724 | | |
| Note: Repeated measured cognitive data nested within participant, indicated by M2ID. Sex coded as 0 = male, 1 = female. Education coded as -1 = high school or less, 0 = some college, 1 = Bachelor’s degree or higher. MIDUS Wave coded as 0 = MIDUS 2, 1 = MIDUS 3. Years between Waves (i.e., between MIDUS 2 and MIDUS 3 Cognitive Projects) and Age at M2 Neuroscience Project are mean centered. Empirical Bayes slope estimation used (Bates et al., 2015). | | | | | | | | | |
| Table S51: Examining the Wave X Aperiodic Exponent X Individual Peak Alpha Frequency interaction from analyses described in Table S50 (parietal pooling) through the slope of the change in cognitive function over waves for each BTACT measure at high and low levels of each EEG metric. | | | | | | | | | |
| **Change in BTACT Episodic Memory Factor from M2 to M3** | | | | | | | | | |

|  | *Slope of IAPF* | | *95% CI* | | *p-value* | |
| --- | --- | --- | --- | --- | --- | --- |
| Low Exponent (-1 SD) | 0.02 | | -0.16 – 0.19 | | 0.860 | |
| High Exponent (+1 SD) | **0.22** | | **0.04 – 0.40** | | **0.015** | |
|  | *Slope of Exponent* | | *95% CI* | | *p-value* | |
| Low IAPF (-1 SD) | -0.17 | | -0.76 – 0.41 | | 0.557 | |
| High IAPF (+1 SD) | 0.59 | | -0.16 – 1.34 | | 0.120 | |
| **Change in BTACT Executive Functioning Factor from M2 to M3** | | | | | | |
|  | | *Slope of IAPF* | | *95% CI* | | *p-value* |
| Low Exponent (-1 SD) | | -0.04 | | -0.14 – 0.06 | | 0.450 |
| High Exponent (+1 SD) | | **0.15** | | **0.05 – 0.25** | | **0.004** |
|  | | *Slope of Exponent* | | *95% CI* | | *p-value* |
| Low IAPF (-1 SD) | | **-0.42** | | **-0.76– -0.08** | | **0.015** |
| High IAPF (+1 SD) | | 0.28 | | -0.15 – 0.70 | | 0.202 |
| **Change in BTACT Overall Composite from M2 to M3** | | | | | | |
|  | | *Slope of IAPF* | | *95% CI* | | *p-value* |
| Low Exponent (-1 SD) | | -0.03 | | -0.13 – 0.06 | | 0.505 |
| High Exponent (+1 SD) | | **0.14** | | **0.04 – 0.24** | | **0.005** |
|  | | *Slope of Exponent* | | *95% CI* | | *p-value* |
| Low IAPF (-1 SD) | | **-0.36** | | **-0.68 – -0.04** | | **0.029** |
| High IAPF (+1 SD) | | 0.28 | | -0.12 – 0.69 | | 0.171 |

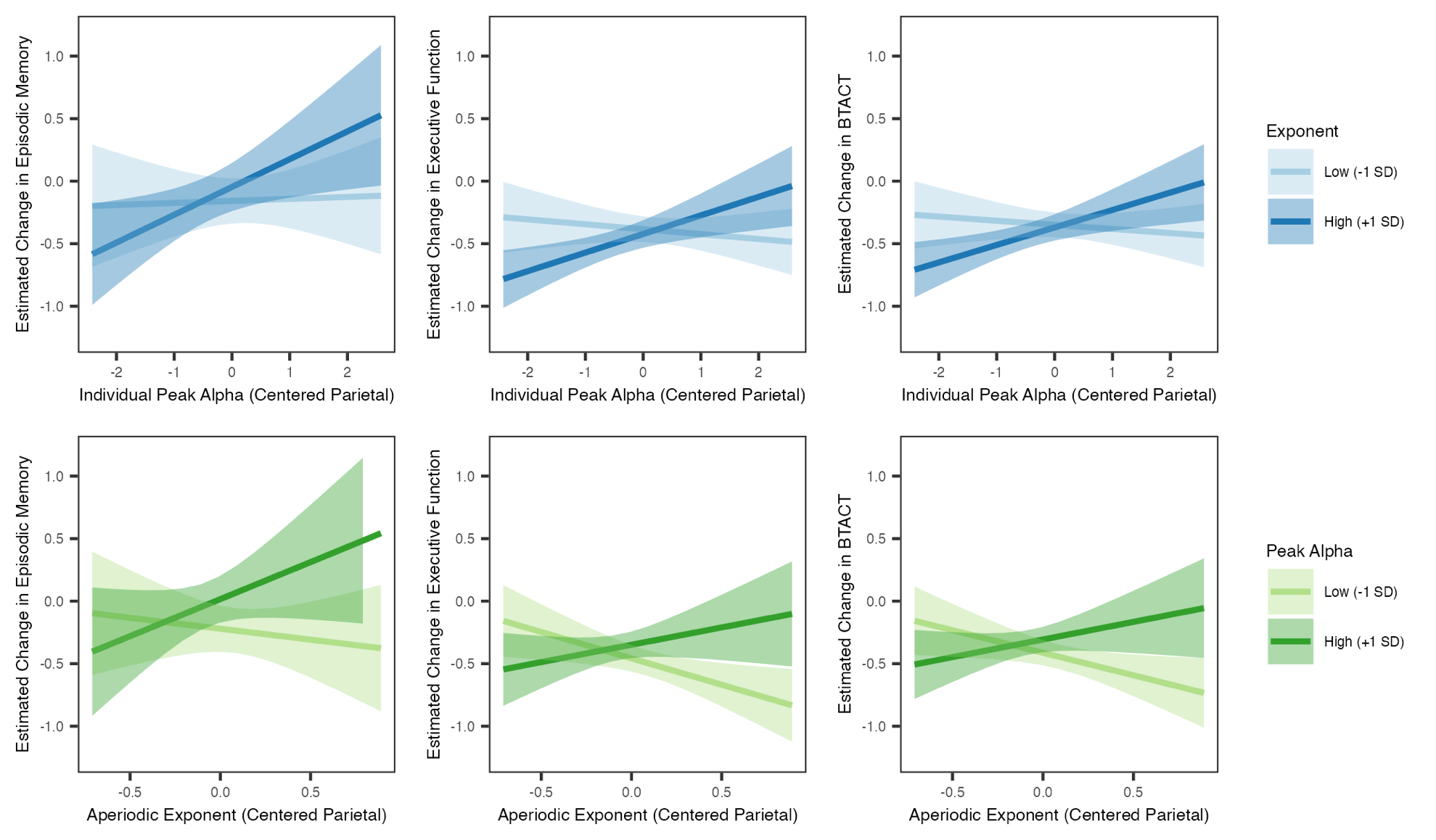

Figure S9. Wave by aperiodic exponent by individual peak alpha frequency interaction plot for models using parietal pooling described in Table S50.

**Eyes Open and Eyes Closed Only Models**

Finally, we examined the correlations between the cognitive measures and EEG metrics calculated using only eyes open and eyes closed data (Table S52-S53), as well as exploratory aperiodic exponent by individual alpha peak frequency by wave interaction using eyes open and eyes closed only data (Tables S54-S57). Overall, as shown in Table S53 the eyes open and eyes closed metrics are highly correlated (r’s = 0.58 to 0.97), and the results are largely the same although attenuated in eyes open only, and the Episodic Memory factor interaction disappears for eyes open only data. Figure S12 depicts the grand-average spectra with FOOOF model fits for the eyes open only, eyes closed only, and combined eyes open and eyes closed data.

Table S52: Correlations, with eyes closed only EEG metrics above the diagonal and eyes open only EEG metrics below the diagonal.

|  | 1. Age (MIDUS 2 Neuro) | 2.  Total Lag | 3. M2 Episodic Memory | 4. M3 Episodic Memory | 5. M2 Executive Function | 6. M3 Executive Function | 7. M2 BTACT Composite | 8. M3 BTACT Composite | 9. Aperiodic Exponent | 10.  IAPF |
| --- | --- | --- | --- | --- | --- | --- | --- | --- | --- | --- |
| 1. Age (MIDUS 2 Neuro) | - | -0.12  [-0.25, 0.01]  *p* = 0.064  *n* = 235 | **-0.18**  **[-0.30,-0.05]**  ***p* = 0.006**  *n* = 228 | **-0.24**  **[-0.36,-0.11]**  ***p* < 0.001**  *n* = 223 | **-0.35**  **[-0.46,-0.23]**  ***p* < 0.001**  *n* = 228 | **-0.32**  **[-0.43,-0.20]**  ***p* < 0.001**  *n* = 230 | **-0.36**  **[-0.47,-0.24]**  ***p* < 0.001**  *n* = 224 | **-0.34**  **[-0.45,-0.21]**  ***p* < 0.001**  *n* = 219 | **-0.25**  **[-0.37,-0.13]**  ***p* < 0.001**  *n* = 235 | **-0.24**  **[-0.35,-0.11]**  ***p* < 0.001**  *n* = 235 |
| 2. Total Lag (M2 to M3 Cog Projects) | -0.12  [-0.25, 0.01]  *p* = 0.064  *n* = 235 | - | -0.06  [-0.19, 0.07]  *p* = 0.345  *n* = 226 | -0.07  [-0.20, 0.06]  *p* = 0.314  *n* = 223 | **-0.17**  **[-0.29,-0.04]**  ***p* = 0.011**  *n* = 228 | **-0.30**  **[-0.41,-0.17]**  ***p* < 0.001**  *n* = 230 | **-0.16**  **[-0.29,-0.03]**  ***p* = 0.015**  *n* = 224 | **-0.29**  **[-0.41,-0.17**]  ***p* < 0.001**  *n* = 219 | **0.17**  **[0.04, 0.13]**  ***p* = 0.009**  *n* = 235 | -0.05  [-0.18, 0.07]  *p* = 0.405  *n* = 235 |
| 3. Episodic Memory MIDUS 2 | **-0.18**  **[-0.30,-0.05]**  ***p* = 0.006**  *n* = 228 | -0.06  [-0.19, 0.07]  *p* = 0.345  *n* = 226 | - | **0.56**  **[0.46, 0.64]**  ***p* < 0.001**  *n* = 217 | **0.36**  **[0.24, 0.47]**  ***p* < 0.001**  *n* = 224 | **0.32**  **[0.19, 0.43]**  ***p* < 0.001**  *n* = 223 | **0.56**  **[0.47, 0.65]**  ***p* < 0.001**  *n* = 222 | **0.41**  **[0.29, 0.52]**  ***p* < 0.001**  *n* = 213 | 0.00  [-0.13, 0.13]  *p* = 0.958  *n* = 228 | **0.15**  **[0.02, 0.28]**  ***p* = 0.020**  *n* = 228 |
| 4. Episodic Memory MIDUS 3 | **-0.24**  **[-0.36,-0.11]**  ***p* < 0.001**  *n* = 223 | -0.07  [-0.20, 0.06]  *p* = 0.314  *n* = 223 | **0.56**  **[0.46, 0.64]**  ***p* < 0.001**  *n* = 217 | - | **0.26**  **[0.13, 0.38]**  ***p* < 0.001**  *n* = 217 | **0.37**  **[0.25, 0.48]**  ***p* < 0.001**  *n* = 219 | **0.39**  **[0.27, 0.50]**  ***p* < 0.001**  *n* = 213 | **0.59**  **[0.49, 0.67]**  ***p* < 0.001**  *n* = 219 | -0.02  [-0.15, 0.12]  *p* = 0.809  *n* = 223 | **0.21**  **[0.08, 0.33]**  ***p* = 0.002**  *n* = 223 |
| 5. Executive Function MIDUS 2 | **-0.35**  **[-0.46,-0.23]**  ***p* < 0.001**  *n* = 228 | **-0.17**  **[-0.29,-0.04]**  ***p* = 0.011**  *n* = 228 | **0.36**  **[0.24, 0.47]**  ***p* < 0.001**  *n* = 224 | **0.26**  **[0.13, 0.38]**  ***p* < 0.001**  *n* = 217 | - | **0.69**  **[0.62, 0.76]**  ***p* < 0.001**  *n* = 223 | **0.97**  **[0.96, 0.98]**  ***p* < 0.001**  *n* = 224 | **0.68**  **[0.60, 0.74]**  ***p* < 0.001**  *n* = 213 | 0.11  [-0.02, 0.24]  *p* = 0.083  *n* = 228 | **0.16**  **[0.07, 0.29]**  ***p* = 0.013**  ***n* = 228** |
| 6. Executive Function MIDUS 3 | **-0.32**  **[-0.43,-0.20]**  ***p* < 0.001**  *n* = 230 | **-0.30**  **[-0.41,-0.17]**  ***p* < 0.001**  *n* = 230 | **0.32**  **[0.19, 0.43]**  ***p* < 0.001**  *n* = 223 | **0.37**  **[0.25, 0.48]**  ***p* < 0.001**  *n* = 219 | **0.69**  **[0.62, 0.76]**  ***p* < 0.001**  *n* = 223 | - | **0.69**  **[0.61, 0.75]**  ***p* < 0.001**  *n* = 219 | **0.97**  **[0.96, 0.98]**  ***p* < 0.001**  *n* = 219 | 0.02  [-0.11, 0.15]  *p* = 0.809  *n* = 230 | **0.19**  **[0.07, 0.32]**  ***p* = 0.003**  *n* = 230 |
| 7. BTACT Composite MIDUS 2 | **-0.36**  **[-0.47,-0.24]**  ***p* < 0.001**  *n* = 224 | **-0.16**  **[-0.29,-0.03]**  ***p* = 0.015**  *n* = 224 | **0.56**  **[0.47, 0.65]**  ***p* < 0.001**  *n* = 222 | **0.39**  **[0.27, 0.50]**  ***p* < 0.001**  *n* = 213 | **0.97**  **[0.96, 0.98]**  ***p* < 0.001**  *n* = 224 | **0.69**  **[0.61, 0.75]**  ***p* < 0.001**  *n* = 219 | - | **0.69**  **[0.61, 0.75]**  ***p* < 0.001**  *n* = 219 | 0.10  [-0.03, 0.23]  *p* = 0.119  *n* = 224 | **0.18**  **[0.05, 0.31]**  ***p* = 0.006**  *n* = 224 |
| 8. BTACT Composite MIDUS 3 | **-0.34**  **[-0.45,-0.21]**  ***p* < 0.001**  *n* = 219 | **-0.29**  **[-0.41,-0.17**]  ***p* < 0.001**  *n* = 219 | **0.41**  **[0.29, 0.52]**  ***p* < 0.001**  *n* = 213 | **0.59**  **[0.49, 0.67]**  ***p* < 0.001**  *n* = 219 | **0.68**  **[0.60, 0.74]**  ***p* < 0.001**  *n* = 213 | **0.97**  **[0.96, 0.98]**  ***p* < 0.001**  *n* = 219 | **0.69**  **[0.61, 0.75]**  ***p* < 0.001**  *n* = 219 | - | 0.01  [-0.12, 0.14]  *p* = 0.886  *n* = 219 | **0.23**  **[0.10, 0.35]**  ***p* < 0.001**  *n* = 219 |
| 9. Aperiodic Exponent | **-0.31**  **[-0.42,-0.19]**  ***p* < 0.001**  *n* = 235 | 0.10  [-0.03, 0.22]  *p* = 0.127  *n* = 235 | 0.05  [-0.08, 0.18]  *p* = 0.410  *n* = 228 | 0.03  [-0.10, 0.17]  *p* = 0.603  *n* = 223 | **0.15**  **[0.02, 0.27]**  ***p* = 0.025**  *n* = 228 | 0.07  [-0.06, 0.19]  *p* = 0.321  *n* = 230 | **0.15**  **[0.02, 0.28]**  ***p* = 0.025**  *n* = 224 | 0.07  [-0.06, 0.20]  *p* = 0.293  *n* = 219 | - | **-0.22**  **[-0.34,-0.10]**  ***p* < 0.001**  *n* = 235 |
| 10. IAPF | -0.13  [-0.26 -0.00]  *p* = 0.044  *n* = 235 | **-0.16**  **[-0.28,-0.03]**  ***p* = 0.017**  *n* = 235 | **0.21**  **[0.08, 0.33]**  ***p* = 0.002**  *n* = 228 | **0.17**  **[0.04, 0.29]**  ***p* = 0.013**  *n* = 223 | **0.15**  **[0.02, 0.28]**  ***p* = 0.022**  *n* = 228 | **0.19**  **[0.06, 0.31]**  ***p* = 0.004**  *n* = 230 | **0.20**  **[0.06, 0.33]**  ***p* = 0.002**  *n* = 224 | **0.20**  **[0.07, 0.32]**  ***p* = 0.004**  *n* = 219 | -0.09  [-0.21, 0.04]  *p* = 0.182  *n* = 235 | - |

Table S53: Correlations and descriptive statistics between combined EEG metrics, eyes open only EEG metrics, and eyes closed only EEG metrics.

|  | Mean (SD) | 2. Aperiodic Exponent Eyes Open | 3. Aperiodic Exponent Eyes Closed | 4. IAPF Combined | 5. IAPF Eyes Open | 6 IAPF Eyes Closed |
| --- | --- | --- | --- | --- | --- | --- |
| 1. Aperiodic Exponent Combined | 1.20 (0.28) | **0.96**  **[0.95, 0.97]**  ***p* < 0.001** | **0.97**  **[0.96, 0.97]**  ***p* < 0.001** | **-0.16**  **[-0.28,-0.03]**  ***p* = 0.023** | -0.10  [-0.23, 0.03]  *p* = 0.159 | **-0.16**  **[-0.28,-0.03]**  ***p* = 0.024** |
| 2. Aperiodic Exponent Eyes Open | 1.12 (0.28) | - | **0.87**  **[0.83, 0.90]**  ***p* < 0.001** | -0.09  [-0.21, 0.04]  *p* = 0.229 | -0.03  [-0.16, 0.10]  *p* = 0.661 | -0.09  [-0.21, 0.04]  *p* = 0.226 |
| 3. Aperiodic Exponent Eyes Closed | 1.27 (0.29) |  | - | **-0.23**  **[-0.35,-0.10]**  ***p* = 0.001** | **-0.16**  **[-0.28,-0.03]**  ***p* = 0.024** | **-0.22**  **[-0.34,-0.10]**  ***p* = 0.001** |
| 4. IAPF Combined | 9.60 (0.98) |  |  | **-** | **0.66**  **[0.58, 0.72]**  ***p* < 0.001** | **0.93**  **[0.91, 0.94]**  ***p* < 0.001** |
| 5. IAPF Eyes Open | 9.65 (1.00) |  |  |  | **-** | **0.58**  **[0.49, 0.66]**  ***p* < 0.001** |
| 6. IAPF Eyes Closed | 9.64 (1.16) |  |  |  |  | **-** |

| Table S54: Multilevel models examine the interaction between aperiodic exponent and individual peak alpha frequency in eyes closed data. | | | | | | | | | |
| --- | --- | --- | --- | --- | --- | --- | --- | --- | --- |
|  | **Episodic Memory** | | | **Executive Functioning** | | | **BTACT** | | |
| *Predictors* | *Estimates* | *CI* | *p* | *Estimates* | *CI* | *p* | *Estimates* | *CI* | *p* |
| Intercept (M2) | **-0.43** | **-0.61 – -0.26** | **<0.001** | -0.07 | -0.19 – 0.05 | 0.233 | **-0.13** | **-0.24 – -0.02** | **0.024** |
| MIDUS Wave | -0.11 | -0.24 – 0.02 | 0.105 | **-0.41** | **-0.49 – -0.34** | **<0.001** | **-0.36** | **-0.43 – -0.29** | **<0.001** |
| Exponent | -0.01 | -0.47 – 0.45 | 0.973 | 0.19 | -0.12 – 0.50 | 0.232 | 0.16 | -0.13 – 0.44 | 0.274 |
| Individual Alpha Peak Frequency | 0.05 | -0.08 – 0.18 | 0.411 | 0.04 | -0.05 – 0.13 | 0.360 | 0.04 | -0.04 – 0.12 | 0.285 |
| Age | **-0.02** | **-0.03 – -0.01** | **<0.001** | **-0.02** | **-0.03 – -0.01** | **<0.001** | **-0.02** | **-0.03 – -0.02** | **<0.001** |
| Sex | **0.72** | **0.51 – 0.93** | **<0.001** | 0.11 | -0.03 – 0.26 | 0.133 | **0.22** | **0.08 – 0.35** | **0.002** |
| Education | **0.17** | **0.05 – 0.30** | **0.007** | **0.14** | **0.06 – 0.22** | **0.001** | **0.14** | **0.06 – 0.21** | **<0.001** |
| Lag between Waves | -0.09 | -0.21 – 0.02 | 0.110 | **-0.21** | **-0.29 – -0.13** | **<0.001** | **-0.20** | **-0.27 – -0.12** | **<0.001** |
| Wave X Exponent | 0.18 | -0.29 – 0.65 | 0.457 | -0.12 | -0.39 – 0.14 | 0.369 | -0.03 | -0.28 – 0.23 | 0.841 |
| Wave X IPAF | 0.10 | -0.03 – 0.23 | 0.144 | 0.04 | -0.04 – 0.11 | 0.336 | 0.05 | -0.02 – 0.12 | 0.182 |
| Exponent X IPAF | -0.14 | -0.54 – 0.26 | 0.492 | -0.08 | -0.35 – 0.19 | 0.558 | -0.10 | -0.35 – 0.15 | 0.422 |
| Wave X Exponent X IPAF | 0.33 | -0.09 – 0.75 | 0.128 | 0.24 | -0.01 – 0.48 | 0.059 | **0.31** | **0.07 – 0.54** | **0.010** |
| **Random Effects** | | | | | | | | | |
| σ^2^ | 0.47 | | | 0.15 | | | 0.13 | | |
| τ_00_ | 0.38 _M2ID_ | | | 0.22 _M2ID_ | | | 0.19 _M2ID_ | | |
| ICC | 0.45 | | | 0.60 | | | 0.59 | | |
| N | 234 _M2ID_ | | | 235 _M2ID_ | | | 234 _M2ID_ | | |
| Observations | 451 | | | 458 | | | 443 | | |
| Marginal R^2^ / Conditional R^2^ | 0.205 / 0.563 | | | 0.302 / 0.718 | | | 0.325 / 0.724 | | |

Table S55: Examining the Wave X Aperiodic Exponent X Individual Peak Alpha Frequency interaction from analyses described in Table S54 (eyes closed only) through the slope of the change in cognitive function over waves for each BTACT measure at high and low levels of each EEG metric.

**Change in BTACT Episodic Memory Factor from M2 to M3**

|  | *Slope of IAPF* | | *95% CI* | | *p-value* | |
| --- | --- | --- | --- | --- | --- | --- |
| Low Exponent(-1 SD) | 0.00 | | -0.18 – 0.19 | | 0.986 | |
| High Exponent (+1 SD) | **0.19** | | **0.02 – 0.37** | | **0.031** | |
|  | *Slope of* Exponent | | *95% CI* | | *p-value* | |
| Low IAPF (-1 SD) | -0.15 | | -0.66 – 0.36 | | 0.573 | |
| High IAPF (+1 SD) | 0.50 | | -0.23 – 1.23 | | 0.176 | |
| **Change in BTACT Executive Functioning Factor from M2 to M3** | | | | | | |
|  | | *Slope of IAPF* | | *95% CI* | | *p-value* |
| Low Exponent (-1 SD) | | -0.03 | | -0.14 – 0.08 | | 0.554 |
| High Exponent (+1 SD) | | **0.11** | | **0.01 – 0.21** | | **0.038** |
|  | | *Slope of* Exponent | | *95% CI* | | *p-value* |
| Low IAPF (-1 SD) | | **-0.36** | | **-0.66– -0.05** | | **0.022** |
| High IAPF (+1 SD) | | 0.11 | | -0.30 – 0.53 | | 0.586 |
| **Change in BTACT Overall Composite from M2 to M3** | | | | | | |
|  | | *Slope of IAPF* | | *95% CI* | | *p-value* |
| Low Exponent (-1 SD) | | -0.04 | | -0.14 – 0.06 | | 0.431 |
| High Exponent (+1 SD) | | **0.14** | | **0.04 – 0.23** | | **0.004** |
|  | | *Slope of* Exponent | | *95% CI* | | *p-value* |
| Low IAPF (-1 SD) | | **-0.33** | | **-0.62 – -0.04** | | **0.024** |
| High IAPF (+1 SD) | | 0.28 | | -0.11 – 0.67 | | 0.164 |

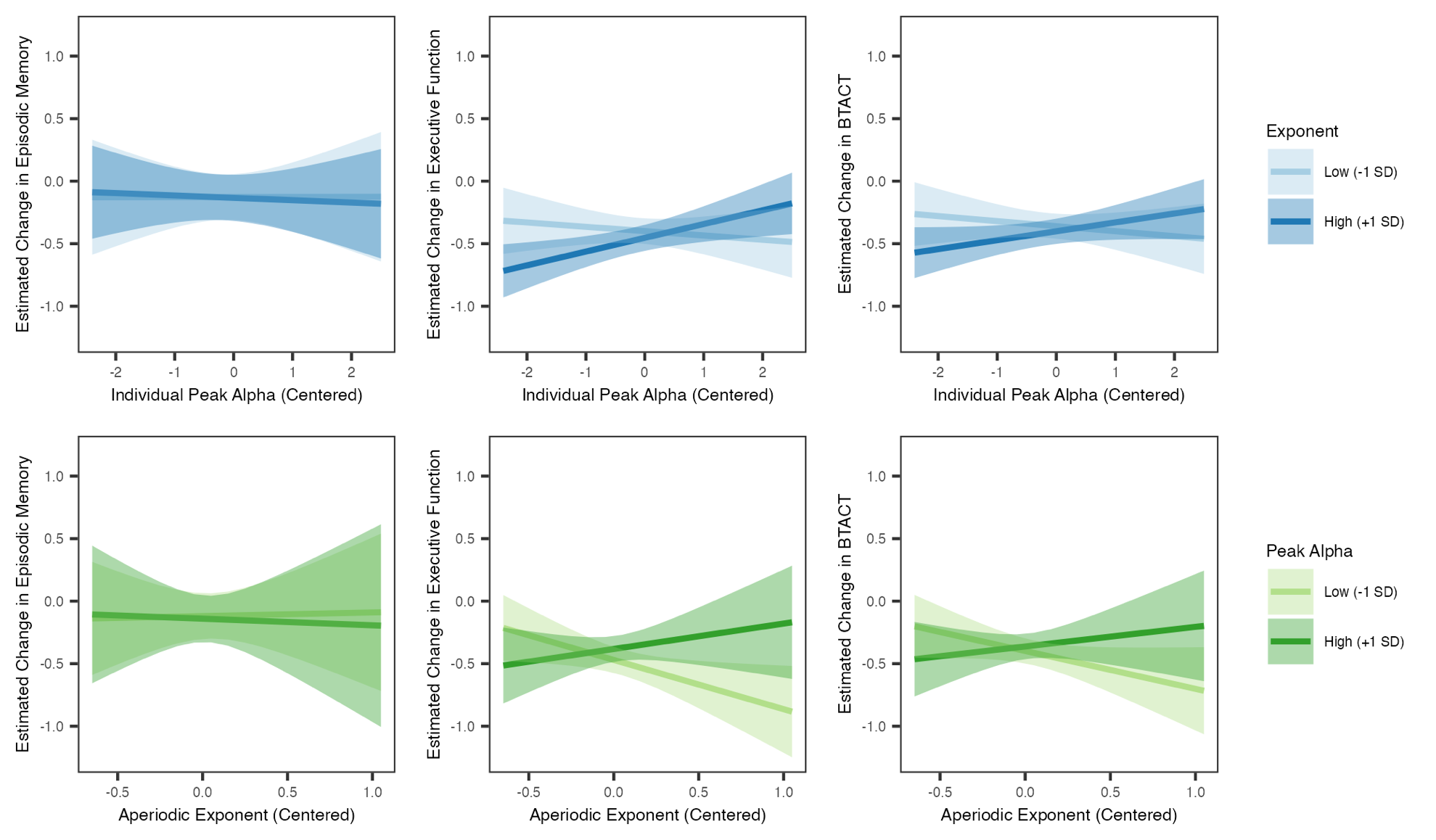

Figure S10. Wave by aperiodic exponent by individual peak alpha frequency interaction plot for models described in Table S54 (eyes closed only).

| Table S56: Multilevel models examine the interaction between aperiodic exponent and individual peak alpha frequency in eyes open data. | | | | | | | | | |
| --- | --- | --- | --- | --- | --- | --- | --- | --- | --- |
|  | **Episodic Memory** | | | **Executive Functioning** | | | **BTACT** | | |
| *Predictors* | *Estimates* | *CI* | *p* | *Estimates* | *CI* | *p* | *Estimates* | *CI* | *p* |
| Intercept (M2) | **-0.41** | **-0.59 – -0.24** | **<0.001** | -0.06 | -0.18 – 0.06 | 0.309 | **-0.12** | **-0.23 – -0.01** | **0.040** |
| Wave | **-0.13** | **-0.26 – -0.00** | **0.048** | **-0.43** | **-0.50 – -0.35** | **<0.001** | **-0.38** | **-0.45 – -0.31** | **<0.001** |
| Exponent | 0.09 | -0.36 – 0.55 | 0.692 | 0.13 | -0.18 – 0.43 | 0.412 | 0.14 | -0.15 – 0.42 | 0.343 |
| Individual Alpha Peak Frequency | 0.10 | -0.01 – 0.21 | 0.072 | 0.01 | -0.06 – 0.09 | 0.699 | 0.03 | -0.03 – 0.10 | 0.310 |
| Age | **-0.02** | **-0.03 – -0.01** | **<0.001** | **-0.02** | **-0.03 – -0.02** | **<0.001** | **-0.02** | **-0.03 – -0.02** | **<0.001** |
| Sex | **0.71** | **0.50 – 0.92** | **<0.001** | 0.10 | -0.05 – 0.25 | 0.175 | 0.21 | 0.07 – 0.34 | **0.003** |
| Education | **0.16** | **0.04 – 0.29** | **0.012** | **0.14** | **0.05 – 0.22** | **0.001** | **0.14** | **0.06 – 0.21** | **0.001** |
| Lag between Waves | -0.08 | -0.19 – 0.04 | 0.192 | **-0.20** | **-0.28 – -0.12** | **<0.001** | **-0.19** | **-0.26 – -0.11** | **<0.001** |
| Wave X Exponent | -0.01 | -0.48 – 0.46 | 0.961 | -0.10 | -0.36 – 0.17 | 0.478 | -0.07 | -0.33 – 0.18 | 0.576 |
| Wave X Alpha Freq | -0.01 | -0.12 – 0.11 | 0.876 | 0.04 | -0.03 – 0.10 | 0.245 | 0.02 | -0.05 – 0.08 | 0.626 |
| Exponent X Alpha Freq | 0.10 | -0.31 – 0.52 | 0.631 | -0.02 | -0.30 – 0.26 | 0.872 | 0.01 | -0.25 – 0.27 | 0.956 |
| Wave X Exponent X Alpha Freq | -0.04 | -0.47 – 0.40 | 0.875 | **0.26** | **0.01 – 0.51** | **0.043** | 0.20 | -0.04 – 0.44 | 0.106 |
| **Random Effects** | | | | | | | | | |
| σ^2^ | 0.47 | | | 0.15 | | | 0.14 | | |
| τ_00_ | 0.37 _M2ID_ | | | 0.23 _M2ID_ | | | 0.19 _M2ID_ | | |
| ICC | 0.44 | | | 0.60 | | | 0.58 | | |
| N | 234 _M2ID_ | | | 235 _M2ID_ | | | 234 _M2ID_ | | |
| Observations | 451 | | | 458 | | | 443 | | |
| Marginal R^2^ / Conditional R^2^ | 0.203 / 0.554 | | | 0.301 / 0.718 | | | 0.319 / 0.714 | | |

Table S57: Examining the Wave X Aperiodic Exponent X Individual Peak Alpha Frequency interaction from analyses described in Table S56 (eyes open only) through the slope of the change in cognitive function over waves for each BTACT measure at high and low levels of each EEG metric.

**Change in BTACT Episodic Memory Factor from M2 to M3**

|  | *Slope of IAPF* | | *95% CI* | | *p-value* | |
| --- | --- | --- | --- | --- | --- | --- |
| Low Exponent (-1 SD) | 0.00 | | -0.18 – 0.19 | | 0.994 | |
| High Exponent (+1 SD) | -0.02 | | -0.17 – 0.13 | | 0.802 | |
|  | *Slope of Exponent* | | *95% CI* | | *p-value* | |
| Low IAPF (-1 SD) | 0.03 | | -0.57 – 0.63 | | 0.925 | |
| High IAPF (+1 SD) | -0.05 | | -0.82 – 0.72 | | 0.894 | |
| **Change in BTACT Executive Functioning Factor from M2 to M3** | | | | | | |
|  | | *Slope of IAPF* | | *95% CI* | | *p-value* |
| Low Exponent (-1 SD) | | -0.03 | | -0.14 – 0.07 | | 0.517 |
| High Exponent (+1 SD) | | **0.11** | | **0.03 – 0.19** | | **0.010** |
|  | | *Slope of Exponent* | | *95% CI* | | *p-value* |
| Low IAPF (-1 SD) | | **-0.39** | | **-0.74– -0.04** | | **0.027** |
| High IAPF (+1 SD) | | 0.20 | | -0.23 – 0.63 | | 0.352 |
| **Change in BTACT Overall Composite from M2 to M3** | | | | | | |
|  | | *Slope of IAPF* | | *95% CI* | | *p-value* |
| Low Exponent (-1 SD) | | -0.04 | | -0.14 – 0.06 | | 0.437 |
| High Exponent (+1 SD) | | 0.07 | | -0.01 – 0.15 | | 0.084 |
|  | | *Slope of Exponent* | | *95% CI* | | *p-value* |
| Low IAPF (-1 SD) | | -0.30 | | -0.64 – 0.03 | | 0.075 |
| High IAPF (+1 SD) | | 0.16 | | -0.26 – 0.58 | | 0.463 |

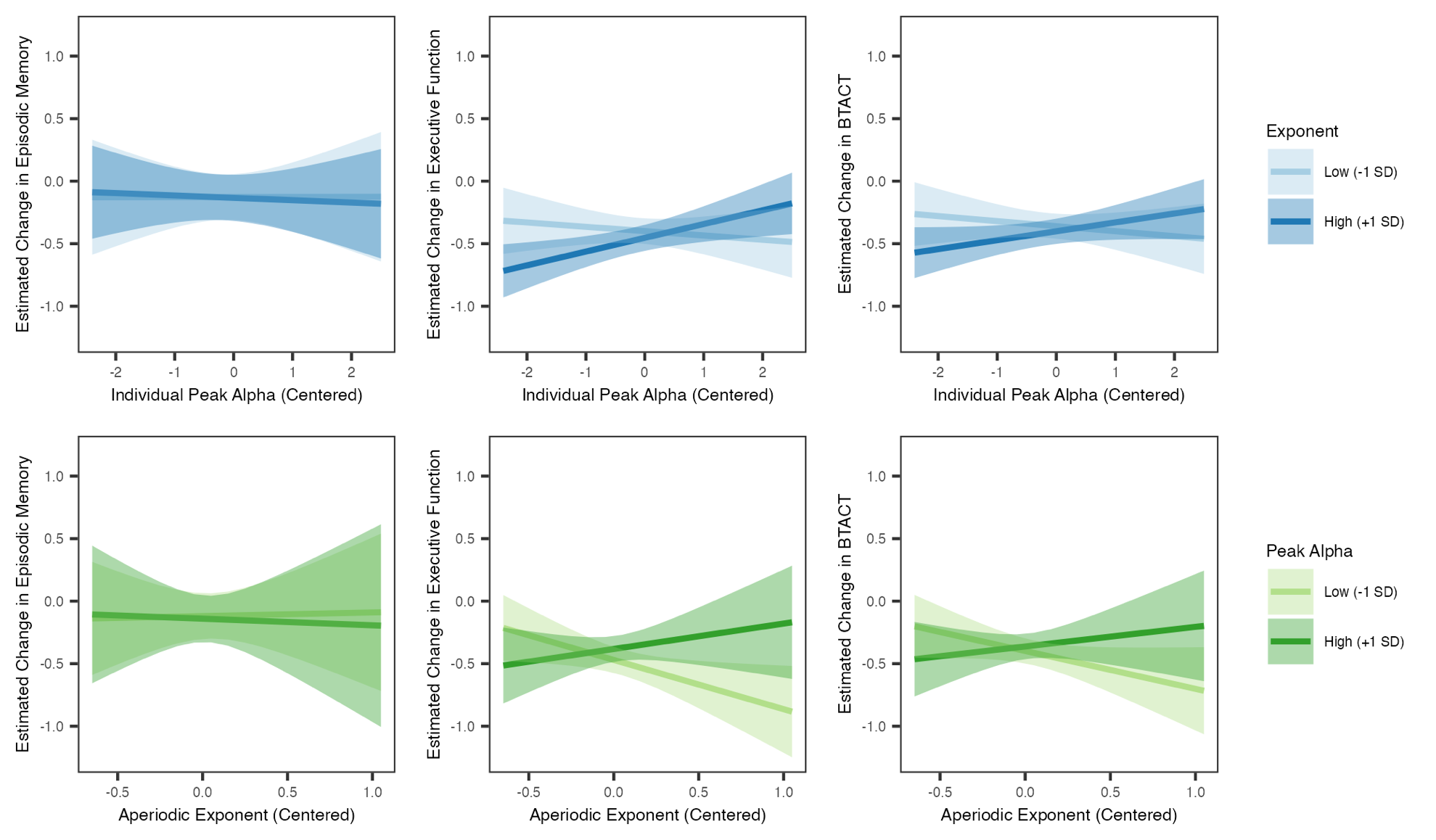

Figure S11. Wave by aperiodic exponent by individual peak alpha frequency interaction plot for models described in Table S56 (eyes open only).

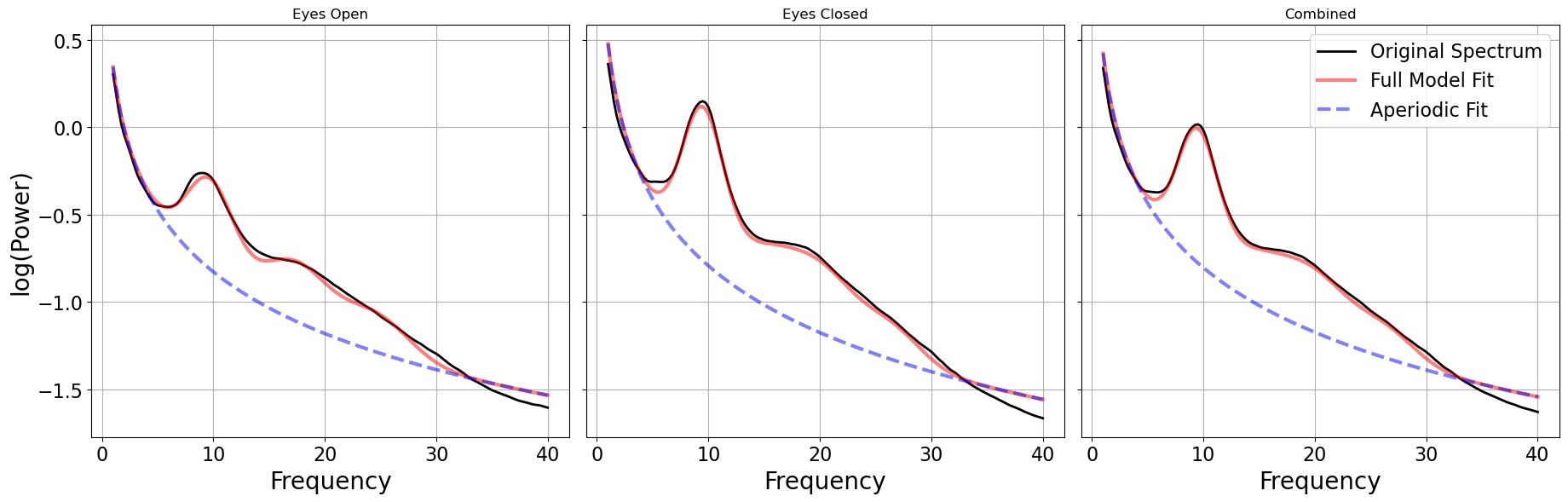
Figure S12. Grand average spectra and fitted models for eyes open only, eyes closed only, and combined eyes open and eyes closed resting EEG recordings.
